# Miniaturized Workflow for Transcriptomic Profiling of Urinary Extracellular RNA during Pregnancy

**DOI:** 10.1101/2025.03.03.639539

**Authors:** Srimeenakshi Srinivasan, Peter De Hoff, Amber L. Morey, Aishwarya Vuppala, Marina Mochizuki, Robert E. Morey, Morgan Meads, Erika Duggan, Derek E. Wildman, John P. Nolan, Priyadarshini Pantham

## Abstract

Urine contains extracellular RNA (exRNA) carried by extracellular vesicles (EVs) and other biomolecular complexes. There is currently a need for studies focused on female cohorts to develop new methods for non-invasive analysis of biofluids to create reference profiles and for identification of biomarkers of reproductive and pregnancy disorders. The objective of this study was therefore to identify optimal methods for transcriptomic profiling of urine by testing different exRNA isolation and scalable library preparation methods that enable detection of biomarkers that reflect pregnancy-associated changes in the placenta and maternal tissues. RNA was extracted from pooled and individual urine samples obtained from normal non-pregnant and pregnant females, as well as males, using input volumes of either 0.6 mL, 1 mL, or 4 mL. Samples were extracted using methods that focused either on isolating vesicular (EV-associated) or total (EV-associated and non EV-associated) exRNA. Small RNA libraries (n=208) were prepared using the NEBNext Small RNA Library Prep kit and long RNA libraries (n=97) were prepared using the SMART-Seq v4 Ultra Low Input RNA or the SMARTer Stranded Total RNA-Seq Kit v2 Pico Input kits (Takara). Principal component analysis showed that the greatest source of variance amongst technical replicates of small RNA libraries (n=176 which passed quality control) was exRNA isolation method, and amongst long RNA libraries (n=97 which passed quality control) was library preparation method. Long RNA libraries prepared from exRNA extracted using miRCURY showed that the SMART-Seq v4 method yielded significantly more uniquely mapped reads compared to the Pico v2 method (p<0.05). We have established a scalable pipeline for small and long RNA-Seq profiling of exRNA in urine in a reproducible manner, which we used to identify differentially expressed urinary exRNAs in pregnancy, and will enable transcriptomic profiling of urinary exRNA in disorders of pregnancy, including preeclampsia.

## 1. INTRODUCTION

Extracellular vesicles (EVs) are lipid bilayer-bound structures that are released from all cell types and play a role in intra-and intercellular communication (1). They vary in size, biogenesis, molecular cargo, and function in normal and pathological conditions. Extracellular RNA (exRNA), including EV-and non-EV-associated RNA found in biofluids may reflect the function of their cell/tissue of origin and are a novel source of biomarkers of disease pathologies (1). Some methods of exRNA extraction yield EV-enriched RNAs by first isolating EVs and then performing RNA extraction (vesicular exRNA), while others co-isolate EV-associated and non-EV-associated exRNA (total exRNA) (2). Extracellular micro-RNA (miRNA) and mRNA biomarkers in urine have been identified in disorders affecting renal function, particularly bladder cancer (3, 4), diabetic nephropathy (5), and prostate cancer (6). Urine contains exRNA that can be derived from various tissue types, including the kidney (7), heart (8), liver, and potentially the placenta during pregnancy. Urine “liquid biopsies” are therefore a potential source of exRNA biomarkers that reflect the functions of a variety of tissues, particularly the kidney.

The placenta is an organ of primarily fetal origin that functions as the means of nutrient and oxygen transport and waste removal between the mother and fetus during pregnancy. The human placenta is ‘hemochorial’ in nature, with its outermost layer in direct contact with maternal blood starting from about the 13^th^ week of gestation (9). The placenta also releases EVs and other exRNA carriers directly into the maternal circulation throughout pregnancy (10). Placenta-derived EVs can localize to various maternal organs, including the kidneys (11), supporting the use of urine as a potential source of RNA biomarkers of placental origin. Given the ease of obtaining maternal urine samples throughout pregnancy, optimizing methods for urinary exRNA isolation and transcriptomic profiling opens novel avenues to identify biomarkers and pathophysiological mechanisms in pregnancy. The maternal kidney undergoes marked changes during normal pregnancy, including an increase in size and length. Maternal plasma volume, glomerular filtration rate, and renal plasma flow also increase (12, 13). Physiological changes during pregnancy may therefore be reflected in urine exRNA. Preeclampsia, a leading cause of maternal mortality and morbidity worldwide, is a hypertensive disorder of pregnancy diagnosed by the onset of hypertension and proteinuria after 20 weeks of gestation (14). There is a critical need to identify biomarkers of preeclampsia to allow for earlier identification and better management of women affected by this syndrome. Preeclampsia causes alterations in maternal kidney function, including glomerular capillary endotheliosis leading to disruption of the glomerular filtration barrier and proteinuria (12). Since the kidney is involved in the pathophysiology of preeclampsia, urine presents a novel source of exRNA biomarkers for diagnosis of this condition as well as other conditions during pregnancy in which the maternal kidneys are affected. Here we have comprehensively profiled urinary exRNA during pregnancy.

Per the position paper by the Urine Task Force of the Rigor and Standardization Subcommittee of ISEV, there remains a great need for “further optimization and standardization to foster scientific advances in uEV research and subsequent successful translation into clinical practice” (15). Standardization of total and vesicular exRNA isolation methods to generate robust and reproducible transcriptomic profiling data for biomarker discovery in urine remains a significant challenge (15). It is essential to test various methods of total and vesicular exRNA isolation methods to measure consistency between replicate reference samples prior to testing samples for clinical diagnosis. Further, the cost of library preparation for next generation sequencing (NGS) remains a significant factor in planning large scale, high-throughput studies. In this study, we have tested five different commercially available methods for isolation of total (Norgen, miRNeasy advanced) or vesicular (miRCURY, ExoRNeasy, SeraMir) urinary exRNA to generate RNA-Seq libraries and transcriptomic profiles from male and female non-pregnant and pregnant urine reference pools and samples from individuals using Illumina NGS platforms. Urinary exRNA was isolated using either PEG-based buffer (miRCURY and SeraMir) or membrane affinity (ExoRNeasy) to precipitate uEVs followed by RNA extraction with the miRNeasy mini kit for “vesicular” exRNA isolation, or direct lysis of neat urine followed by spin column chromatography for “total” exRNA isolation (Norgen and miRNeasy advanced).

We have also quantified the number and size of uEVs in urinary reference samples using three orthogonal methods: microfluidic resistive pulse sensing (MRPS), nanoparticle tracking analysis (NTA), and high-resolution vesicle flow cytometry (vFC). These methods are based on different biophysical principles, and thus vary in their limits of detection and ability to quantify EVs and detect surface markers. Using vFC, we determined staining characteristics for antibodies to tetraspanins CD9, CD63, and CD81, surface markers of exosomes (20-150nm), and annexin V, a surface marker of exosomes and microvesicles (100-1000nm) (16) in pregnant and non-pregnant urine pools, as well as urine pools enriched for uEVs using a PEG-based enrichment method. These analyses are essential to contribute to our understanding of the analysis of cell-free urine, and the development of clinical applications of uEVs in future (15).

We present a miniaturized high-throughput workflow to generate reproducible urinary exRNA-Seq data, allowing for the analysis of up to 384 samples in a single batch. We utilized one method for small RNA library preparation and two methods for long RNA library preparation of urinary exRNA on a miniaturized scale, reducing reaction volumes to 1/5^th^ of the volumes in the manufacturers’ protocols by using automated small volume liquid handlers (Mosquito HV/HTS/X1, SPT Labtech, Royston, UK). The use of Mosquito robotic liquid handlers (HV and X1 single-channel ‘hit-picker’: 500nL-5μL; HTS/LV: 25nL-1.2μL) significantly reduce the per-sample cost of library preparation and enables large-scale screening of samples for clinical studies. Miniaturized workflows using these instruments have been established for wastewater genomic surveillance of SARS-CoV2 variant transmission (17) and identification of exRNA biomarkers in the serum of women with preeclampsia (18, 19). Using this miniaturized workflow for urinary exRNA, we have identified RNAs that are differentially expressed and enriched in maternal urine during pregnancy compared to the non-pregnant state. We have generated reference transcriptomic profiles for maternal urine pools during pregnancy which can be utilized in future studies investigating the urinary transcriptomic profile of pregnancy disorders such as preeclampsia.

## 2. MATERIALS & METHODS

### 2.1. Urine Sample Collection

Urine was collected from healthy adult donors who provided written informed consent under IRB protocols approved by the Human Research Protections program at the University of California, San Diego (UCSD). Urine samples were collected by spontaneous voiding, at any time of day into sterile containers manufactured from polypropylene and polyethylene, free from latex and bovine contamination (Parter Medical Products^TM^ sterile specimen containers, Fisher Scientific, cat. no. 243517). Samples were centrifuged at 2000 X *g* for 10 minutes at room temperature to pellet cells and cellular debris, supernatant was aliquoted, and stored at-80°C within 2 hours of collection until further processing. Upon thawing, samples were centrifuged at 2000 X *g* for 10 minutes at 4°C to remove any debris following the freeze-thaw cycle, and supernatant was collected for processing. Equal volumes of cell-free urine from ten healthy individuals (Supplementary Table 1) per group were pooled to create three separate urine pools: male (M pool); non-pregnant female (NP pool); and pregnant female (P pool). To create the pregnant urine pool, samples were collected from normal pregnancies with an average gestational age (GA) of 25 weeks (2^nd^ trimester). To compare the exRNA profile of pregnant and non-pregnant subjects, samples were collected from healthy non-pregnant females (NP, n=5) and pregnant individuals with an average GA of 34 weeks (3^rd^ trimester) (P, n=5) (Supplementary Table 1). Samples were extracted as described below, and the number of samples extracted using each RNA isolation method is summarized in Table 1.

**Table 1:**
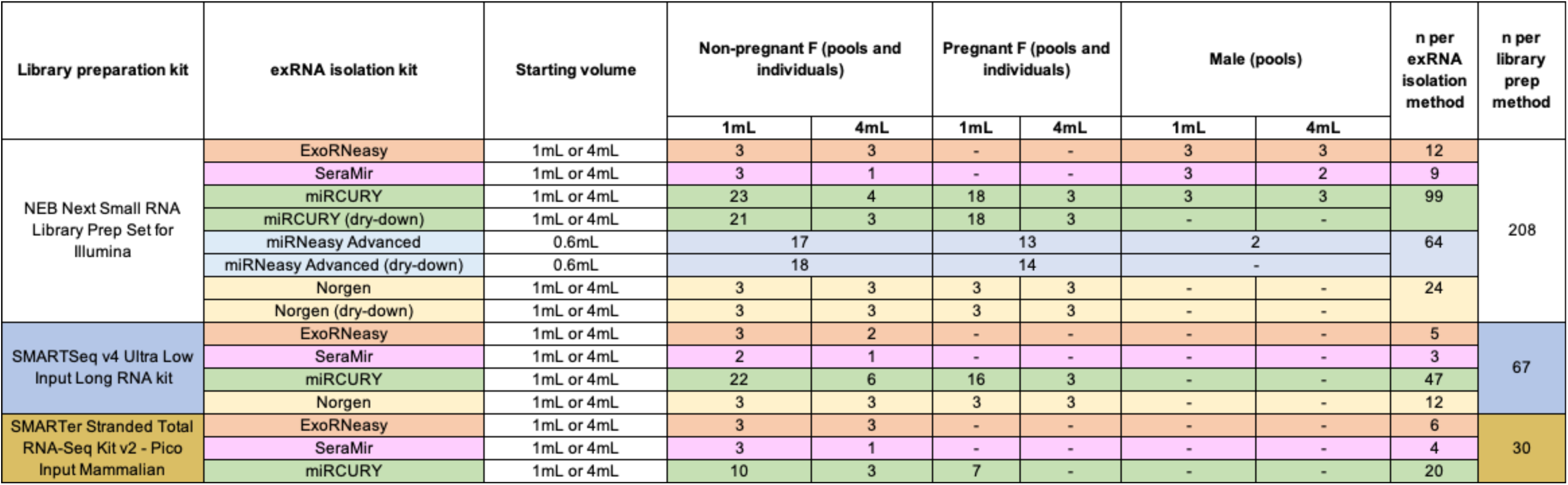
Summary of exRNA isolation and library preparation methods along with number and type of samples for which RNA-Seq libraries were generated in this study.

### 2.2. Kit-based Urinary Extracellular RNA Isolation Methods

Commercially available exRNA isolation kits were used to extract total or vesicular RNA from urine following the manufacturers’ protocols unless otherwise indicated. The different methods, sample types (pooled urine samples: male, female non-pregnant, female pregnant; individual urine samples: female non-pregnant and pregnant), and number of samples prepared using each method are summarized in detail Table 1 and Figure 1A. Urine samples were thawed on ice prior to extraction. ‘Vesicular’ methods involve an initial step to enrich for uEVs, followed by exRNA isolation. ‘Total’ methods involve a lysis step followed by phenol-free exRNA isolation. Total and vesicular exRNA was extracted from urine pools or individual samples using the methods described below. RNA extracted using each method was checked for quality in a subset of samples using the Agilent 6000 RNA Pico chip and Bioanalyzer (data not shown).

**Figure 1:**
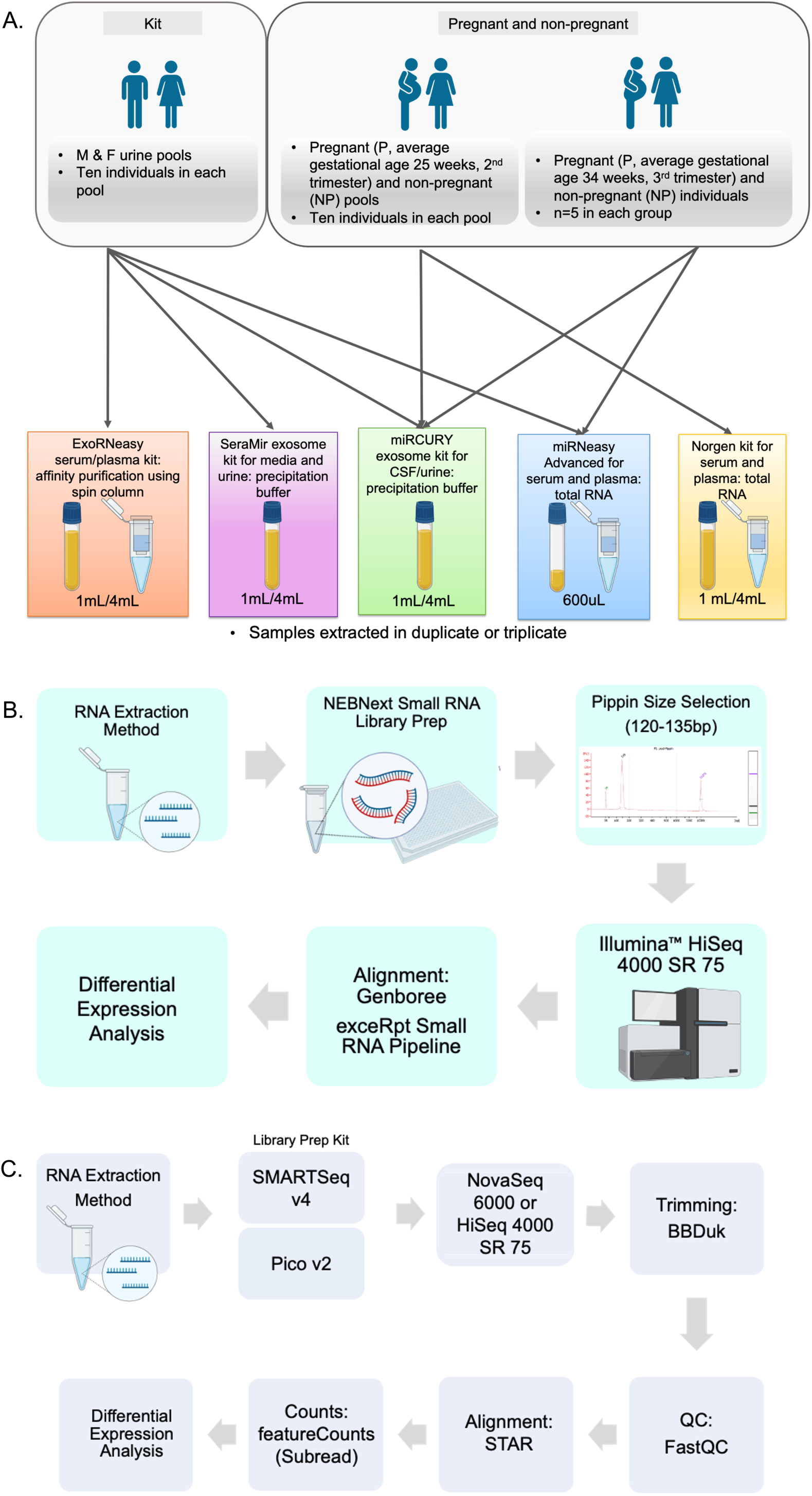
Schematic showing methods of urinary exRNA isolation and pipeline for small and long RNA-Seq. **(A)** Different populations and sample types used in the study: Male or female (non-pregnant and pregnant) urine pools were prepared by combining equal volumes of urine from n=10 individuals in each pool. The average gestational age (GA) of women in the pregnant urine pool was 25 weeks. For studies investigating pregnant and non-pregnant individuals, urine was collected from n=5 individuals. The average GA of pregnant individuals was 34 weeks. Samples were extracted using the following kits: For purification of uEVs followed by RNA isolation, ExoRNeasy (affinity purification using a spin column), SeraMir (precipitation buffer), and miRCURY (precipitation buffer) were used. For purification of total urinary extracellular RNA, the miRNeasy Advanced (mAdv) and Norgen kits were used. **(B)** Pipeline for small RNA library prep, RNA-Seq, and data analysis. **(C)** Pipeline for long RNA library prep using either the SMARTSeq v4 Ultra Low Input or Pico v2 Mammalian Kits (TakaraBio), RNA-Seq, and data analysis.

#### 2.2.1. Vesicular exRNA Isolation Methods from Urine

The *ExoRNeasy Serum/Plasma Midi Kit* (“ExoRNeasy”, Qiagen, cat. no: 77044) was used to extract RNA from 1 mL or 4 mL starting volumes of urine pools in triplicate. Urine EVs were isolated using membrane-based affinity spin columns, followed by extraction of RNA using Qiazol and the miRNeasy Micro Kit spin columns using the manufacturer’s protocol (Qiagen, cat. no. 217084). RNA was eluted with addition of 20μL of RNAse-free water and centrifugation at 100 x *g* for 1 minute, followed by a second elution using the eluate from the first step and full speed centrifugation for 1 minute.

The *SeraMir Exosome RNA Amplification Kit* (“SeraMir”, System Biosciences, cat. no: RA806TC-1) was used to extract 1 mL or 4 mL starting volumes of urine pools in triplicate (1 mL) or single samples (4 mL). Urine EVs were precipitated by incubating with the ExoQuick-TC at a 1:5 ratio overnight at 4°C, lysed using a phenol-free lysis buffer and RNA was bound to small RNA binding columns. RNA was eluted with addition of 20 μL of RNAse-free water and centrifugation at 100 x *g* for 1 minute, followed by a second elution using the eluate from the first step and full speed centrifugation for 1 minute.

The *miRCURY Exosome Cell/Urine/CSF Kit* (“miRCURY”, Qiagen, cat. no. 76743) was used to extract 1 mL or 4 mL starting volumes of urine pools or individual urine samples in triplicate. EVs were precipitated from 1 mL or 4 mL of urine by incubating with 400 μL or 1.6 mL of Precipitation Buffer B respectively overnight at 4°C. Samples were centrifuged at 10,000 x *g* (for 4 mL urine) or 3200 x *g* (for 1 mL urine) for 30 minutes at 20°C, supernatant was discarded, and the remaining pellets were processed using Qiazol and the miRNeasy Micro Kit spin columns using the manufacturer’s protocol (Qiagen, cat. no. 217084). Samples were eluted either with the same 20 μL of RNAse-free water twice, or with 12 μL followed by 2 μL water, with a slow 1-minute spin at 100 x *g* followed by a 1-minute spin at full speed.

#### 2.2.2. Total exRNA Isolation Methods from Urine

The *miRNeasy Serum/Plasma Advanced Kit* (“mAdv”, Qiagen, cat. no. 217204) was used to extract 0.6 mL starting volumes of urine pools or individuals in triplicate. Samples were incubated with 60 µL of RPL buffer followed by 20 µL of RPP buffer and centrifuged for 12000 x *g* for 3 minutes. Supernatant was transferred to the RNeasy UCP MinElute spin column, 1 volume of isopropanol was added, samples were washed using buffers RWT and RPE, and 500 µL of 80% ethanol was added. Samples were eluted using the same 20 μL of RNAse-free water twice. Samples were then cleaned using Zymo RNA Clean and Concentrator (Zymo Research).

The *Plasma/Serum Circulating and Exosomal RNA Purification Mini Kit* (Slurry Format) (“Norgen”, Norgen Biotek, cat. no: 51000) was used to extract 1 mL and 4 mL starting volumes of urine pools or individual urine samples in triplicate. Samples were incubated with either 1.9 mL (for 1 mL urine) or 7.9 mL (for 4 mL urine) warmed Lysis Buffer A containing 2-Mercaptoethanol and 100μL slurry for 10 minutes at 60°C. Samples were washed twice with either 3 mL (for 1 mL urine) or 12 mL (for 4 mL urine) of 100% Ethanol, vortexed and centrifuged for 2 minutes at 1000rpm. Supernatant was decanted, 300 μL of Lysis Buffer A was added and incubated for 10 minutes at 60°C. 300 μL 100% Ethanol was added to the samples, loaded onto the spin columns, and centrifuged at 16000 x *g* for 1 minute. Samples were washed thrice with 400 μL Wash Solution A and centrifuged at 16000 x *g* for 1 minute, and once more for 3 minutes to dry the membrane. RNA was eluted by adding 15 μL of RNAse-free water followed by a slow spin for 2 minutes at 300 x *g*, and then 5 μL water was added followed by a second spin at 16000 x *g* for 3 minutes.

### 2.3. Miniaturized Library Preparation

#### 2.3.1. Small RNA Library Preparation and Sequencing

Small RNA libraries were prepared for sequencing using the NEB Next Small RNA Library Prep Set for Illumina (New England Biosciences, cat. no. E7330L) using the manufacturer’s protocols with the following modifications. All reactions were conducted at 1/5^th^ of the volume recommended in the manufacturer’s protocols, adapters were diluted 1:6 of the supplied concentration, and 18 PCR cycles were conducted. For the miRNeasy and miRCURY samples extracted from pools and individuals, either 1.2 μL RNA starting volume was used as input, or 4 μL of RNA was dried down using a speedvac without heat and resuspended in 1.2 μL water was used as input (dry-down samples, DD). Small RNA libraries were cleaned using the Zymo DNA Clean and Concentrate kit (Zymo research, cat. no. D4103). Libraries were pooled and size-selected using the Pippin Prep HT with a 125-150 bp size window to remove adapter dimers. Size-selected libraries were sequenced on the Illumina HiSeq 4000 as 75 bp single-end reads by the Institute for Genomic Medicine (IGM) at UCSD (Figure 1B) in four sequencing batches.

#### 2.3.2. Long RNA Library Preparation and Sequencing

The *SMART-Seq® v4 Ultra^TM^ Low Input RNA Kit* (“SMARTSeq v4”, TakaraBio Inc., cat.no. 634891) was used to prepare long RNA libraries for RNA-Seq with the following modifications. All reactions were conducted at 1/5^th^ of the volume recommended in the manufacturer’s protocol, with 1.9 μL of RNA starting volume as input, and 3’ SMART-Seq CDS Primer II A diluted 1:1 from the supplied concentration. cDNA amplification was conducted using 18 PCR cycles and amplified cDNA was purified using the Agencourt AMPure XP Kit (Fisher Scientific, cat. no. A63881). cDNA concentration was measured using Qubit, normalized to 0.8 ng/μL, and libraries were prepared using the Nextera XT DNA Library Preparation and Index Kits (Illumina, cat. no. FC-131-1096) with 5 minutes of tagmentation. cDNA libraries underwent purification and size-selection using Agencourt AMPure XP beads added at 0.6X the sample volume. Size-selected libraries were sequenced either on the Illumina HiSeq 4000 or the NovaSeq 6000 as 75 bp single-end reads by the IGM Center at UCSD in two sequencing batches (Figure 1C).

The *SMARTer Stranded Total RNA-Seq v2 – Pico Input Mammalian Kit* (“Pico v2”, TakaraBio Inc., cat.no. 634412) was also used to prepare long RNA libraries for RNA-Seq with the following modifications. All reactions were conducted at 1/5^th^ of the volume recommended in the manufacturer’s protocol, with 1.6 μL of RNA starting volume as input, and 0.2 μL of SMART Pico Oligos Mix v2. RNA was fragmented at 94°C for 2 min, cDNA was synthesized, and 5-cycle indexing PCR was conducted. Ribosomal RNA was depleted using ZapR v2, final cDNA amplification was conducted using 16 PCR cycles, and cDNA libraries underwent purification and size-selection using Agencourt AMPure XP beads added at 1X the sample volume. Size-selected libraries were sequenced on the Illumina HiSeq 4000 as 75 bp single-end reads by the IGM Center at UCSD in a single sequencing batch (Figure 1C).

### 2.4. RNA-Seq Data Analysis

#### 2.4.1. Analysis Pipeline and Data Quality

Small RNA-Seq data were processed using the Genboree exceRpt Small RNA-Seq Pipeline (20). Briefly, FASTQ files underwent adaptor trimming and mapping to endogenous human genome (hg38) using STAR v2.4.2a, with a minimum insert length set to 15 nucleotides and no mismatches permitted on the Genboree workbench. Long RNA-Seq data were processed by trimming reads of Illumina adapters using bbduk (21), trimmed FASTQ files were then aligned to the human genome (GRCh38) with STAR v2.7.3a and underwent quality control analysis using FastQC and MultiQC. Reads aligned to exons were counted using featureCounts in the Subread v2.0 package (22).

Complexity, defined as the total number of individual miRNAs or protein-coding mRNAs with raw counts ≥5 was calculated for each sample library. Analysis of quality control metrics, such as differences in % miRNA, % tRNA, and complexity, by exRNA isolation method was conducted using GraphPad Prism (version 10.2.3) with the Kruskal-Wallis test or the Mann Whitney U test for pairwise comparisons, with a p-value < 0.05 considered significant. For each miRNA/exRNA isolation method combination, the mean, range, interquartile range of expression (in CPM), expression window (1-10CPM, 10-100CPM, 100-1000CPM, 10000-100000CPM, 100000-100000 CPM, 100000-1000000 CPM), standard deviation and %CV (coefficient of variation) were calculated to determine the robustness and reproducibility of each exRNA isolation method.

Filtering for visualization and differential expression analysis was performed by retaining miRNAs and mRNAs present in at least 50% of all samples within each kit or group by calculating the value for 5 raw counts in cpm (counts per million) in the lowest complexity sample within each analysis ((5 ÷ total miRNA or mRNA read counts of lowest complexity sample) x 10^6^). The cutoff for filtering is summarized in Supplementary Table 3 for each analysis. Principal Component Analysis (PCA) to visualize the origin of variance amongst samples was conducted using the Qlucore data analysis and visualization software package.

#### 2.4.2 Differential Expression Analysis

Differentially expressed (DE) miRNAs and mRNAs were identified using the edgeR package in R v3.6.3, with normalization of filtered read counts using the TMM (trimmed mean of M component) method followed by the Exact test for miRNA counts, and TMM normalization followed by Limma-Voom and the Exact test for mRNA counts (23). Reported p-values were corrected for multiple hypothesis testing using the Benjamini-Hochberg method (FDR < 0.05). Differential expression analysis of miRNA data was focused on samples that were not vacuum concentrated. The number of samples used in each analysis for visualization or differential expression, total miRNA or mRNA read count cutoffs, and number of differentially expressed RNAs in each analysis is provided in Supplementary Table 3. Data are expressed in log2 cpm, and heatmaps were created using the Qlucore data analysis and visualization software package, with blue indicating lower expression and yellow indicating higher expression of miRNAs or mRNAs. Analysis of cellular component and molecular pathways of interest was conducted using lists of mRNAs that were enriched (significantly increased) in pregnant pools or individuals following differential expression analysis of non-pregnant vs. 2^nd^ trimester pregnant pools or non-pregnant vs. 3^rd^ trimester pregnant individual samples, processed using the Database for Annotation, Visualization and Integrated Discovery (DAVID) for Gene Ontology (GO) Cellular Component (24).

#### 2.4.3. Deconvolution Analysis to Estimate Cell/Tissue of Origin of miRNAs in Urine during Pregnancy

Previous studies have conducted deconvolution analysis of mRNA and proteomic data in uEVs and have demonstrated that the majority of protein-coding genes and proteins respectively are enriched for kidney-specific markers (25–27). We therefore focused our deconvolution analysis on the potential origin of urinary miRNAs. In order to determine the potential origin of urinary miRNAs in pregnancy, we conducted deconvolution analysis to calculate the fractional contribution of different cells and tissue types to miRNAs in maternal urine using CIBERSORTx package, which applies a linear support vector regression model to estimate proportions (28). A previously published dataset containing 438 miRNAs that were differentially expressed across different cell/tissue types, averaged, and expressed as cpm was utilized as a background gene expression profile (29). This profile included miRNAs that are highly expressed across lung, intestine, pancreas, liver, kidney, heart, brain, placenta, lymphocytes, and platelets (29). We then created an “intersect” dataset by looking at the overlap between the 438 miRNAs that were differentially expressed across cell/tissue types and the 93 miRNAs that passed a detection filter in at least 50% of pregnant female urine samples extracted using the miRCURY kit (pools and individuals, n=19), and found 80 miRNAs to overlap between the two datasets. To run CIBERSORTx using the “Impute cell fractions” function, we input a “mixture” file containing the cpm values of 80 miRNAs that passed the detection filter in pregnant urine samples (n=19), and a “signature matrix” file contained average cpm values for each cell/tissue type for the same 80 miRNAs from the background gene expression profile dataset.

### 2.5. Methods to Characterize Urinary Extracellular Vesicles

#### 2.5.1 Nanoparticle Tracking Analysis and Microfluidic Resistive Pulse Sensing

Samples analyzed using NTA and MRPS included neat and concentrated female non-pregnant urine pools. Aliquots of female non-pregnant urine pools were concentrated to 300 μL from 4 mL using 100 kDa Amicon filters (∼10 nm pore size) and resuspended in phosphate buffered saline (PBS). For NTA, “nanoparticle” or uEV concentration and size were measured using the Nanosight LM-10 (Malvern Instruments, Worcestershire, UK), and a 532 nm laser and high-sensitivity scMOS camera. For MRPS, concentration and size of uEVs was measured using the Spectradyne nCS1 and TS400 cartridges (400 nm pore size). A dilution series was prepared to determine optimal urine sample dilution (1:3, 1:9) for NTA and MRPS respectively (Supplementary Methods Figure 4). ‘Buffer only’ controls were included in each experiment and polystyrene beads (100 nm) and liposomes (Lipo100^TM^) were used as reference standards. Samples were treated with detergent (0.1% Triton X-100) to assess whether most events were ‘vesicular’ and could be disrupted by addition of a detergent. Further details of each method are included in the Supplementary Methods.

#### 2.5.2. Vesicle Flow Cytometry

Samples analyzed using vFC included neat as well as miRCURY-precipitated female non-pregnant and pregnant urine pools from 1 mL starting volume resuspended in 100 μL resuspension buffer (10x concentrated). Concentration, size, and surface marker expression of uEVs was measured by single vesicle flow cytometry using a commercial kit (vFC Assay kit, Cellarcus^TM^ Biosciences, La Jolla, CA). The vFC^TM^ EV analysis assay was performed by staining samples with vFRed^TM^, a fluorogenic membrane stain, and multiplexed antibodies to the three canonical tetraspanin EV markers, CD9, CD63, and CD81, as well as annexin V or CFSE (Carboxyfluorescein Diacetate Succinimidyl Ester) for 1h at RT. CFSE staining indicates esterase activity and internal volume of EVs. Samples were analyzed using the CytoFlexS (Beckman Coulter). Controls included ‘buffer only’, ‘reagent only’ and positive and negative controls for antigen staining. Data were validated using single stained controls, analyzed using FCS Express version 7 (De Novo Software) and calibrated using standards for vesicle size and fluorescence intensity. Optimal sample dilution was determined using a dilution series and positive and negative controls per ISEV guidelines. Further details are included in the Supplementary Methods.

## 3. RESULTS

### 3.1. Small RNA Biotype, Complexity, and Separation by exRNA Isolation Method

Small RNA libraries with ≥10,000 trimmed reads were obtained from a total of 208 urine samples encompassing male pools, and female non-pregnant and pregnant pools and individuals (“Ind”). The number of small RNA libraries for each isolation method and group are summarized in Table 1. The distribution of reads mapping to each small RNA biotype, rRNA, miRNA, tRNA, piwi-interacting RNA (piRNA), and other GENCODE sequences, as well as unmapped reads, were calculated as a percentage of total clipped reads after adapter trimming (Supplementary Figure S1 A). Three separate statistical analyses were conducted using the Kruskal-Wallis test and the Dunn’s correction for multiple comparisons: (1) % miRNA in all female urine samples extracted using each of the exRNA isolation methods, split by starting volume and dry-down (DD) for vacuum concentrated samples or no dry down (Figure 2A) (2) % miRNA in miRCURY-extracted female urine samples that were not vacuum concentrated split by pregnancy status and starting volume (Figure 2B), and (3) % tRNA in female urine samples extracted using all methods that were not vacuum concentrated (Figure 2A). The number of samples, descriptive statistics and p-values of significant results in each comparison are summarized in Supplementary Table 2. We found that (1) miRCURY-extracted samples that were not vacuum-concentrated yielded the highest average % miRNA (4.2%) compared to all other exRNA isolation methods, and this was significantly increased compared to mAdv-extracted samples that were not DD (p<0.05), but not all other exRNA isolation methods (Supplementary Table 2). There was also no increase in % miRNA in DD samples (Figure 2A). (2) Within miRCURY extracted samples that were not DD, non-pregnant pools extracted from 4 mL of starting volume yielded the highest average % miRNA (15.2%) (Figure 2B). (3) Norgen-extracted samples yielded the highest average % tRNA (34.3%), followed by ExoRNeasy (23.6%), and these were significantly increased compared to both miRCURY and mAdv-extracted samples (p<0.05) (Figure 2A). We selected the EV-associated exRNA and total exRNA extraction methods with the highest % miRNA (miRCURY and mAdv, respectively) for extraction of exRNA from the individual NP and P female urine samples (Table1).

**Figure 2:**
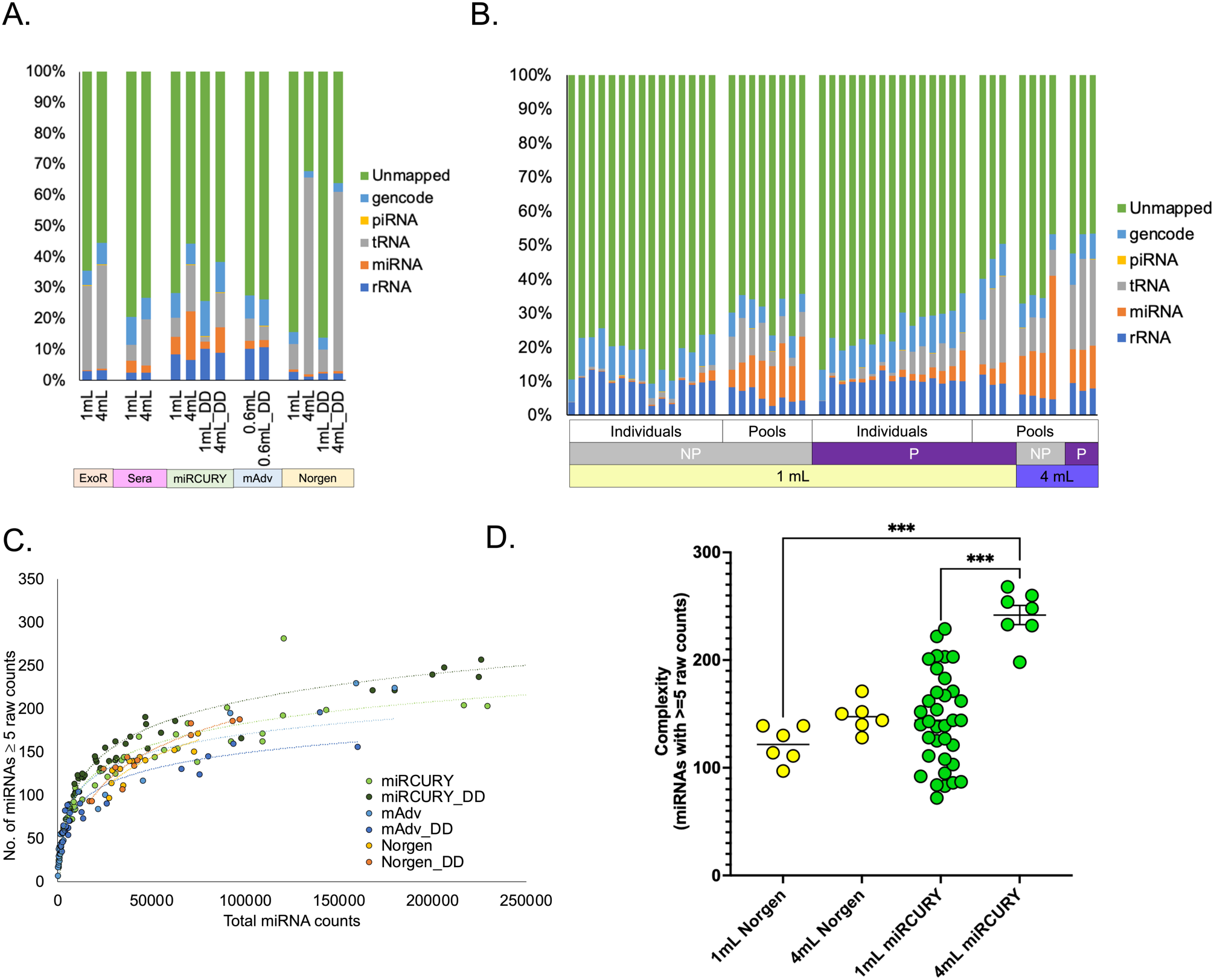
Small RNA biotypes and complexity by exRNA isolation method. **(A)** Small RNA biotypes showing rRNA, miRNA, tRNA, piRNA, other GENCODE transcripts and unmapped reads averaged across a total of 189 female urine samples expressed as a percentage of total clipped reads. Samples are split into groups according to exRNA isolation method (ExoR=ExoRNEasy, Sera=SeraMir, miRCURY, mAdv=miRNeasy Advanced, Norgen), starting volume (0.6mL, 1mL, or 4mL), and dry-down (DD) or as is. Sample numbers for each method are provided in Table 1. **(B)** Small RNA biotypes for 48 female urine samples extracted using the miRCURY method, split into groups according to individuals/pools, pregnancy status (NP=non-pregnant, P=pregnant), and starting volume (1mL or 4mL). **(C)** Complexity (individual miRNAs with ≥ 5 raw counts in each sample) split by exRNA isolation method shows a plateau with increasing total miRNA read counts, indicating optimal sequencing depth for samples by extraction method for Norgen, miRCURY, and mAdv extracted samples. **(D)** miRNA complexity of samples extracted using the miRCURY method with a starting volume of 4mL of urine was significantly higher than samples extracted from 1mL of urine using Norgen and miRCURY methods (p<0.05).

The miRNA complexity was directly correlated with the total miRNA read counts and plateaued in the range of 150-200 unique miRNAs detected for miRCURY, mAdv, and Norgen (Figure 2C). Samples extracted using ExoRNeasy and SeraMir yielded samples with markedly lower miRNA complexity compared to the other three methods, likely due to low % miRNA, which made it unfeasible to obtain adequate numbers of miRNA reads, and are represented separately (Supplementary Figure S1 B). The complexity of the libraries plateaued at ∼250 unique miRNAs with ∼200,000 total miRNA read counts for samples prepared using miRCURY, at ∼200 unique miRNAs with ∼100,000 total miRNA read counts for Norgen, and at ∼150 unique miRNAs with ∼100,000 total miRNA read counts for mAdv (Figure 2C). The average complexity for samples extracted using ExoRNeasy and SeraMir was 52 and 63 respectively, with average total miRNA read counts of ∼3000 or ∼6000, respectively (Supplementary Figure S1B). Comparison of miRNA complexity between starting volumes of 1mL and 4mL using Norgen and miRCURY methods showed that the complexity of samples was significantly higher (p<0.05) using 4mL starting volume and the miRCURY method compared to 1mL starting volume with miRCURY and 4mL starting volume with Norgen (Figure 2D).

Out of the 189 small RNA libraries from female non-pregnant and pregnant pools and individuals, 176 libraries had ≥1000 total miRNA read counts and were retained for unsupervised clustering analysis. Principal component analysis (PCA) conducted on filtered miRNAs that were present in at least 50% of samples within each exRNA isolation method indicated that samples clustered based on exRNA isolation method (Figure 3A) and whether they were pools or from individuals (Figure 3C), rather than pregnancy status (Figure 3B). Of the miRNAs that passed this filter in each kit, 16 miRNAs overlapped amongst all five methods of exRNA isolation from female non-pregnant and pregnant urine and were present in at least 50% of all samples extracted using each method (Figure 3, Table 2). Samples extracted using the Norgen method contained all but 7 of the miRNAs present in all exRNA isolation methods combined, as well as 54 miRNAs that were not detected in the samples from any of the other methods (Figure 3D).

**Figure 3:**
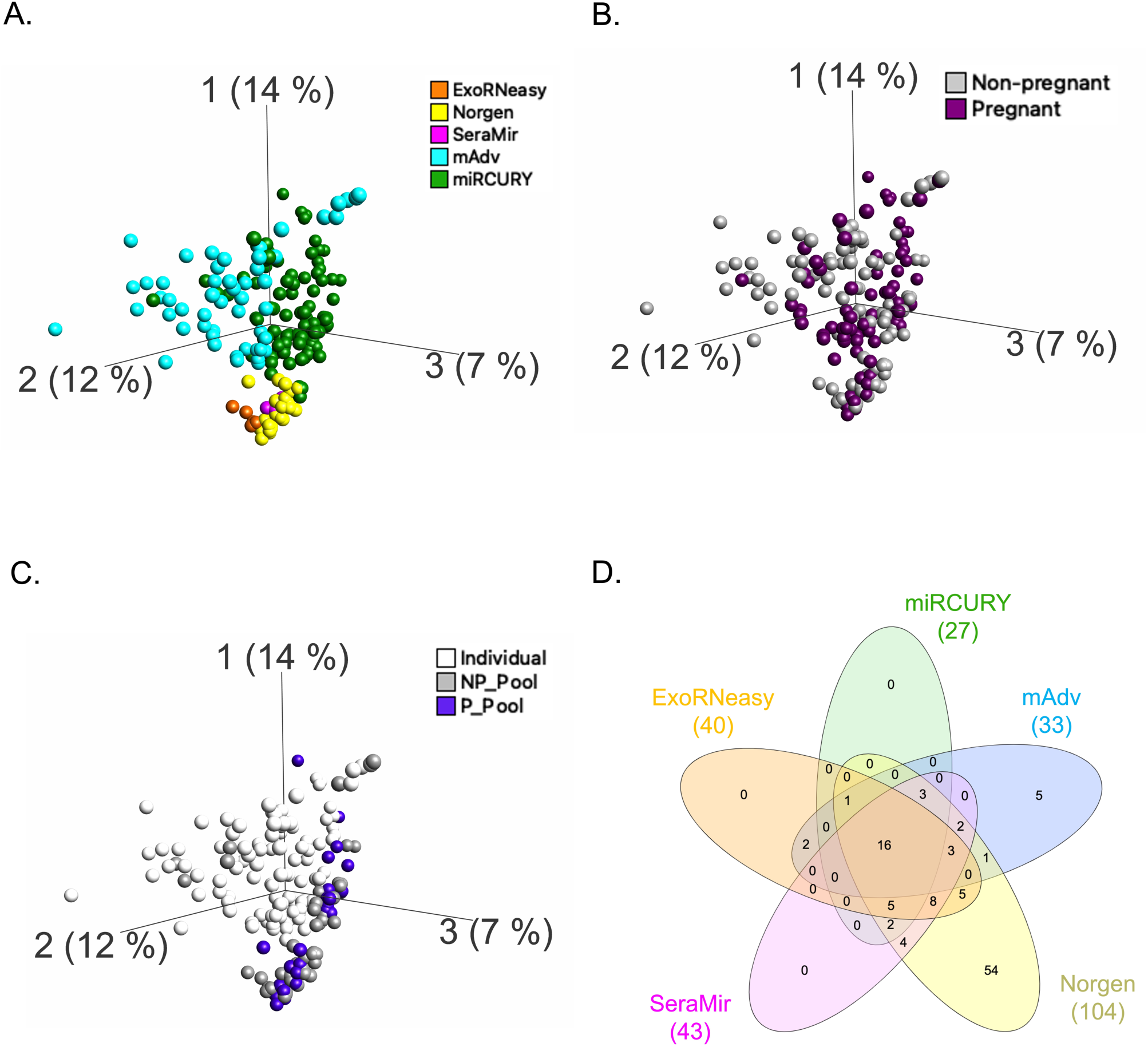
Visualization of pass-filter miRNAs across each exRNA isolation method in female urine pools and individuals. MicroRNAs were filtered across libraries (n=176) by calculating the value for 5 raw counts in counts per million (cpm) in the sample with the lowest complexity within each RNA extraction method and using this value as the cutoff for miRNAs present in at least 50% of samples extracted using each method. **(A)** Small RNA libraries extracted using different kits appear to cluster together as shown using principal component analysis (PCA) of pass-filter miRNAs in rather than **(B)** pregnancy status. **(C)** Separation of urine pools and individual samples can also be seen. **(D)** Overlap in miRNAs that passed filter within each RNA extraction method: 16 urinary extracellular miRNAs overlapped amongst all the methods used to extract female urine samples (Table 2).

**Figure 4:**
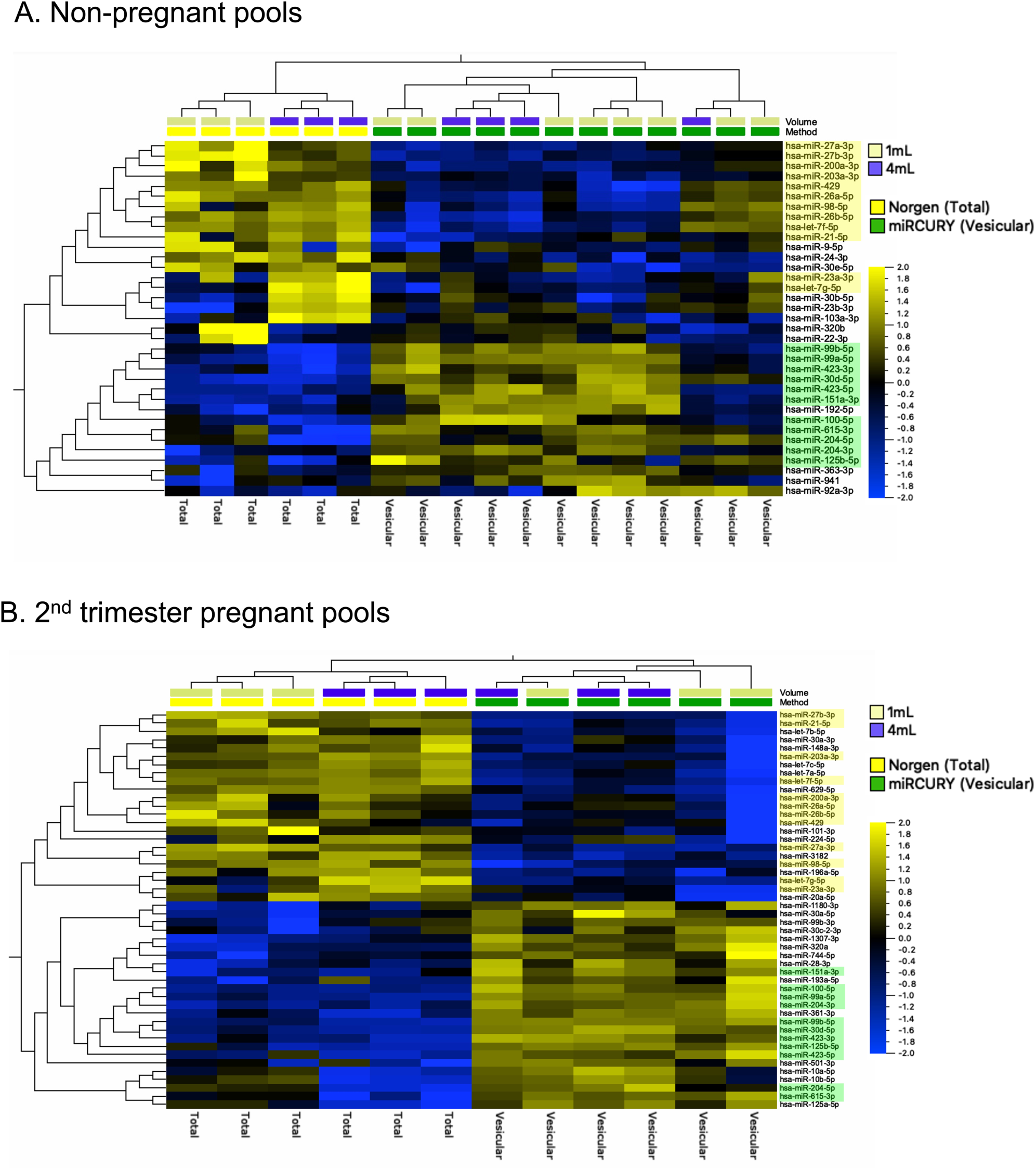
**Differentially expressed miRNAs across female non-pregnant and 2^nd^ trimester pregnant urine pools comparing total (Norgen) or vesicular (miRCURY) methods to isolate urinary exRNA**. **(A)** Comparison of non-pregnant female urine pools extracted using miRCURY (n=6) and Norgen (n=12) methods with starting volumes of 1 mL or 4 mL of urine showed that 35 miRNAs were differentially expressed (FDR<0.05). Samples hierarchically clustered by RNA extraction method. **(B)** Comparison of 2^nd^ trimester pregnant urine pools extracted using miRCURY (n=6) and Norgen (n=6) methods with starting volumes of either 1 mL or 4 mL of urine showed that 48 miRNAs were differentially expressed (FDR<0.05). Samples hierarchically clustered by RNA extraction method. Amongst the miRNAs that were differentially expressed within non-pregnant and pregnant urine pools, 23 miRNAs overlapped between the two analyses, 25 were differentially expressed only in pregnant urine pools, and 12 were differentially expressed only in non-pregnant urine pools. The 23 miRNAs that overlapped between (A) and (B) are highlighted in yellow (decreased) and green (increased).

**Table 2:** Sixteen overlapping miRNAs detected in female urine using all five methods of exRNA isolation.

| miRNA | Vesiclepedia urine miRNA entry |
| --- | --- |
| hsa-miR-99a-5p | ✓ |
| hsa-let-7a-5p |  |
| hsa-miR-192-5p | ✓ |
| hsa-miR-10b-5p | ✓ |
| hsa-miR-10a-5p | ✓ |
| hsa-miR-30a-5p | ✓ |
| hsa-miR-21-5p | ✓ |
| hsa-miR-148a-3p | ✓ |
| hsa-miR-30d-5p | ✓ |
| hsa-miR-100-5p | ✓ |
| hsa-miR-26a-5p | ✓ |
| hsa-miR-151a-3p | ✓ |
| hsa-miR-423-5p | ✓ |
| hsa-let-7f-5p |  |
| hsa-let-7b-5p |  |
| hsa-let-7i-5p |  |

Analysis of the % CV (coefficient of variation) as a function of mean miRNA expression level showed that samples extracted using Norgen had lower % CV for highly expressed miRNAs indicating that miRNAs >100CPM express low variability in Norgen-extracted samples and this method may be suitable to investigate miRNAs that are highly expressed in urine (Supplementary Figure S2). miRNeasy Advanced showed the highest % CV across all expression levels of miRNA and was therefore the least reproducible method. miRCURY, ExoRNeasy and Seramir extracted samples showed higher % CV in miRNAs with expression <10,000 CPM. % CV decreased with increasing miRNA expression for all exRNA isolation methods, indicating that miRNAs with lower expression showed higher variability in urine samples.

### 3.2. Comparisons to Identify Differentially Expressed (DE) miRNAs

#### 3.2.1. Total vs. Vesicular Extraction Methods in Non-pregnant and Pregnant Pools

Urine pool samples with a total miRNA read count of ≥25,000 were used in the differential expression analyses, which excluded samples extracted using SeraMir, ExoRNeasy, and mAdv from these analyses. Samples were split into non-pregnant and pregnant pools prior to analysis to identify DE miRNAs in samples extracted using total (Norgen, 0.44 x 10^5^ average miRNA reads) or vesicular (miRCURY, 2.57 x 10^5^ average miRNA reads) exRNA isolation methods in these two pool types. Analysis of non-pregnant urine pools extracted using the miRCURY (n=12) and Norgen (n=6) methods with starting volumes of either 1 mL or 4 mL showed that 35 miRNAs were DE (15↑ and 20↓ in vesicular exRNA isolated samples, FDR<0.05, Figure 4A). Hierarchical clustering by row and column showed that samples clustered by RNA extraction method (Figure 4A). Analysis of 2^nd^ trimester pregnant urine pools extracted using the miRCURY (n=6) and Norgen (n=6) methods with starting volumes of either 1 mL or 4 mL showed that 48 miRNAs were DE (25↑ and 23↓ in vesicular exRNA isolated samples, FDR<0.05, Figure 4B). Hierarchical clustering by row and column showed that samples clustered by RNA extraction method. Comparing the DE miRNAs in the pregnant and non-pregnant pools, 23 miRNAs overlapped: 11 miRNAs were enriched in the vesicular exRNA preparation in both pregnant and non-pregnant pools (highlighted in green, Figures 4A and 4B), while 12 miRNAs were enriched in the total exRNA preparation in both pregnant and non-pregnant pools (highlighted in yellow, Figures 4A and 4B). A complete list of miRNAs enriched following vesicular or total exRNA isolation that were unique to either non-pregnant or pregnant pools, as well as overlapping in both pools, is provided in Supplementary Table 4.

#### 3.2.2. Pregnant vs. Non-pregnant Pools and Individuals (miRCURY-extracted samples)

Since the miRCURY kit provided the highest % miRNA and most complex miRNAs amongst exRNA isolation methods, we focused on this method for further differential expression comparisons. We conducted three separate differential expression analyses to identify miRNAs that were DE in pregnancy in samples extracted using the miRCURY kit: 1) Non-pregnant vs. 2^nd^ trimester pregnant urine pools (NP and P pools); 2) Non-pregnant (NP1-5) vs. pregnant (P1-5) individuals; and 3) 2^nd^ trimester pregnant pools (average GA 25 weeks, 2^nd^ trimester) vs. 3^rd^ trimester pregnant individuals (average GA 34 weeks, 3^rd^ trimester). Urine samples with a total miRNA read count of ≥6000 were used in these comparisons, which excluded one triplicate each from individuals NP4, P4 and P5, and all triplicates from individual NP3.

Analysis of non-pregnant female (n=12) and pregnant (n=6) urine pools extracted from starting volumes of either 1 mL or 4 mL showed that 37 miRNAs were DE (14↑ and 23−↓ in 2^nd^ trimester pregnant pools, FDR<0.05, Figure 5A). Hierarchical clustering by row and column showed that samples clustered by pregnancy status (Figure 5A). Comparison of 3^rd^ trimester pregnant (n=13 samples from 5 individuals) and non-pregnant (n=11 samples from 4 individuals) samples extracted in triplicate from 1 mL of starting volume showed that 6 miRNAs were DE (3↑ and 3−↓ in 3^rd^ trimester pregnant individuals, FDR<0.05, Supplementary Table 5). One liver-specific miRNA, hsa-miR-122-5p, was significantly decreased in both 2^nd^ trimester pregnant pools and 3^rd^ trimester individual samples compared to non-pregnant pools and non-pregnant individual samples respectively (Supplementary Table 5).

**Figure 5:**
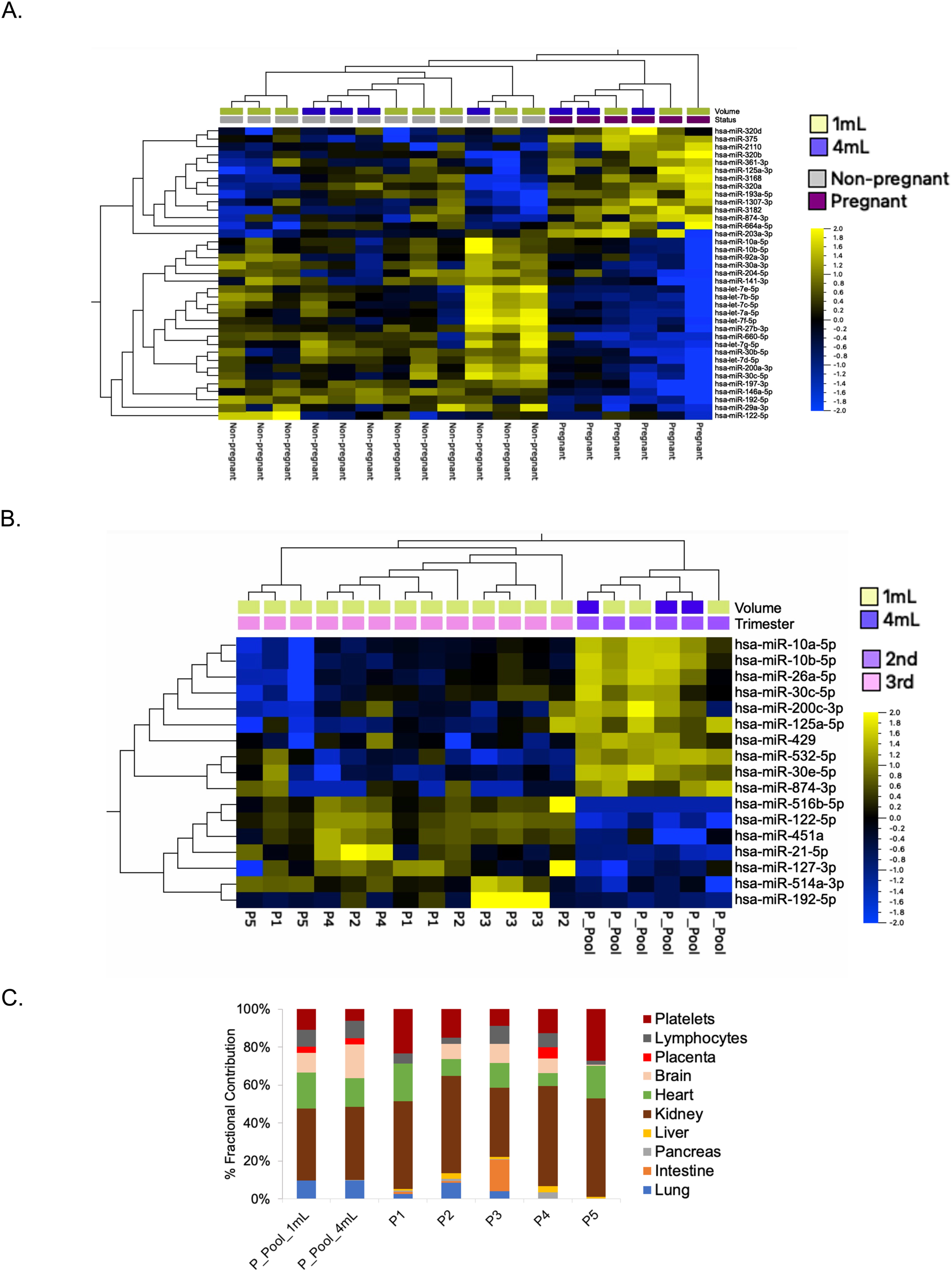
Comparison of pregnant vs. non-pregnant urine pools and individuals extracted using the miRCURY kit. **(A)** Comparison of non-pregnant female (n=12) and 2^nd^ trimester pregnant (n=6) urine pools extracted from starting volumes of either 1 mL or 4 mL showed that 37 miRNAs were differentially expressed (FDR<0.05). Samples hierarchically clustered by RNA extraction method. **(B)** Comparison of 2^nd^ trimester individual urine samples and 3^rd^ trimester urine pools extracted from 1 mL and 4 mL starting volumes showed differential expression of 17 miRNAs (FDR<0.05). Samples hierarchically clustered by trimester. **(C)** Computational deconvolution analysis showed that the majority of miRNAs in pregnant urine (n=19) may be derived from the kidney, with a smaller fraction from the placenta in pregnant pools and individual urine samples. Values are averaged across replicates.

Comparison of 2^nd^ trimester urine pools (n=6) and 3^rd^ trimester individual samples (n=13 samples from 5 individuals) extracted from 1 mL and 4 mL starting volumes showed differential expression of 17 miRNAs (7↑ and 10−↓ in 3^rd^ trimester pregnant individual samples, FDR<0.05, Figure 5B). Samples hierarchically clustered by GA (2^nd^ or 3^rd^ trimester). The miRNAs that were enriched in 2^nd^ or 3^rd^ trimester pregnant urine following miRCURY (vesicular) extraction in the above comparisons are provided in Table 3, and a complete list of miRNAs that were DE in pregnancy are provided in Supplementary Table 5.

**Table 3:**
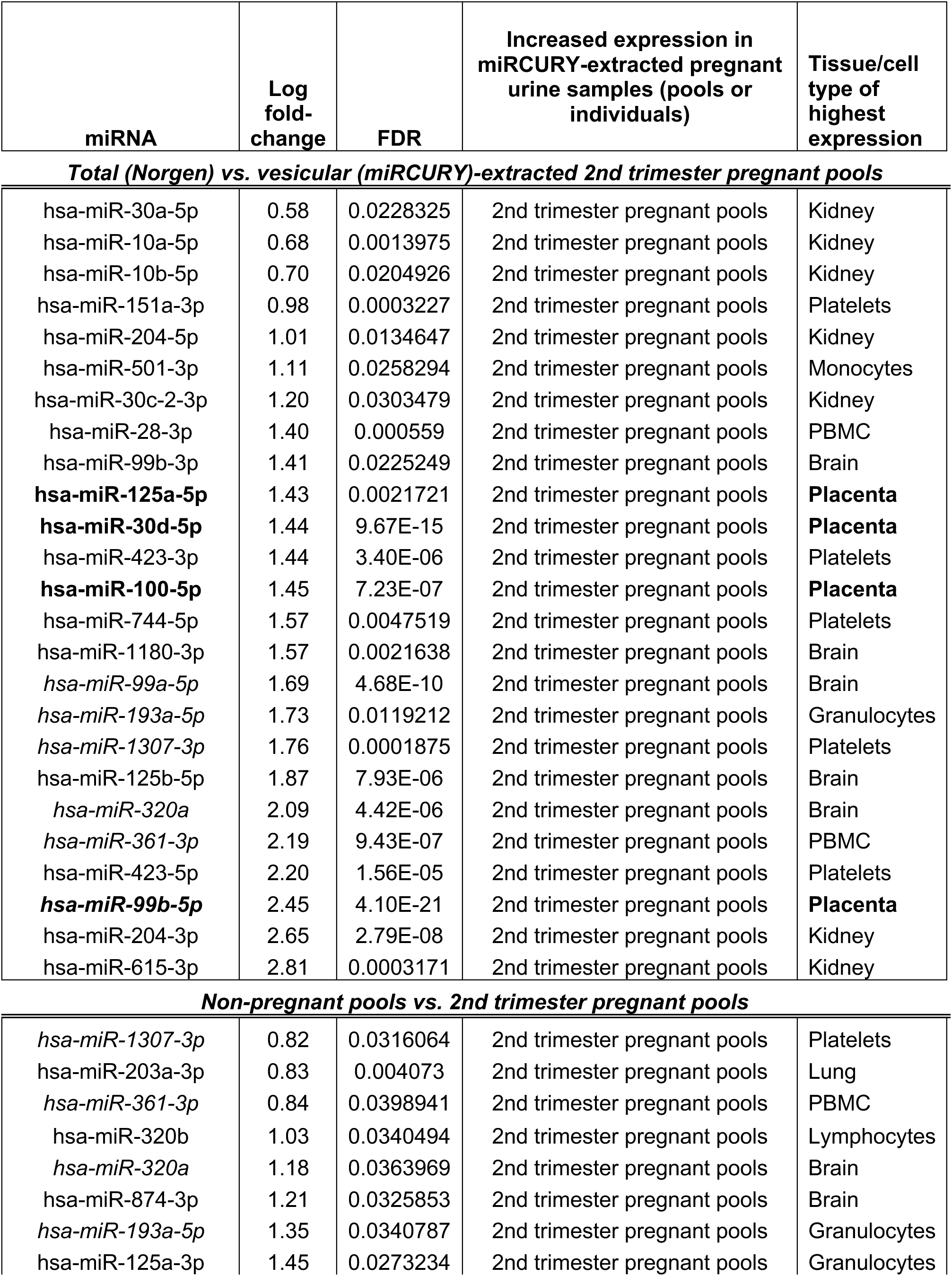
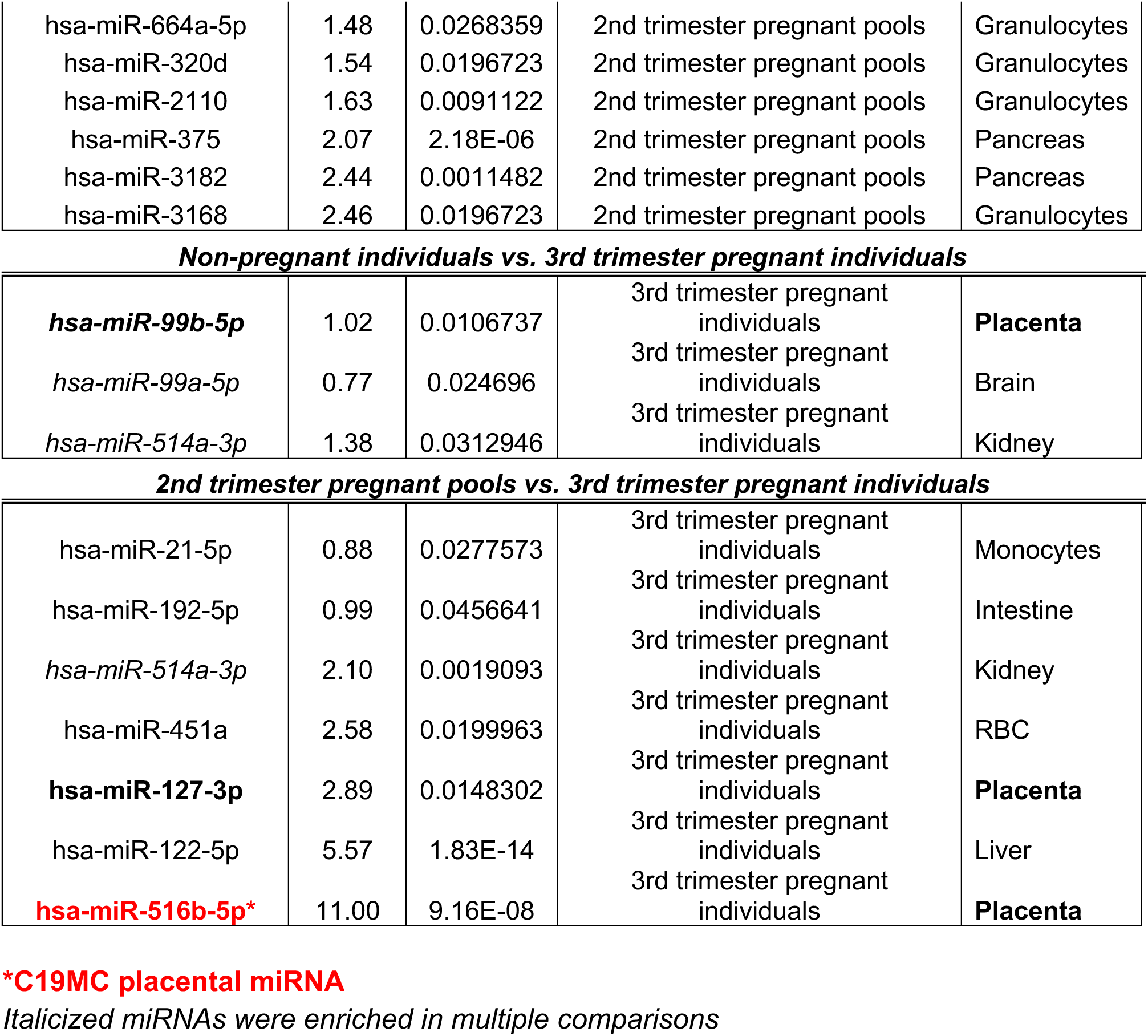
MicroRNAs overexpressed in miRCURY-extracted female urine samples (pools or individuals) along with their corresponding tissue of highest expression (29).

| miRNA | Log fold-change | FDR | Increased expression in miRCURY-extracted pregnant urine samples (pools or individuals) | Tissue/cell type of highest expression |
| --- | --- | --- | --- | --- |
| ***Total (Norgen) vs. vesicular (miRCURY)-extracted 2nd trimester pregnant pools*** |  |  |  |  |
| hsa-miR-30a-5p | 0.58 | 0.0228325 | 2nd trimester pregnant pools | Kidney |
| hsa-miR-10a-5p | 0.68 | 0.0013975 | 2nd trimester pregnant pools | Kidney |
| hsa-miR-10b-5p | 0.70 | 0.0204926 | 2nd trimester pregnant pools | Kidney |
| hsa-miR-151a-3p | 0.98 | 0.0003227 | 2nd trimester pregnant pools | Platelets |
| hsa-miR-204-5p | 1.01 | 0.0134647 | 2nd trimester pregnant pools | Kidney |
| hsa-miR-501-3p | 1.11 | 0.0258294 | 2nd trimester pregnant pools | Monocytes |
| hsa-miR-30c-2-3p | 1.20 | 0.0303479 | 2nd trimester pregnant pools | Kidney |
| hsa-miR-28-3p | 1.40 | 0.000559 | 2nd trimester pregnant pools | PBMC |
| hsa-miR-99b-3p | 1.41 | 0.0225249 | 2nd trimester pregnant pools | Brain |
| **hsa-miR-125a-5p** | 1.43 | 0.0021721 | 2nd trimester pregnant pools | **Placenta** |
| **hsa-miR-30d-5p** | 1.44 | 9.67E-15 | 2nd trimester pregnant pools | **Placenta** |
| hsa-miR-423-3p | 1.44 | 3.40E-06 | 2nd trimester pregnant pools | Platelets |
| **hsa-miR-100-5p** | 1.45 | 7.23E-07 | 2nd trimester pregnant pools | **Placenta** |
| hsa-miR-744-5p | 1.57 | 0.0047519 | 2nd trimester pregnant pools | Platelets |
| hsa-miR-1180-3p | 1.57 | 0.0021638 | 2nd trimester pregnant pools | Brain |
| *hsa-miR-99a-5p* | 1.69 | 4.68E-10 | 2nd trimester pregnant pools | Brain |
| *hsa-miR-193a-5p* | 1.73 | 0.0119212 | 2nd trimester pregnant pools | Granulocytes |
| *hsa-miR-1307-3p* | 1.76 | 0.0001875 | 2nd trimester pregnant pools | Platelets |
| hsa-miR-125b-5p | 1.87 | 7.93E-06 | 2nd trimester pregnant pools | Brain |
| *hsa-miR-320a* | 2.09 | 4.42E-06 | 2nd trimester pregnant pools | Brain |
| *hsa-miR-361-3p* | 2.19 | 9.43E-07 | 2nd trimester pregnant pools | PBMC |
| hsa-miR-423-5p | 2.20 | 1.56E-05 | 2nd trimester pregnant pools | Platelets |
| **hsa-miR-99b-5p** | 2.45 | 4.10E-21 | 2nd trimester pregnant pools | **Placenta** |
| hsa-miR-204-3p | 2.65 | 2.79E-08 | 2nd trimester pregnant pools | Kidney |
| hsa-miR-615-3p | 2.81 | 0.0003171 | 2nd trimester pregnant pools | Kidney |
| ***Non-pregnant pools vs. 2nd trimester pregnant pools*** |  |  |  |  |
| *hsa-miR-1307-3p* | 0.82 | 0.0316064 | 2nd trimester pregnant pools | Platelets |
| hsa-miR-203a-3p | 0.83 | 0.004073 | 2nd trimester pregnant pools | Lung |
| *hsa-miR-361-3p* | 0.84 | 0.0398941 | 2nd trimester pregnant pools | PBMC |
| hsa-miR-320b | 1.03 | 0.0340494 | 2nd trimester pregnant pools | Lymphocytes |
| *hsa-miR-320a* | 1.18 | 0.0363969 | 2nd trimester pregnant pools | Brain |
| hsa-miR-874-3p | 1.21 | 0.0325853 | 2nd trimester pregnant pools | Brain |
| *hsa-miR-193a-5p* | 1.35 | 0.0340787 | 2nd trimester pregnant pools | Granulocytes |
| hsa-miR-125a-3p | 1.45 | 0.0273234 | 2nd trimester pregnant pools | Granulocytes |
| hsa-miR-664a-5p | 1.48 | 0.0268359 | 2nd trimester pregnant pools | Granulocytes |
| hsa-miR-320d | 1.54 | 0.0196723 | 2nd trimester pregnant pools | Granulocytes |
| hsa-miR-2110 | 1.63 | 0.0091122 | 2nd trimester pregnant pools | Granulocytes |
| hsa-miR-375 | 2.07 | 2.18E-06 | 2nd trimester pregnant pools | Pancreas |
| hsa-miR-3182 | 2.44 | 0.0011482 | 2nd trimester pregnant pools | Pancreas |
| hsa-miR-3168 | 2.46 | 0.0196723 | 2nd trimester pregnant pools | Granulocytes |
| ***Non-pregnant individuals vs. 3rd trimester pregnant individuals*** |  |  |  |  |
| ***hsa-miR-99b-5p*** | 1.02 | 0.0106737 | 3rd trimester pregnant individuals | **Placenta** |
| *hsa-miR-99a-5p* | 0.77 | 0.024696 | 3rd trimester pregnant individuals | Brain |
| *hsa-miR-514a-3p* | 1.38 | 0.0312946 | 3rd trimester pregnant individuals | Kidney |
| ***2nd trimester pregnant pools vs. 3rd trimester pregnant individuals*** |  |  |  |  |
| hsa-miR-21-5p | 0.88 | 0.0277573 | 3rd trimester pregnant individuals | Monocytes |
| hsa-miR-192-5p | 0.99 | 0.0456641 | 3rd trimester pregnant individuals | Intestine |
| *hsa-miR-514a-3p* | 2.10 | 0.0019093 | 3rd trimester pregnant individuals | Kidney |
| hsa-miR-451a | 2.58 | 0.0199963 | 3rd trimester pregnant individuals | RBC |
| ***hsa-miR-127-3p*** | 2.89 | 0.0148302 | 3rd trimester pregnant individuals | **Placenta** |
| hsa-miR-122-5p | 5.57 | 1.83E-14 | 3rd trimester pregnant individuals | Liver |
| ***hsa-miR-516b-5p\**** | 11.00 | 9.16E-08 | 3rd trimester pregnant individuals | **Placenta** |

**\*C19MC placental miRNA**
*Italicized miRNAs were enriched in multiple comparisons*

Deconvolution analysis using CIBERSORTx showed that the majority of pass-filter miRNAs in miRCURY-extracted pregnant urine samples (pools and individuals, n=19) are highly expressed in kidney (45±2.7%), followed by miRNAs expressed in platelets (14.9±2.9%) and heart (14.2±1.9%). Placental miRNAs accounted for an average of 1.7±0.9% in pregnant urine pools and individual urine samples. Other cell and tissue types including brain, lymphocytes, lung, liver, intestine, and pancreas may also contribute miRNAs to urine in pregnancy (Figure 5C).

### 3.2. Long RNA Mapping, Complexity, and Separation by Library Preparation Method

The number of long RNA libraries split by library prep method and group are summarized in Table 1. Comparing SMARTSeq v4 and Pico v2, the SMARTSeq v4 gave higher uniquely mapped reads across all methods of exRNA isolation (Figure 6A). Within SMARTSeq v4, miRCURY exRNA isolation yielded the most uniquely mapped reads followed by Norgen, and the miRCURY method yielded significantly higher uniquely mapped reads compared to ExoRNeasy but not other exRNA isolation methods (Supplementary Figure S3B). The mRNA complexity, calculated as the number of individual protein-coding mRNAs with raw counts ≥ 5 following counting of reads that were uniquely mapped to exons using featureCounts, was directly correlated with the total mRNA read count and plateaus with increased read depth for the SMARTSeq v4 lib prep kit. The complexity of the remaining long RNA libraries prepared using the Pico v2 kit (n=19) was significantly lower compared to the SMARTSeq v4 method (n=67) (p<0.05, Figure 6B). The average mRNA complexity of samples prepared using the SMARTSeq v4 kit was ∼7500 unique mRNAs, while average complexity of samples prepared using the Pico v2 kit was ∼2600 mRNAs (Figure 6B, Supplementary Figure S3 A). Comparison of complexity between starting volumes of 1mL and 4mL using Norgen and miRCURY methods showed that the mRNA complexity of samples was significantly increased (p<0.05) in 4 mL vs. 1 mL miRCURY; 4 mL Norgen vs. 1 mL miRCURY; and 1mL Norgen vs. 1mL miRCURY, but not between 1 mL vs. 4 mL Norgen extractions (Figure 6D). Unsupervised PCA with the mRNAs that passed filter within each library prep method indicated that samples clustered based on long RNA library prep method (Figure 6C). Samples with <20,000 total mRNA reads or complexity <1000 mRNAs were removed from subsequent differential expression analysis.

**Figure 6:**
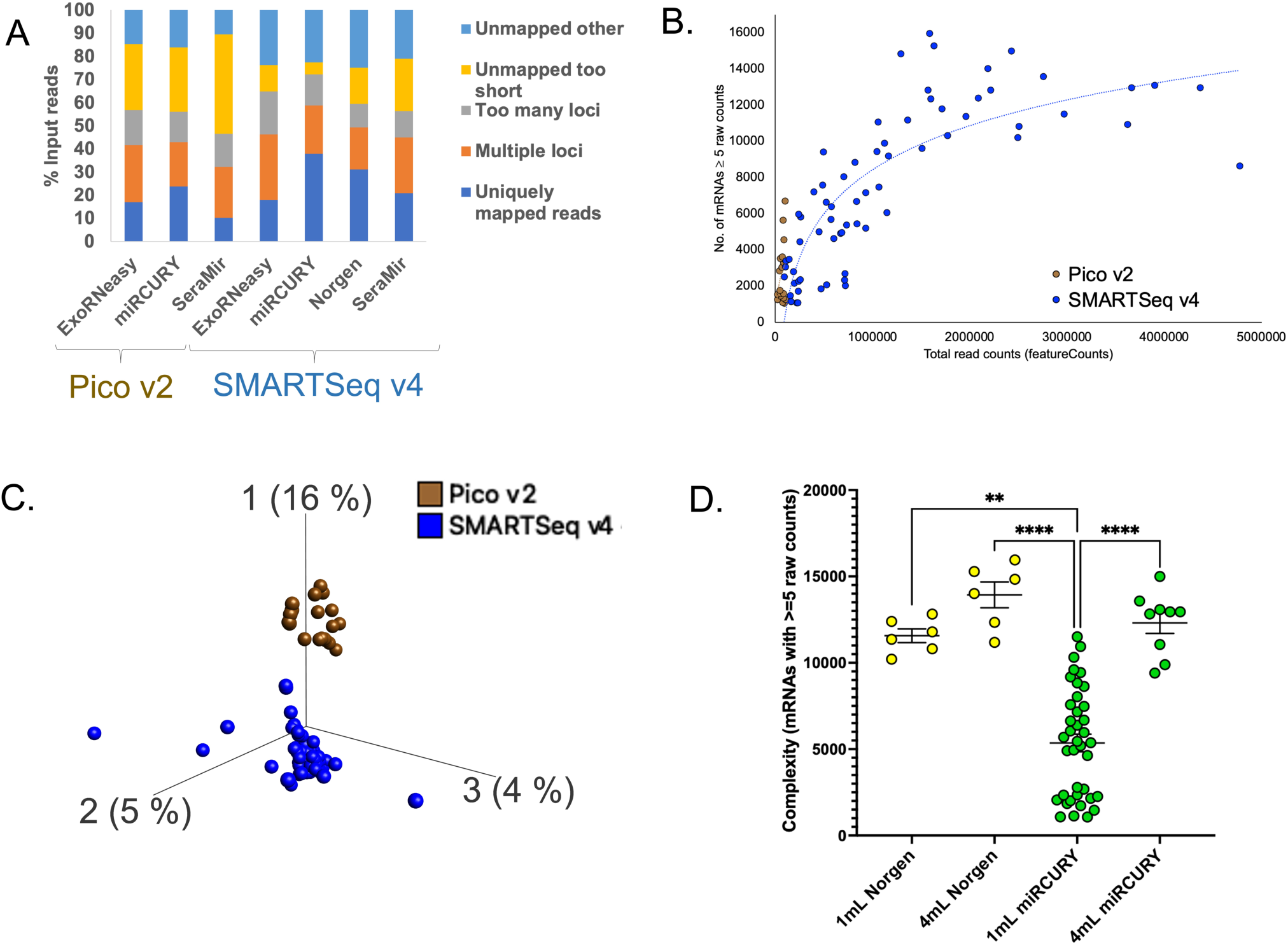
Long RNA biotypes, mRNA complexity, and separation by long RNA library preparation method. **(A)** Long RNA mapping after STAR alignment and featureCounts expressed as a percentage of total input reads, split into groups according to exRNA isolation method and library preparation method (total n=97 long RNA libraries). **(B)** Complexity, defined as individual mRNAs with ≥ 5 raw counts in each sample, split by library preparation method showing higher complexity of samples prepared using the SMARTSeq v4 Ultra Low Input Library Prep Kit compared to the Pico v2 kit. **(C)** PCA plot of pass-filter mRNAs, filtered by calculating 5 raw mRNA counts in cpm over the total read counts in the lowest complexity sample within each library preparation method, showing separation of mRNA features by library preparation kit. **(D)** mRNA complexity of samples extracted from 4 mL starting volume of urine using both miRCURY and Norgen methods and 1 mL using the Norgen method was significantly higher (p<0.05) than miRCURY-extracted samples using 1 mL starting volume of urine. All samples were prepared using the SMARTSeq v4 Ultra Low Input Library Prep Kit.

### 3.3. Comparisons to Identify Differentially Expressed mRNAs

#### 3.3.1. Total vs. Vesicular Extraction Methods in Non-pregnant and Pregnant Pools

We concentrated on Norgen (Norgen, 1.8 x 10^6^ average mRNA reads) and miRCURY (miRCURY, 2 x 10^6^ average mRNA reads) extracted exRNA in combination with the SMARTSeq v4 sequencing method for further analysis. Samples were split into non-pregnant and pregnant pools prior to analysis to identify DE protein-coding mRNAs in samples in these two pool types. Two separate analyses were conducted: 1) miRCURY vs. Norgen-extracted non-pregnant urine pools, and 2) miRCURY vs. Norgen-extracted pregnant urine pools. Analysis of non-pregnant urine pools extracted using miRCURY (n=14) and Norgen (n=6) methods with starting volumes of either 1 mL or 4 mL showed that 176 protein coding mRNAs were DE (13↑ and 163−↓ in vesicular exRNA isolated samples, FDR<0.05). Analysis of 2^nd^ trimester pregnant urine pools extracted using miRCURY (n=6) and Norgen (n=6) methods with starting volumes of either 1 mL or 4 mL indicated that 75 protein coding mRNAs were DE (9↑ and 66−↓ in vesicular exRNA isolated samples, FDR<0.05). Out of the DE mRNAs in pregnant and non-pregnant pools, 8 overlapped: 2 were enriched in the vesicular exRNA preparationand 6 were enriched in the total exRNA preparation. A complete list of protein-coding mRNAs enriched following vesicular or total exRNA isolation that were unique to either non-pregnant or pregnant pools, as well as overlapping in both pools, are provided in Supplementary Table 6.

#### 3.3.2. Pregnant vs. Non-pregnant Pools and Individuals (miRCURY-extracted samples)

We conducted three separate analyses to identify mRNAs that were differentially expressed in pregnancy, in samples extracted using the miRCURY kit: 1) Non-pregnant vs. 2^nd^ trimester pregnant urine pools (NP and P pools); 2) Non-pregnant (NP1-5) vs. 3^rd^ trimester pregnant (P1-5) individuals; and 3) 2^nd^ trimester pregnant pools (average GA 25 weeks) vs. 3^rd^ trimester pregnant individuals (average GA 34 weeks). Urine samples with total mRNA read counts <200,000 were removed from these comparisons. Analysis of non-pregnant (n=15) and 2^nd^ trimester pregnant (n=6) urine pools extracted from starting volumes of either 1 mL or 4 mL showed that 50 protein coding mRNAs were DE (44↑ and 6−↓ in 2^nd^ trimester pregnant pools, FDR<0.05). Comparison of 3^rd^ trimester pregnant (n=10 samples from 4 individuals) and non-pregnant individuals (n=8 samples from 5 individuals) extracted in triplicate from 1 mL of starting volume showed that 208 protein coding mRNAs were DE (204↑ and 4−↓ in 3^rd^ trimester pregnant individuals, FDR<0.05). Finally, comparison of 2^nd^ trimester urine pools (n=6) and 3^rd^ trimester individual samples (n=10 samples from 4 individuals) extracted from 1 mL and 4 mL starting volumes showed differential expression of 230 protein coding mRNAs (5↑ and 225−↓ in 3^rd^ trimester pregnant individual samples, FDR<0.05). A complete list of mRNAs that were DE in pregnancy in these comparisons are provided in Supplementary Table 7.

Analysis of molecular pathways involving mRNAs that were increased in the first two comparisons: 1) Non-pregnant vs. 2^nd^ trimester pregnant pools, and 2) Non-pregnant (NP1-5) vs. 3^rd^ trimester pregnant (P1-5) individuals using the DAVID database indicated that upregulated mRNAs in pregnant urine in both these comparisons were significantly enriched for the Gene Ontology pathway “GO: 0070062: Extracellular Exosome” Cellular Component (FDR<0.05) pathway (Table 4). This included 16 mRNAs that were increased in 2^nd^ trimester pregnant compared to non-pregnant urine pools (Figure 7A), and 45 mRNAs that were increased in 3^rd^ trimester pregnant urine compared to non-pregnant urine samples from individuals (P1-5, NP1-5, Figure 7B).

**Figure 7:**
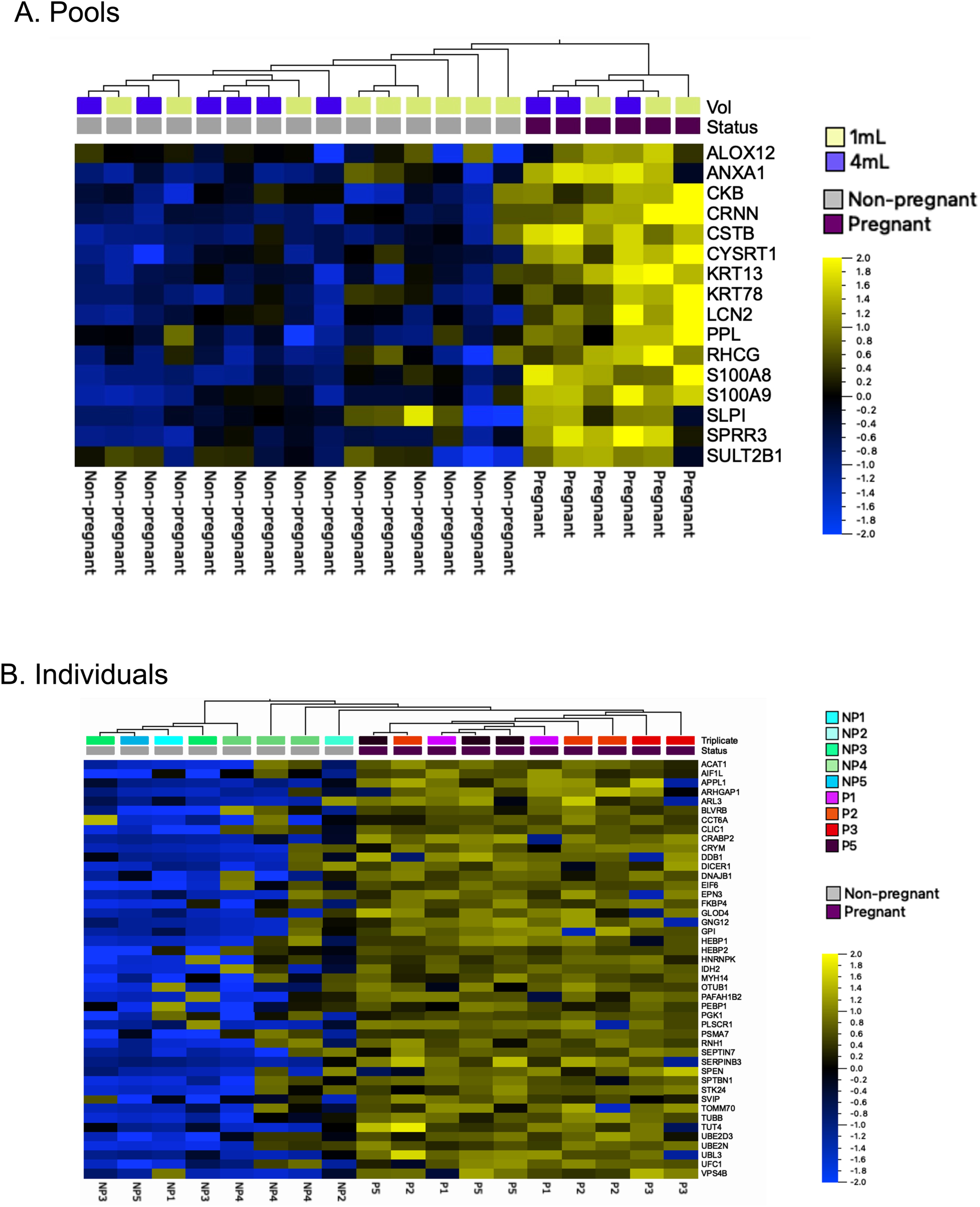
Gene Ontology pathway “GO: 0070062: Extracellular Exosome” Cellular Component pathway was significantly enriched (FDR<0.05) amongst mRNAs that were upregulated in pregnant urine extracted using the miRCURY method. **(A)** Non-pregnant vs. 2^nd^ trimester pregnant pools: Out of 44 mRNAs that were increased in 2^nd^ trimester urine pools, 16 were associated with “Extracellular Exosome” (1 mL or 4 mL starting volume). **(B)** Non-pregnant vs. 3^rd^ trimester pregnant individual samples: Out of 208 mRNAs that were increased in 3^rd^ trimester individual urine samples, 45 mRNAs were associated with “Extracellular Exosome” (1 mL starting volume).

**Table 4:** Protein coding mRNAs that were significantly increased in miRCURY-extracted pregnant urine samples and enriched in Gene Ontology pathway “GO:0070062: Extracellular Exosome” (FDR<0.05).

| ENSEMBL ID | Gene Name | Gene symbol | Increased expression in pregnant urine samples |
| --- | --- | --- | --- |
| ***Non-pregnant pools vs. 2nd trimester pregnant pools*** |  |  |  |
| ENSG00000140519 | Rh family C glycoprotein | *RHCG* | 2nd trimester pregnant pools |
| ENSG00000143546 | S100 calcium binding protein A8 | *S100A8* | 2nd trimester pregnant pools |
| ENSG00000163220 | S100 calcium binding protein A9 | *S100A9* | 2nd trimester pregnant pools |
| ENSG00000135046 | annexin A1 | *ANXA1* | 2nd trimester pregnant pools |
| ENSG00000108839 | arachidonate 12-lipoxygenase, 12S type | *ALOX12* | 2nd trimester pregnant pools |
| ENSG00000143536 | cornulin | *CRNN* | 2nd trimester pregnant pools |
| ENSG00000166165 | creatine kinase B | *CKB* | 2nd trimester pregnant pools |
| ENSG00000160213 | cystatin B | *CSTB* | 2nd trimester pregnant pools |
| ENSG00000197191 | cysteine rich tail 1 | *CYSRT1* | 2nd trimester pregnant pools |
| ENSG00000171401 | keratin 13 | *KRT13* | 2nd trimester pregnant pools |
| ENSG00000170423 | keratin 78 | *KRT78* | 2nd trimester pregnant pools |
| ENSG00000148346 | lipocalin 2 | *LCN2* | 2nd trimester pregnant pools |
| ENSG00000118898 | periplakin | *PPL* | 2nd trimester pregnant pools |
| ENSG00000124107 | secretory leukocyte peptidase inhibitor | *SLPI* | 2nd trimester pregnant pools |
| ENSG00000163209 | small proline rich protein 3 | *SPRR3* | 2nd trimester pregnant pools |
| ENSG00000088002 | sulfotransferase family 2B member 1 | *SULT2B1* | 2nd trimester pregnant pools |
| ***Non-pregnant individuals vs. 3rd trimester pregnant individuals*** |  |  |  |
| ENSG00000138175 | ADP ribosylation factor like GTPase 3 | *ARL3* | 3rd trimester pregnant individuals |
| ENSG00000132002 | DnaJ heat shock protein family | *Hsp40 member B1* | 3rd trimester pregnant individuals |
| ENSG00000004478 | FKBP prolyl isomerase 4 | *FKBP4* | 3rd trimester pregnant individuals |
| ENSG00000172380 | G protein subunit gamma 12 | *GNG12* | 3rd trimester pregnant individuals |
| ENSG00000167770 | OTU deubiquitinase, ubiquitin aldehyde binding 1 | *OTUB1* | 3rd trimester pregnant individuals |
| ENSG00000175220 | Rho GTPase activating protein 1 | *ARHGAP1* | 3rd trimester pregnant individuals |
| ENSG00000075239 | acetyl-CoA acetyltransferase 1 | *ACAT1* | 3rd trimester pregnant individuals |
| ENSG00000157500 | adaptor protein, phosphotyrosine interacting with PH domain and leucine zipper 1 | *APPL1* | 3rd trimester pregnant individuals |
| ENSG00000126878 | allograft inflammatory factor 1 like | *AIF1L* | 3rd trimester pregnant individuals |
| ENSG00000090013 | biliverdin reductase B | *BLVRB* | 3rd trimester pregnant individuals |
| ENSG00000143320 | cellular retinoic acid binding protein 2 | *CRABP2* | 3rd trimester pregnant individuals |
| ENSG00000146731 | chaperonin containing TCP1 subunit 6A | *CCT6A* | 3rd trimester pregnant individuals |
| ENSG00000213719 | chloride intracellular channel 1 | *CLIC1* | 3rd trimester pregnant individuals |
| ENSG00000103316 | crystallin mu | *CRYM* | 3rd trimester pregnant individuals |
| ENSG00000167986 | damage specific DNA binding protein 1 | *DDB1* | 3rd trimester pregnant individuals |
| ENSG00000100697 | dicer 1, ribonuclease III | *DICER1* | 3rd trimester pregnant individuals |
| ENSG00000049283 | epsin 3 | *EPN3* | 3rd trimester pregnant individuals |
| ENSG00000242372 | eukaryotic translation initiation factor 6 | *EIF6* | 3rd trimester pregnant individuals |
| ENSG00000105220 | glucose-6-phosphate isomerase | *GPI* | 3rd trimester pregnant individuals |
| ENSG00000167699 | glyoxalase domain containing 4 | *GLOD4* | 3rd trimester pregnant individuals |
| ENSG00000013583 | heme binding protein 1 | *HEBP1* | 3rd trimester pregnant individuals |
| ENSG00000051620 | heme binding protein 2 | *HEBP2* | 3rd trimester pregnant individuals |
| ENSG00000165119 | heterogeneous nuclear ribonucleoprotein K | *HNRNPK* | 3rd trimester pregnant individuals |
| ENSG00000182054 | isocitrate dehydrogenase | *NADP* | 3rd trimester pregnant individuals |
| ENSG00000105357 | myosin heavy chain 14 | *MYH14* | 3rd trimester pregnant individuals |
| ENSG00000089220 | phosphatidylethanolamine binding protein 1 | *PEBP1* | 3rd trimester pregnant individuals |
| ENSG00000102144 | phosphoglycerate kinase 1 | *PGK1* | 3rd trimester pregnant individuals |
| ENSG00000188313 | phospholipid scramblase 1 | *PLSCR1* | 3rd trimester pregnant individuals |
| ENSG00000168092 | platelet activating factor acetylhydrolase 1b catalytic subunit 2 | *PAFAH1B2* | 3rd trimester pregnant individuals |
| ENSG00000101182 | proteasome 20S subunit alpha 7 | *PSMA7* | 3rd trimester pregnant individuals |
| ENSG00000023191 | ribonuclease/angiogenin inhibitor 1 | *RNH1* | 3rd trimester pregnant individuals |
| ENSG00000122545 | septin 7 | *SEPTIN7* | 3rd trimester pregnant individuals |
| ENSG00000102572 | serine/threonine kinase 24 | *STK24* | 3rd trimester pregnant individuals |
| ENSG00000057149 | serpin family B member 3 | *SERPINB3* | 3rd trimester pregnant individuals |
| ENSG00000198168 | small VCP interacting protein | *SVIP* | 3rd trimester pregnant individuals |
| ENSG00000115306 | spectrin beta, non-erythrocytic 1 | *SPTBN1* | 3rd trimester pregnant individuals |
| ENSG00000065526 | spen family transcriptional repressor | *SPEN* | 3rd trimester pregnant individuals |
| ENSG00000134744 | terminal uridylyl transferase 4 | *TUT4* | 3rd trimester pregnant individuals |
| ENSG00000154174 | translocase of outer mitochondrial membrane 70 | *TOMM70* | 3rd trimester pregnant individuals |
| ENSG00000196230 | tubulin beta class I | *TUBB* | 3rd trimester pregnant individuals |
| ENSG00000109332 | ubiquitin conjugating enzyme E2 D3 | *UBE2D3* | 3rd trimester pregnant individuals |
| ENSG00000177889 | ubiquitin conjugating enzyme E2 N | *UBE2N* | 3rd trimester pregnant individuals |
| ENSG00000122042 | ubiquitin like 3 | *UBL3* | 3rd trimester pregnant individuals |
| ENSG00000143222 | ubiquitin-fold modifier conjugating enzyme 1 | *UFC1* | 3rd trimester pregnant individuals |
| ENSG00000119541 | vacuolar protein sorting 4 homolog B | *VPS4B* | 3rd trimester pregnant individuals |

### 3.4. Characterization of Urinary Extracellular Vesicles

#### 3.4.1 Nanoparticle Tracking Analysis and Microfluidic Resistive Pulse Sensing of Non-Pregnant Female Urine Pools

We used Nanoparticle Tracking Analysis (NTA) and Microfluidic Resistive Pulse Sensing (MRPS) to characterize uEVs prior to applying exRNA isolation methods in non-pregnant female urine pools in order to estimate size and concentration of uEVs. The use of two methods with different limits of detection provides a robust estimate of size distribution and number of uEVs prior to enrichment for RNA-Seq. The size range of urinary nanoparticles detected using MRPS was 65-250nm, with a median diameter of 96nm (Figure 8B) and concentration of 5.6 x 10^7^ particles/μL. In contrast to NTA, over 95% of events detected using MRPS were detergent labile, indicating that most events detected using this method were likely EVs. The size range of “nanoparticles” in urine estimated using NTA in neat non-pregnant female urine pools concentrated using Amicon filters was 50-250nm, with a median diameter of 124 nm (Figure 8D) and concentration of 9.6 x 10^6^ particles/μL. Fewer than 5% of events measured using NTA were disrupted by addition of detergent, indicating that the large majority of nanoparticles detected using this method in urine are non-vesicular. These detergent-resistant nanoparticles may represent Tamm-Horsfall (uromodulin) protein aggregates.

**Figure 8:**
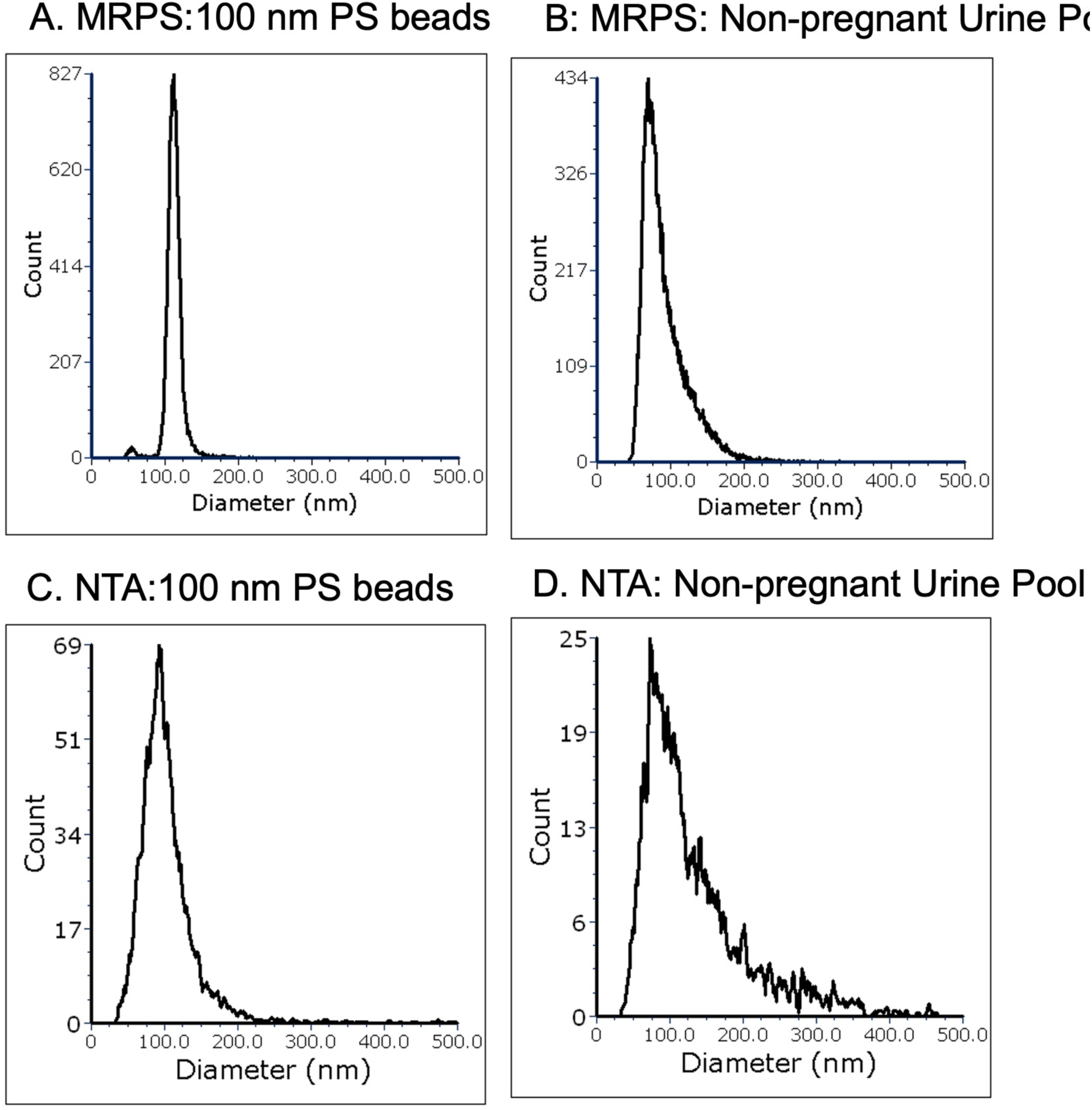
Microfluidic Resistive Pulse Sensing (MRPS) and Nanoparticle Tracking Analysis (NTA) of cell-free non-pregnant female urine pools. **(A)** and **(B)** MRPS showing the mean diameter (nm) and concentration of neat female non-pregnant urine pools along with 100nm polystyrene (PS) beads standard. **(C)** and **(D)** NTA showing the mean diameter (nm) and concentration of neat female non-pregnant urine pools along with 100nm polystyrene (PS) beads standard.

#### 3.4.2 Vesicle Flow Cytometry of Pregnant and Non-Pregnant Female Urine Pools

We applied vFC to characterize staining for antibodies to EV-associated tetraspanins (CD63, CD9, and CD81), annexin V and CFSE in neat pregnant and non-pregnant female urine pools, as well as the same pools enriched using the miRCURY kit, since the miRCURY method provided the most reproducible RNA-Seq results across exRNA isolation methods. EVs in neat and miRCURY (PEG)-precipitated pregnant and non-pregnant female urine pools had a median diameter of 122-128 nm. EVs from both non-pregnant and pregnant pooled urine showed low positivity for the tetraspanins CD63, CD9, and CD81, and positivity for annexin V between 10.2-35.6%, with higher annexin positivity in uEVs concentrated by PEG precipitation compared to neat (Figure 9). Urine EVs in all sample types showed 22.5-26.8% positivity for CFSE.

**Figure 9:**
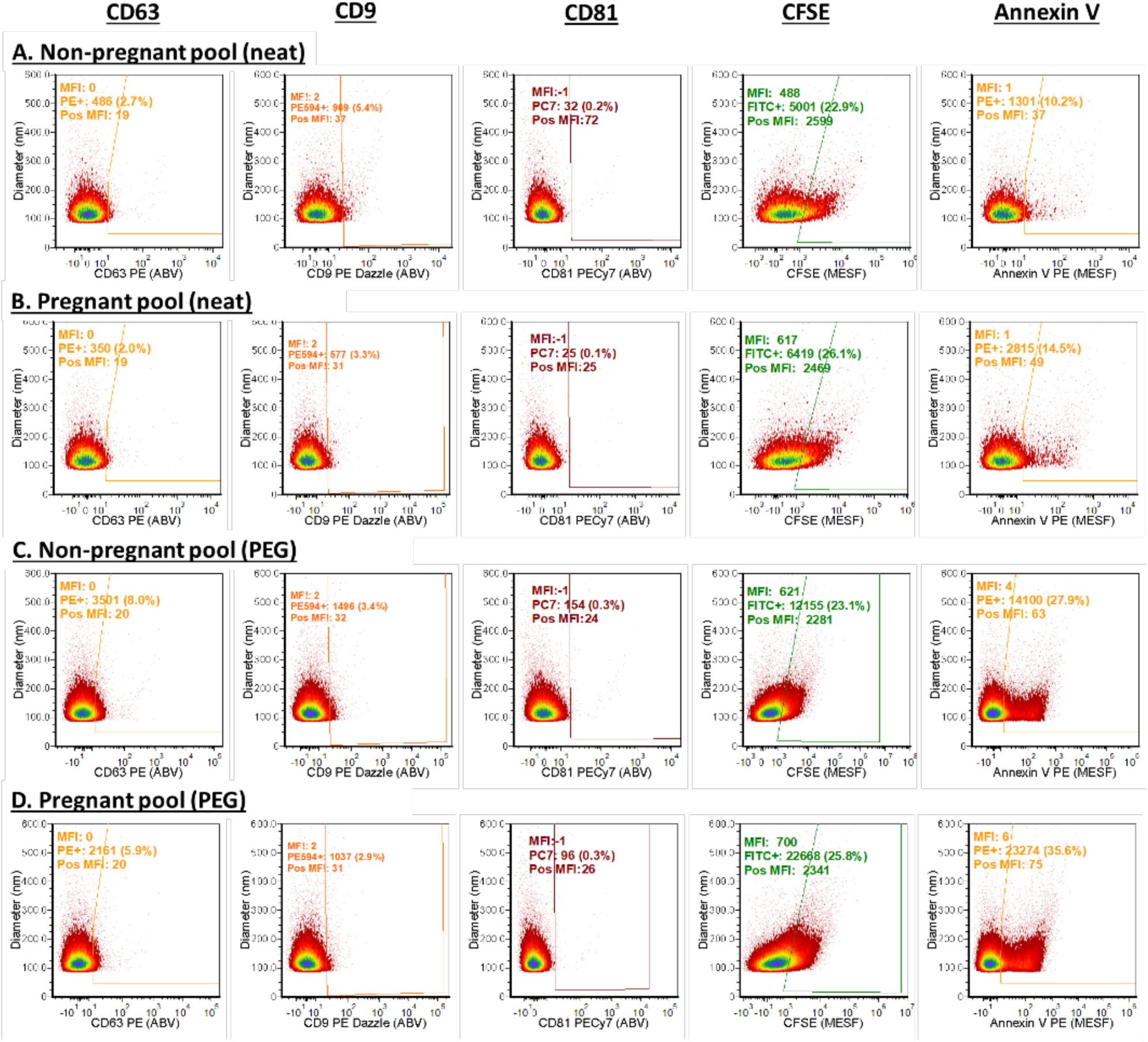
Vesicle Flow Cytometry of pregnant and non-pregnant female urine pools (neat and miRCURY PEG-precipitated). Expression of tetraspanins CD63, CD9, and CD81, along with CFSE and annexin V staining in **(A)** Non-pregnant neat urine pool, **(B)** pregnant neat urine pool, **(C)** mIRCURY (PEG)-precipitated non-pregnant urine pool, and **(D)** miRCURY (PEG)-precipitated pregnant urine pool samples.

## 4. DISCUSSION

Over the past decade, there have been advances in developing methods for isolation and characterization of uEVs and exRNA in liquid biopsies for the purpose of biomarker discovery in various disorders, especially those involving pathology of the kidney (25, 30, 31). Comparisons among previously published papers in this area are confounded by variability in urine sample collection and storage methods (e.g., time of collection, centrifugation, filtering, storage temperature, freeze/thaw cycles), uEV and exRNA isolation methods, and exRNA analysis methods (15). Transcriptomic profiling studies conducted using urine have largely focused on extracellular small RNAs, with a few studies profiling only long RNAs or both small and long RNAs (Table 5). The studies summarized in Table 5 focus on those that have utilized commercial kits for library preparation and Illumina platforms for RNA-Seq to enable direct comparison to the results in our study. For this reason, three studies that utilized “homebrew” methods of cDNA library preparation (32–34) and one study that utilized Ion Torrent sequencing (35) have been excluded from this summary of current literature. Most of the studies to date have profiled exRNA in urine from male donors, half of the study cohorts have included non-pregnant women, and none have profiled exRNA in urine during pregnancy (Table 5). There is therefore a great need for studies focused on non-pregnant and pregnant female cohorts to create reference profiles and develop new methods for non-invasive analysis of biofluids for identification of biomarkers of pregnancy disorders. We describe herein a comparative evaluation of five methods of urinary exRNA isolation and two methods of extracellular long RNA library preparation using reference urine pools created from non-pregnant and pregnant females as well as individual urine samples (Table 1). ExRNA isolation methods applied in this study included uEV isolation using either PEG-based precipitation (miRCURY and SeraMir) or membrane affinity (ExoRNeasy) to enrich for uEVs followed by exRNA extraction, or direct lysis of neat urine followed by spin column chromatography for total exRNA isolation (Norgen and miRNeasy advanced/mAdv).

**Table 5:**
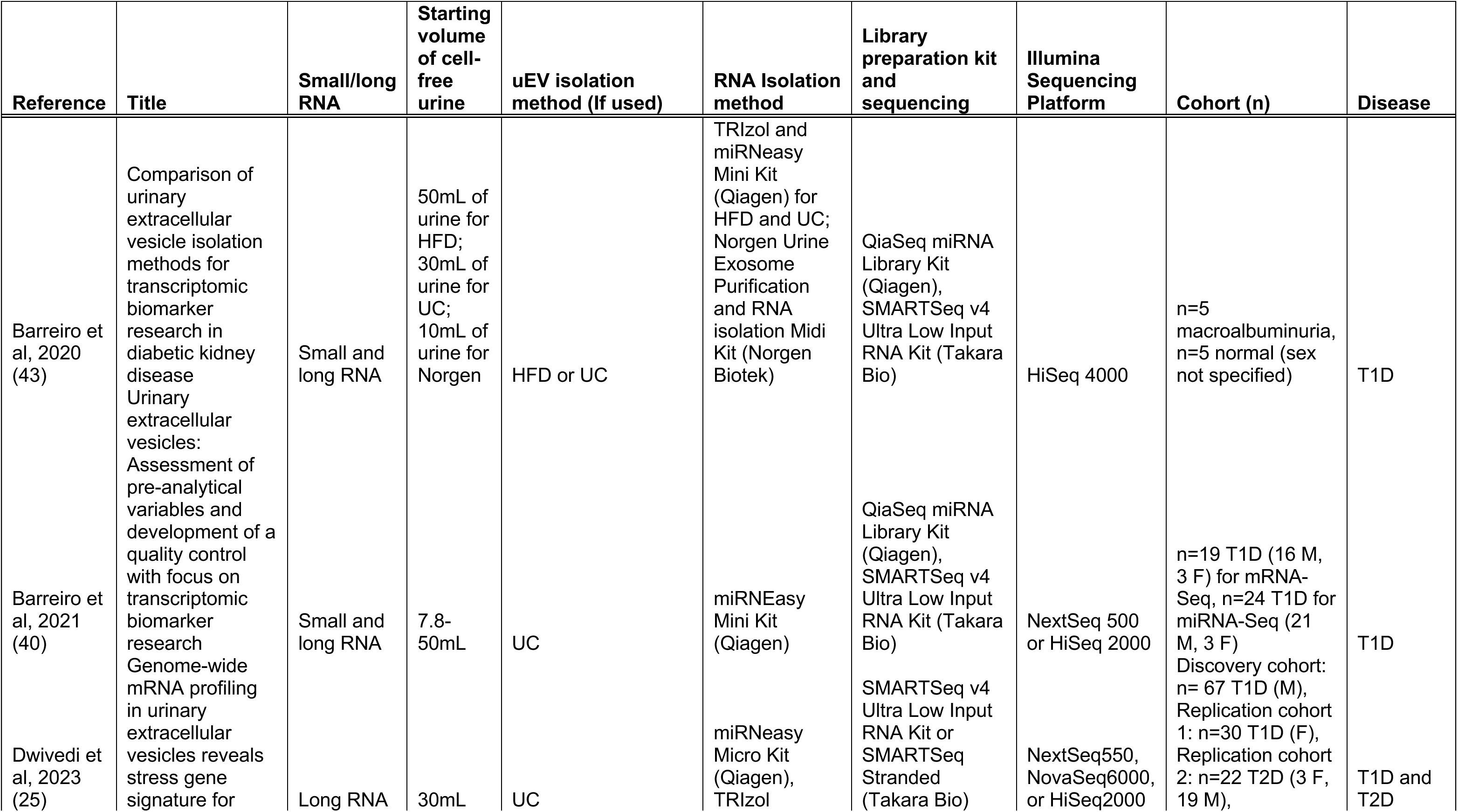
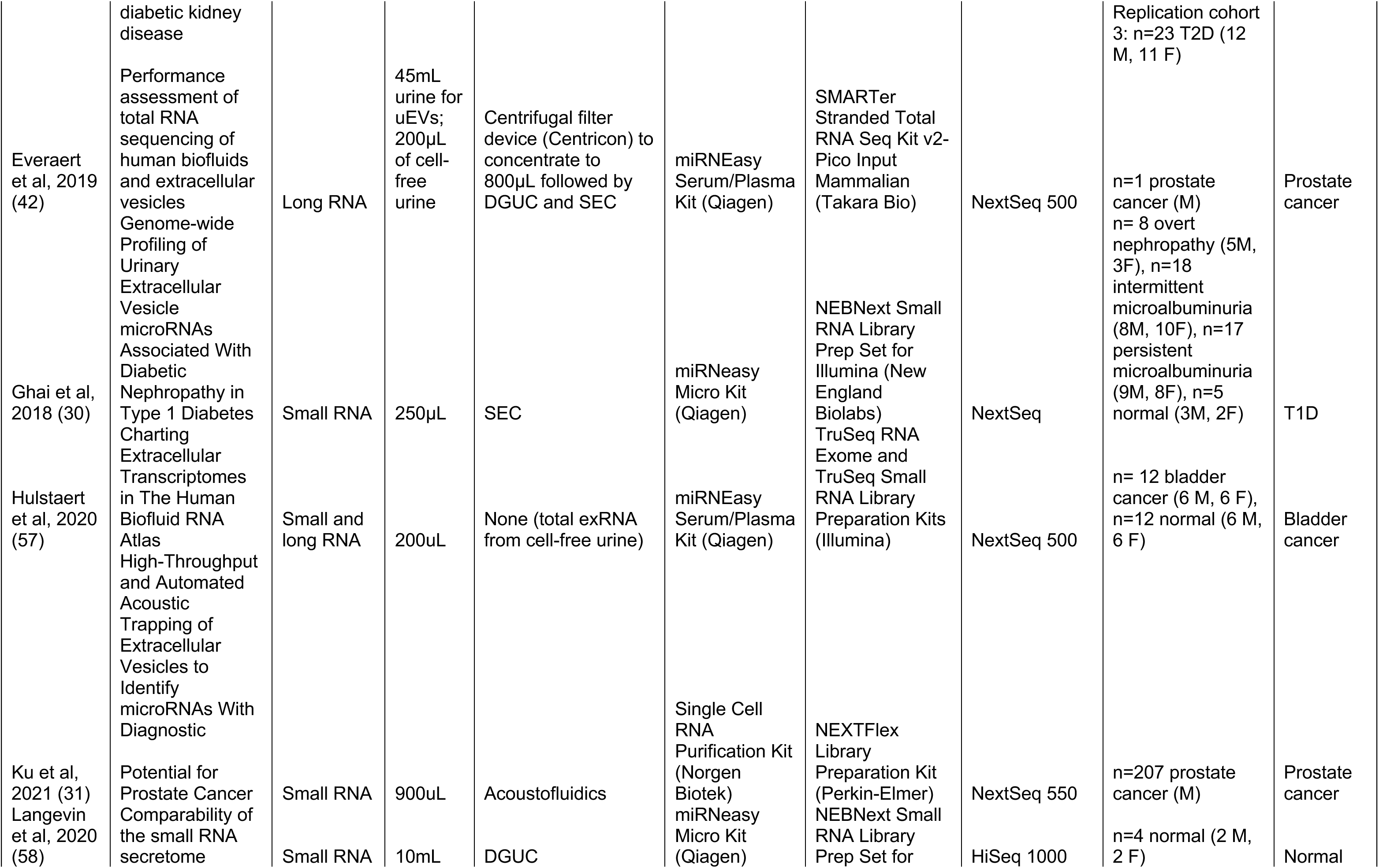
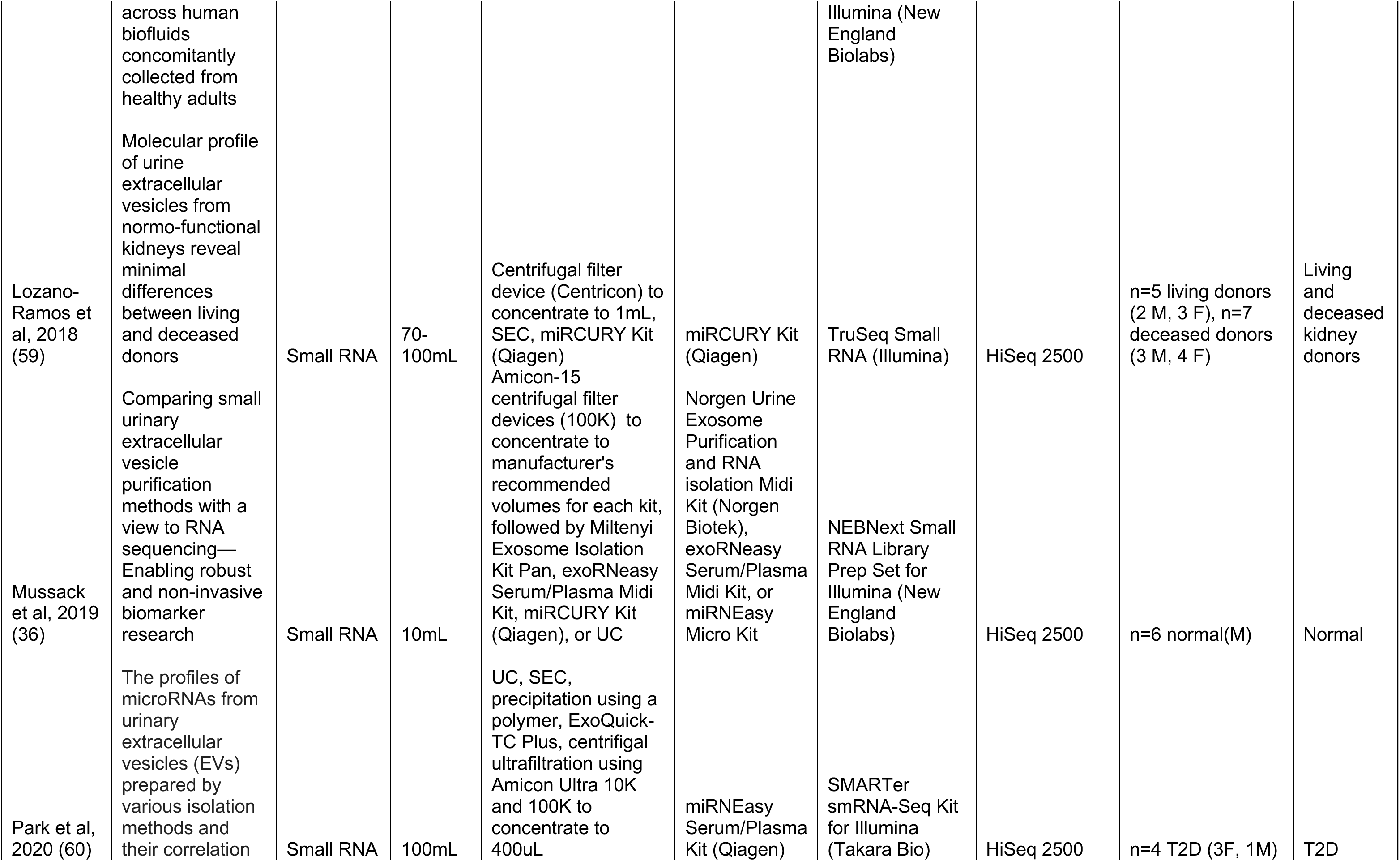
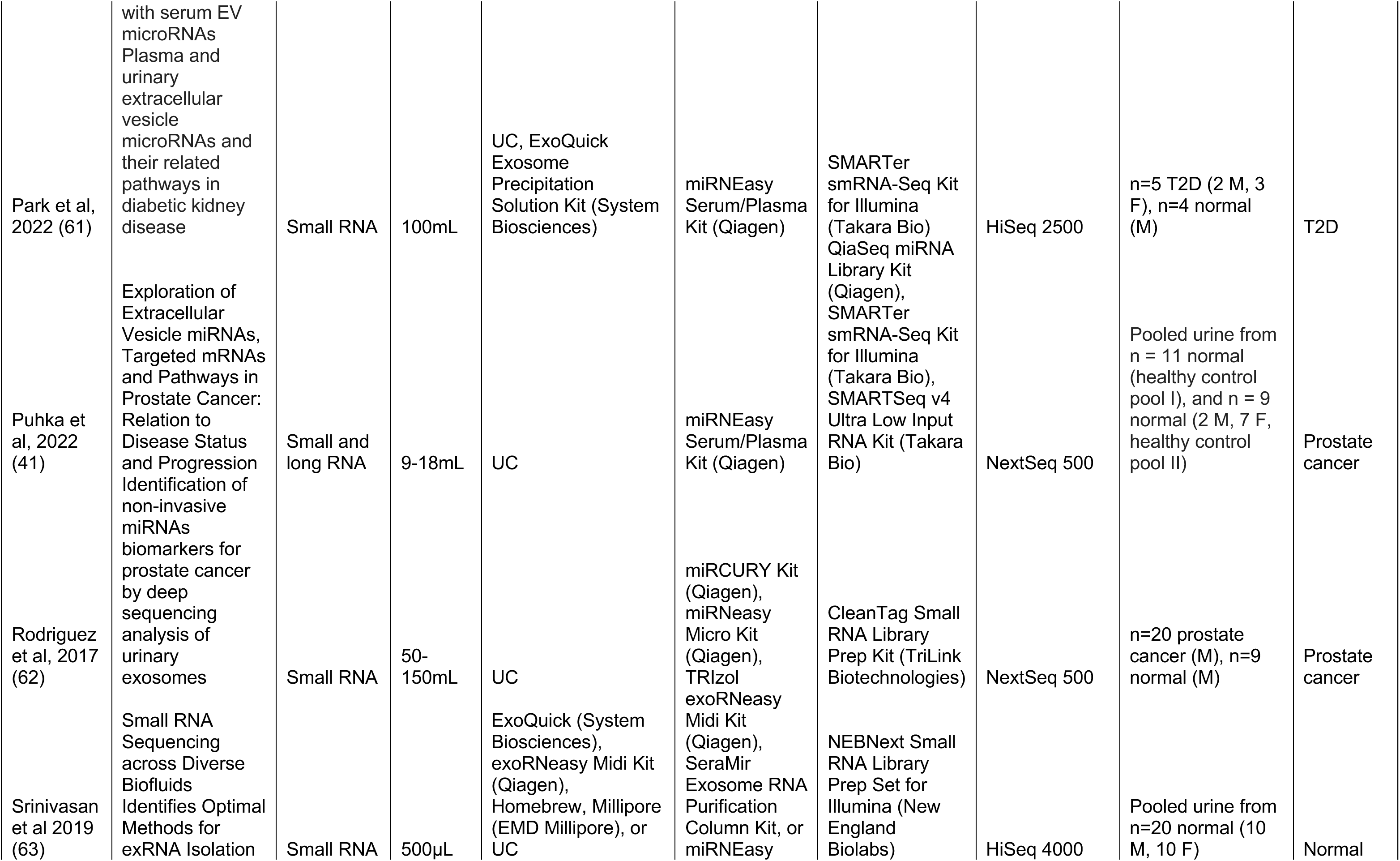
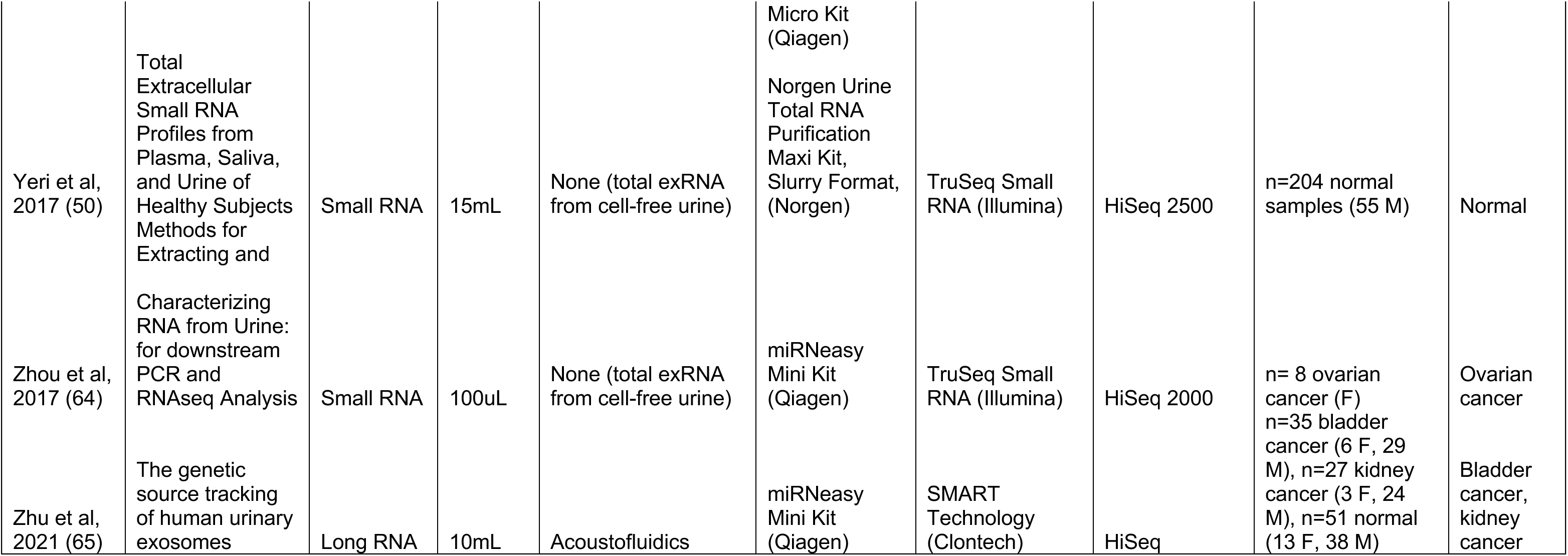
Summary of studies that have conducted RNA-Seq of urine samples using commercial library preparation kits and Illumina sequencing platforms.

Different exRNA isolation methods across plasma, serum, bile, and urine identify differential small RNA transcriptomic profiles which vary in RNA biotype and complexity or diversity across biofluids (2). Our findings reinforce this observation in female urine samples. Principal component analysis in our study showed clustering of miRNA profiles from female urine pools and individual samples by exRNA method, with distinct clusters of miRCURY and mAdv methods, and some overlap amongst samples extracted using the ExoRNeasy, Norgen, and SeraMir methods (Figure 3A). Separation was also observed between the urine reference pools and samples obtained from individuals (Figure 3C). We sought to identify miRNAs that are reproducibly identified in female urine pools and individual samples by all exRNA isolation methods, highlighting their potential utility for biomarkers. Mussack *et al.* identified 80 overlapping miRNAs amongst all exRNA isolation methods in male urine using a starting volume of 10 mL, filtered for miRNAs with ≥ 20 DESeq2 normalized reads per group (36). In our study, only 16 miRNAs overlapped amongst all exRNA isolation methods used to extract female pregnant and non-pregnant urine. This lower number of overlapping miRNAs amongst methods may be due to higher variability in the extracellular miRNA profile of female compared to male urine samples, differences in pregnant vs. non-pregnant urine, and the more stringent data filtering methods applied to identify pass-filter miRNAs detected by each kit in the current study (Supplementary Table 3). We applied a stringent filtering cutoff for samples extracted by each kit by calculating the value for 5 raw counts ((5 ÷ total miRNA read counts of lowest complexity sample) x 10^6^) in at least half of all the samples. This method of calculating cpm cutoffs requires that a miRNA be robustly expressed in at least half of the samples from a given kit in order to be considered detected using that kit (cpm cutoffs for each analysis are summarized in Supplementary Table 3). Out of the 16 overlapping miRNAs identified in this study, 12 have entries in Vesiclepedia and have been identified as abundant in urine following different methods of uEV and exRNA isolation, summarized in Table 2 (30, 35). These 16 miRNAs are therefore robust candidates for further study as biomarkers in urine in disease states due to their consistency, replicability and abundance across isolation methods and sex in different studies. In fact, among these 16 miRNAs, hsa-miR-30a-5p has been identified as a urinary biomarker in pediatric nephrotic syndrome (37), hsa-miR-21-5p as a biomarker in the urine of patients with prostate cancer (38), and hsa-miR-423-5p is increased in urine of patients with acute kidney injury (39).

When profiling exRNA, libraries should ideally be sequenced until a plateau in complexity is reached (2). For small RNA libraries in our study, the reduction in library preparation volumes to 1/5^th^ the manufacturer’s recommendation to allow for high-throughput processing of urine exRNA samples did not affect the complexity of samples as they are comparable to other studies (summarized in Table 5) which used higher starting volumes of urine and full volume library preparation reactions. In our study, the plateau in miRNA sequencing depth was reached for libraries from the miRCURY, mAdv, and Norgen methods, which displayed an average complexity of ∼152, ∼70, and ∼138 respectively in our study. Mussack *et al.* identified up to 124 miRNAs in urine samples from 6 healthy males across different exRNA isolation methods from 10 mL of urine, including the Norgen Urine miRNA Purification Kit (Slurry format) extraction kit (which is distinct from the kit used in this study), Miltenyi, miRCURY, ExoRNeasy kits and ultracentrifugation with density gradient (36). Similarly, we generated long RNA libraries using the SMARTSeq v4 kit with comparable mRNA complexity and read depth as studies utilizing higher starting volumes of urine and full volume library preparation methods. The average complexity of libraries prepared using the SMARTSeq v4 kit was significantly higher than the Pico v2 kit (Supplementary Figure S3A). The difference in performance between the Pico v2 and SMARTSeq v4 kits has been attributed to the different methods of priming for the initial reverse transcription step. The Pico v2 kit uses random priming, results in a higher proportion of intergenic reads (40), while the SMARTSeq v4 kit uses oligo-dT primers and thus targets poly-A tailed mRNAs (40, 41). Previous RNA-Seq analysis of cell-free urine and uEV isolates have identified sequences from >10,000 protein-coding genes in addition to non-coding RNA and abundant rRNA species (15, 42, 43). A recent study by Dwivedi *et al.* identified expression of >10,000 mRNAs (25) in urine from a cohort with Type I diabetes using the SMARTSeq v4 kit; however the cutoff used was a count of ≥1, while the cutoff we have applied in this study to identify the number of unique mRNAs is ≥5 raw counts and is therefore more stringent. We obtained a comparable number of average total mRNA reads in samples isolated from 4 mL of urine using Norgen (for plasma) and 1/5^th^ volume library preparation reactions with SMARTSeq v4 (∼1.19 x 10^6^) as that reported by Barreiro *et al.* from 10 mL starting volume of urine extracted using the Norgen method (for urine) and full volume SMARTSeq v4 reactions for library preparation (∼2.1 x 10^6^) (43). Barreiro *et al.* report a higher number of mRNA reads from hydrostatic filtration dialysis from 50 mL (∼7.8 x 10^6^) and ultracentrifugation using 30 mL of urine (∼8.4 x 10^6^). Everaert *et al.* identified >10,000 unique mRNAs from 45 mL starting volume of urine which was concentrated and uEVs were isolated using size exclusion chromatography. This type of pre-processing is a comprehensive and time-consuming process which is not scalable for high-throughput clinical biomarker studies. Our study has therefore detected a comparable number of unique protein-coding mRNAs expressed with at least ≥5 raw counts and comparable read depth using 1-4 mL neat, unconcentrated cell-free urine as the starting volume and scalable methods of library preparation using the SMARTSeq v4 kit as studies that have employed full volume library preparation reactions from ≥10 mL starting volume of urine.

We are particularly interested in methods for robust miRNA and mRNA profiling in urine in pregnancy in order to identify biomarkers and decipher the pathology of pregnancy disorders such as preeclampsia in larger clinical studies. In this study, we defined the ‘optimal’ method as the method that yields the highest percentage of miRNA and uniquely mapped reads in small and long RNA profiling respectively, as well as the highest complexity (individual RNAs with ≥5 raw counts in each sample), and ability to identify differentially expressed RNAs between different biological groups. Using these metrics, the miRCURY method using 4 mL starting volume of neat, unconcentrated cell-free urine appeared to perform the best across exRNA isolation methods. Samples extracted from 4 mL of urine using the Norgen (for plasma) method in this study yielded miRNA and mRNA complexity that was not significantly different from samples extracted using 4 mL of urine using the miRCURY method (Figure 2D, Figure 6D). However, we were unable to identify differentially expressed RNAs using the Norgen method between biological groups compared in this study (data not shown). The miRCURY method includes an initial uEV precipitation step followed by lysis and an RNA-binding column. The Norgen method involves direct lysis of the biofluid followed by addition of RNA-binding slurry, and yields not only EV-associated RNA, but also exRNA associated with ribonucleoproteins and lipoproteins. The miRCURY method may therefore precipitate uEVs containing RNAs with lower expression that may be biologically relevant to disease processes. This is reinforced by two observations: (1) Analysis of the coefficient of variation (% CV) across exRNA isolation methods using miRNA expression data showed that the Norgen method is able to profile highly expressed miRNAs (>100 cpm) with less variability across samples, while the miRCURY method is able to detect miRNAs expressed between 10-100 cpm (Supplementary Figure S2), and (2) Pathways analysis of mRNAs enriched in miRCURY-extracted samples highlighted that mRNAs for “GO: Extracellular Exosome” are enriched in 2^nd^ and 3^rd^ trimester pregnant urine (Table 4). This included upregulation of mRNA for *DICER1* which codes for Dicer, a cytoplasmic endoribonuclease that is essential for the cleavage and maturation of pre-miRNAs, and loads small RNAs into Argonaute, part of the RNA-induced silencing complex (RISC) which degrades mRNA (44).

We used the miRCURY method to profile an expanded number of female non-pregnant and pregnant urine pools and individual samples using 1 mL and 4 mL starting volumes of urine, and observed that hsa-miR-516b-5p, which forms part of the C19MC miRNA cluster that is highly expressed in anthropoid primate placentas (45) is significantly upregulated in 3^rd^ trimester urine samples from pregnant individuals compared to 2^nd^ trimester pregnant urine pools. Urine from pregnancy was enriched for five miRNAs that are highly expressed in the placenta compared to other cell/tissue types such as the lungs, liver, and kidney (29): hsa-miR-125a-5p,-30d-5p,-100-5p,-99b-5p,-127-3p, and-516b-5p (Table 3). Out of these, hsa-miR125a-5p is significantly increased in placental tissue from preeclamptic pregnancies (46), hsa-miR-30d-5p is significantly reduced in placentas from patients with gestational diabetes mellitus (GDM) (47), hsa-miR-100-5p showed a trend of down-regulation in intrauterine growth restriction placentas requiring delivery before 34 weeks (48), and hsa-miR-127-3p is decreased two-fold in placentas from preeclampsia (49). However, it is unknown whether decreased expression of miRNAs in the placenta is reflected as an increase in those miRNAs in urine. More research is required to track the mechanism through which potentially placenta-derived miRNAs may be transported into maternal urine. Deconvolution analysis using miRNA expression data from various cell and tissue types (29) demonstrated that most miRNAs in pregnant urine from pools and individuals extracted using the miRCURY method are likely derived from the kidney, with a small fraction originating from the placenta. Since the contribution of the placenta to maternal urine pools in deconvolution analysis was relatively low compared to other tissue sources such as the kidney, maternal urine may be more comprehensively utilized to identify changes in RNAs reflecting kidney function in pregnancy disorders involving the kidney such as preeclampsia.

In selecting a method for urinary exRNA isolation and library preparation to be used for analysis of clinical cohorts in future, investigators should consider the small and long RNA biotype of interest and the optimal exRNA isolation method for that biotype of interest. For example, in this study the Norgen and ExoRNeasy kits yielded significantly higher percentages of tRNA compared to other methods amongst small RNA libraries. A previous study using the Norgen kit (for urine) showed significantly higher fractions of certain tRNA fragments, particularly tRNA^Gly^ (GlyGCC) which constituted 86% of detected tRNA fragments in urine compared to plasma and saliva (50). This was replicable in female urine samples using the Norgen kit for plasma/serum in this study, as tRNA^Gly^ constituted ∼91% of detected tRNA fragments in Norgen-extracted pregnant and non-pregnant female urine pools from 4 mL starting volume (data not shown). The high proportion of tRNA fragments may be related to the selective secretion of specific RNA biotypes into urine, or proximity to an organ with high expression levels of this tRNA or fragment (50).

In addition to transcriptomic analysis, we also conducted quantification and characterization of nanoparticles in cell-free non-pregnant female urine pools prior to enrichment of EVs and/or exRNA isolation using NTA and MRPS. NTA is a commonly used method to measure particle concentration and size distribution in biofluids (15). NTA may be less sensitive in detecting particles <70 nm in diameter in urine (51), and in our analysis NTA detected fewer particles/μL than MRPS in non-pregnant female urine. Further, lack of disruption of the large majority of NTA events following addition of detergent may indicate that NTA measurements are confounded by protein aggregates, such as uromodulin (15). Nearly all events detected by MRPS were susceptible to detergent and may represent smaller EVs not detected by NTA. Both NTA and MRPS estimated the median size range of uEVs to be between ∼100nm in diameter. Direct analysis of cell-free urine prior to EV enrichment has utility for high-throughput clinical implementation (15), and here we demonstrate methods for dilution and analysis of neat female non-pregnant urine pools prior to exRNA isolation methods for transcriptomic analysis. Vesicle flow cytometry (vFC) is a more precise method for characterization of smaller uEVs (15), which may be the predominant EV subpopulation in urine (26). Nano-flow cytometry (52) and imaging flow cytometry (53) are novel techniques that are being developed for single uEV analysis in cell-depleted urine (15). Few studies analyzing uEVs using high resolution vFC exist, as such analysis requires specialized cytometers with high scatter sensitivity (15). We conducted vFC using non-pregnant and pregnant female urine reference pools, as well as pools that were enriched for EVs using the miRCURY PEG-based precipitation buffer, since this was the method we used to perform the largest number of exRNA isolations in the current study. Interestingly, we demonstrate that uEVs detected using vFC demonstrate low tetraspanin expression in non-pregnant and pregnant urine pools, both with and without enrichment using miRCURY. Amongst, the tetraspanins, CD9 was the most commonly expressed in urine samples analyzed in this study (2.9-5.4% of EVs). Notably, CD9 is reportedly abundant on small uEVs detected using imaging flow cytometry compared to CD63 and CD81 (54). Despite the relatively low expression of tetraspanins detected using vFC, we note that the events detected by vFC were vesicular, since they were disrupted by the addition of detergent (Supplementary Methods Figure 4 A). Precipitation with PEG-based buffer using the miRCURY method resulted in enrichment of the compartment of uEVs that were positive for annexin V, potentially representing exosomes and microvesicles (100-1000nm). Our vFC analysis of urine was conducted strictly following MIFlowCyt-EV guidelines (55) and parameters are reported in Supplementary Methods.

Our study has several limitations. First, we cannot exclude the possibility that extracted RNA samples contain non-EV RNAs, as we did not pre-treat our samples with RNase prior to extraction and were focused on developing a reproducible approach to urinary exRNA extraction and library preparation rather than focusing on specific compartments associated with exRNAs. Second, we did not conduct in-depth analysis of lncRNAs or circRNAs or include these studies in our summary of the literature on urinary exRNA, as we focused on urinary miRNAs and mRNAs. We note that one study has reported that lncRNAs are the predominant RNA subtype in urine (34), but this could not be assessed in our data because the size selection parameters we used for small RNA-Seq library preparation focused on RNA molecules 15-30 nt in length. Third, we did not account for changes in urine concentration during pregnancy, which may be reflected in the large number of down-regulated genes in the 3^rd^ trimester compared to the 2^nd^ trimester (Supplementary Table 7). Efforts to normalize urine RNA-Seq data accounting for intra-and inter-individual variability of this biofluid due to circadian rhythm, changes in salt and water homeostasis, and across disease states are ongoing, and measurement of urine specific gravity, total protein, or total exRNA concentration, though not routinely used, may be applied in clinical studies in the future (56). Finally, the exRNA isolation methods presented here are manual. To be truly scalable for clinical studies, it is essential to optimize automated methods for urinary exRNA isolation in addition to automated library preparation methods.

In summary, we have outlined in this study an evaluation of five exRNA isolation methods and two library preparation methods for miniaturized extracellular long RNA library preparation. Our analyses indicate that the miRCURY exRNA isolation method with 4 mL starting volume of neat, unconcentrated cell-free urine, combined with 1/5^th^ reaction volumes for library preparation yield comparable small (NEB Next kit) and long (SMARTSeq v4 Ultra Low Input kit) RNA libraries and RNA-Seq data compared to previously published studies with higher starting volumes and more steps to pre-process urine. By incorporating miniaturization and automation of the small and long RNA library preparation workflows, which increase throughput and decrease cost, our results demonstrate the feasibility of future large-scale studies that involve large numbers of urine samples with modest starting volumes and minimal pre-processing before exRNA isolation. Standard processes for small and long RNA library preparation for RNA-Seq are lengthy and expensive. A reduction in library preparation volumes enables cost-effective exRNA sequencing of large numbers of samples, enabling biomarker discovery and validation in large clinical cohorts.

## Supporting information

Supplementary vFC methods

## Acknowledgements

We would like to thank Louise C. Laurent (LCL) MD, PhD for her insightful comments on this manuscript. This work was supported by funds from the National Institutes of Health: UG3CA241687 (National Cancer Institute) to LCL (Co-I: PP) and R00HD096125 (*Eunice Kennedy Shriver* National Institute of Child Health and Human Development) to PP. This publication includes data generated at the UC San Diego IGM Genomics Center utilizing an Illumina NovaSeq 6000 that was purchased with funding from a National Institutes of Health SIG grant (#S10 OD026929). Data management, storage, and analysis were performed using the Extreme Science and Engineering Discovery Environment (XSEDE) Comet at the San Diego Supercomputing Center through allocation MCB140074, as well as the Genboree exceRpt Small RNA-Seq pipeline. We would like to thank the Clinical Research Coordinators at the Center for Perinatal Discovery, UC San Diego, for their help in recruiting patients and collecting samples, and the patients for donating their samples to this study. We would also like to thank Mariko Horii (MD) for her help in identifying the clinical data for the cohort of pregnant women included in this study.

## Author contributions

Conceptualization, SS, DEW, JPN, PP. Methodology, SS, PDH, JPN, PP. Formal Analysis, SS, REM, PDH, JPN, PP. Investigation: SS, PDH, ALM, AV, MM, MM, ED, JPN, PP. Data Curation: SS, JPN, PP. Writing-Original Draft: SS, PP. Visualization: SS, JPN, PP. Supervision: DEW, JPN, PP. Funding Acquisition: LCL and PP.

