## Supplementary figures and tables for "Miniaturized Workflow for Transcriptomic Profiling of Urinary Extracellular RNA during Pregnancy"

### Miniaturized Workflow for Transcriptomic Profiling of Urinary Extracellular RNA during Pregnancy: Supplementary Figures and Tables

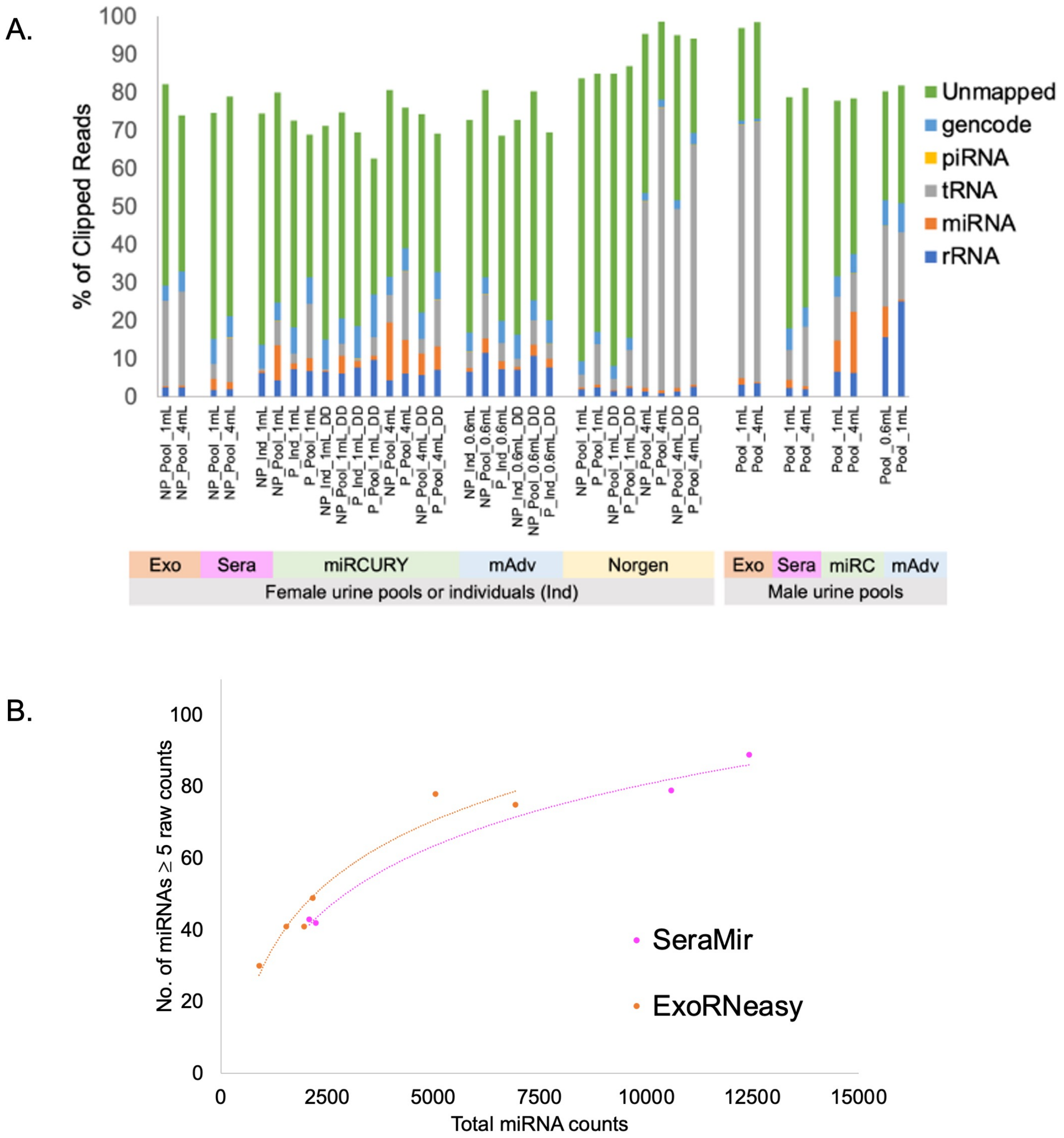

**Figure S1: Small RNA biotypes and complexity by exRNA isolation method for all sample types. (A)** Small RNA biotypes by exRNA isolation method expressed as a percentage of total clipped reads, split into groups according to pregnant (P), non-pregnant (NP) pools or individuals with or without dry-down (DD) from female (F) and male (M) samples with starting volumes of either 1mL or 4mL of urine (total n=208 small RNA libraries), processed using the Genboree exceRpt small RNA pipeline. **(B)** Complexity (individual miRNAs with ≥ 5 raw counts in each sample) split by extraction method for SeraMir (Sera) and ExoRNeasy (Exo)-extracted samples showing lower complexity and total miRNA read counts compared to miRCURY, miRNeasy Advanced, and Norgen methods.

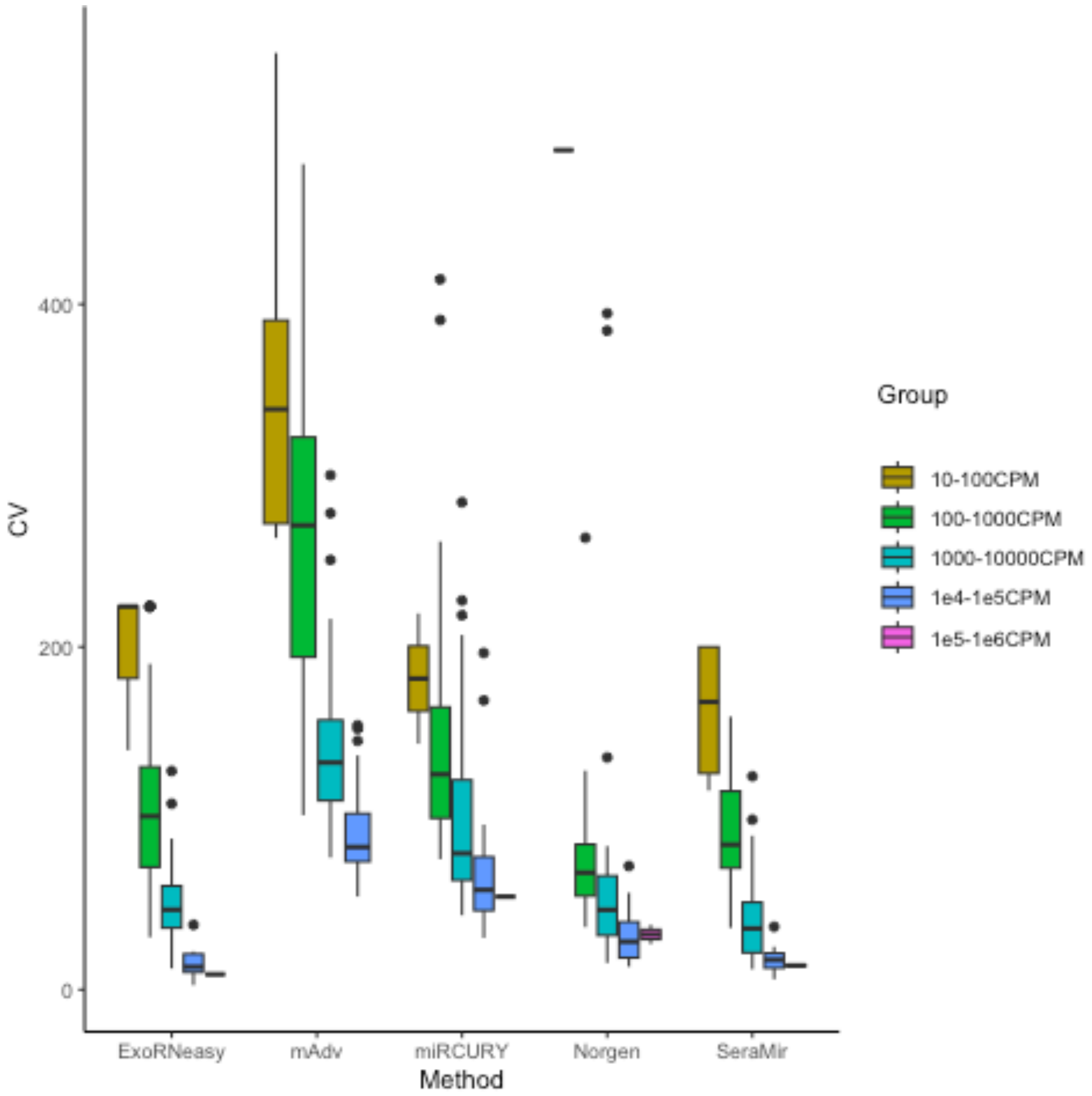

**Figure S2: Coefficient of variation (% CV) as a function of miRNA expression levels in counts per million (CPM).** %CV decreased with increasing miRNA expression for all exRNA isolation methods, indicating that miRNAs with lower expression showed higher variability in the samples across exRNA isolation methods.

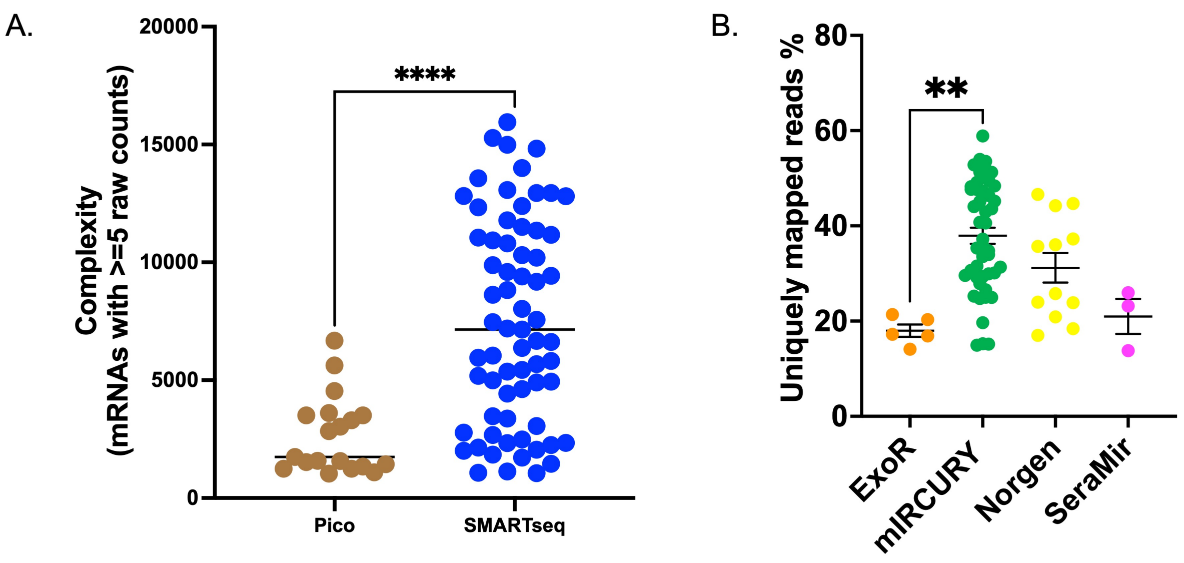

**Figure S3**: **(A)** Complexity (individual mRNAs with ≥ 5 raw counts in each sample) split by library preparation method for long RNA libraries showing significantly higher complexity of libraries prepared using SMARTSeq v4 compared to Pico v2 ($p<0.05$). **(B)** Percentage of uniquely mapped reads split by exRNA isolation method for long RNA libraries prepared using the SMARTSeq v4 kit showing significantly higher uniquely mapped reads for samples isolated using miRCURY compared to ExoRNeasy ($p<0.05$).

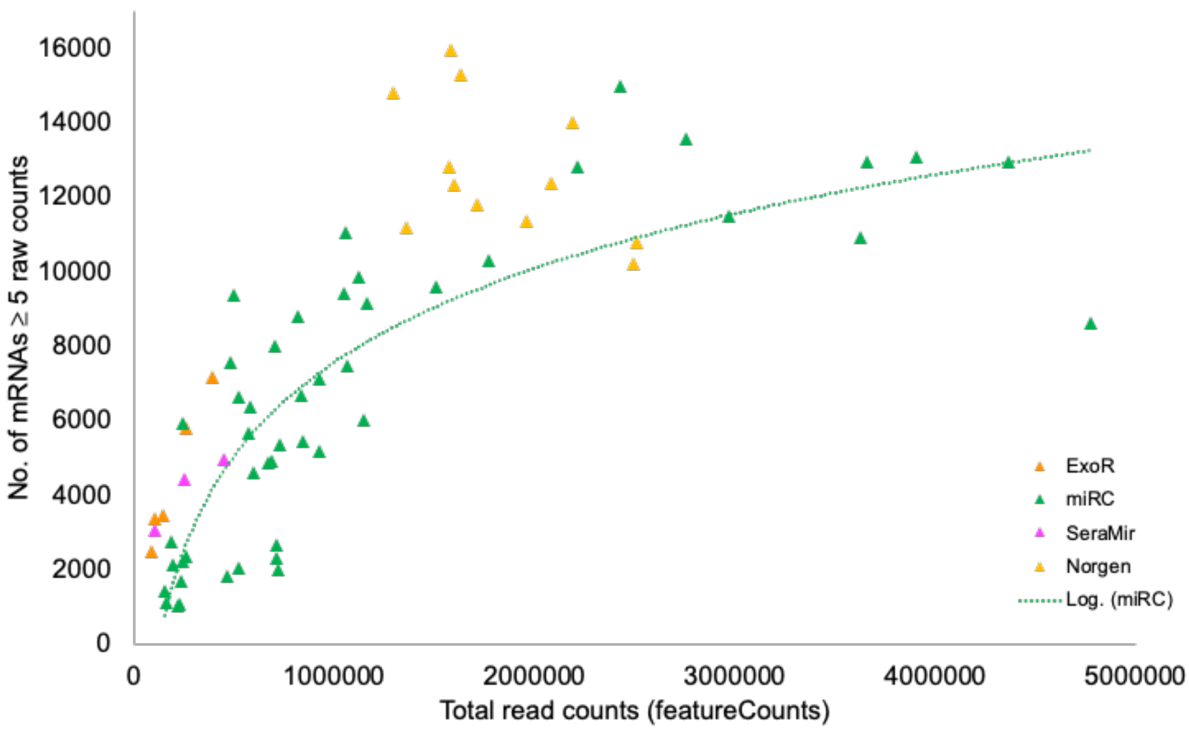

**Figure S4:** Complexity (individual mRNAs with ≥ 5 raw counts in each sample) split by exRNA isolation method for samples prepared using the SMARTSeq v4 kit.

**Supplementary Table 1**: Clinical information of patients from which urine samples were used to create reference pools and individual samples.

| Sex | Group | Type | Sample name | Age | | |
|---|---|---|---|---|---|---|
| F | Non-pregnant female | Pool | NP_pool | 20 | | |
| F | Non-pregnant female | Pool | | 19 | | |
| F | Non-pregnant female | Pool | | 19 | | |
| F | Non-pregnant female | Pool | | 19 | | |
| F | Non-pregnant female | Pool | | 20 | | |
| F | Non-pregnant female | Pool | | 19 | | |
| F | Non-pregnant female | Pool | | 19 | | |
| F | Non-pregnant female | Pool | | 21 | | |
| F | Non-pregnant female | Pool | | 21 | | |
| F | Non-pregnant female | Pool | | 20 | | |
| M | Male | Pool | M_pool | 19 | | |
| M | Male | Pool | | 23 | | |
| M | Male | Pool | | 25 | | |
| M | Male | Pool | | 21 | | |
| M | Male | Pool | | 19 | | |
| M | Male | Pool | | 32 | | |
| M | Male | Pool | | 22 | | |
| M | Male | Pool | | 20 | | |
| M | Male | Pool | | 21 | | |
| M | Male | Pool | | 21 | | |
| F | Non-pregnant female | Individual | NP1 | 20 | | |
| F | Non-pregnant female | Individual | NP2 | 19 | | |
| F | Non-pregnant female | Individual | NP3 | 19 | | |
| F | Non-pregnant female | Individual | NP4 | 19 | | |
| F | Non-pregnant female | Individual | NP5 | 19 | **Gestational age (weeks.days)** | **First BMI (kg/m^2)** |
| F | Pregnant female | Individual | P1 | 24 | 32.2 | 37.4 |
| F | Pregnant female | Individual | P2 | 29 | 34.2 | 21.3 |
| F | Pregnant female | Individual | P3 | 37 | 34.6 | 24.2 |
| F | Pregnant female | Individual | P4 | 26 | 34 | 19.4 |
| F | Pregnant female | Individual | P5 | 30 | 36.4 | 31.1 |
| F | Pregnant female | Pool | P_pool | 32 | 27.6 | 21.5 |
| F | Pregnant female | Pool | | 37 | 20.2 | Not available |
| F | Pregnant female | Pool | | 34 | 28.3 | 17.6 |
| F | Pregnant female | Pool | | 30 | 27.3 | 26.9 |
| F | Pregnant female | Pool | | 33 | 24.2 | 24.1 |
| F | Pregnant female | Pool | | 27 | 22.3 | 22.7 |
| F | Pregnant female | Pool | | 35 | 21.4 | 24.1 |
| F | Pregnant female | Pool | | 38 | 19.1 | Not available |
| F | Pregnant female | Pool | | 32 | 24.3 | 41 |
| F | Pregnant female | Pool | | 36 | 25.2 | 38.3 |

**Supplementary Table 2:** Statistical analyses conducted to compare biotypes amongst small RNA libraries.

| Comparison of %miRNA in F only samples split by dry-down (DD) or without dry-down | | | |
|---|---|---|---|
| **All F samples** | **Number of values** | **Mean** | **Std. Error of Mean** |
| ExoR | 6 | 0.3 | 0.03334 |
| SeraMir | 4 | 2.7 | 0.59 |
| miRC | 48 | **4.2** | 0.8263 |
| miRC_DD | 45 | 2.1 | 0.4478 |
| mAdv | 30 | 1.7 | 0.5622 |
| mAdv_DD | 32 | 1.7 | 0.4213 |
| Norgen | 12 | 0.7 | 0.0636 |
| Norgen_DD | 12 | 0.6 | 0.0748 |
| **Dunn's multiple comparisons test** | **Adjusted P Value** | | |
| miRCv vs. mAdv | 0.011 | | |
| **Comparison of %miRNA in miRCURY-extracted samples without dry-down, split by pool, individual, and volume** | | | |
| **miRCURY F samples without DD** | **Number of values** | **Mean** | **Std. Error of Mean** |
| NP_Pool_1mL | 8 | 9.2 | 1.335 |
| NP_Ind_1mL | 15 | 0.5 | 0.1743 |
| NP_Pool_4mL | 4 | **15.2** | 5.321 |
| P_Pool_1mL | 3 | 3.3 | 0.6919 |
| P_Ind_1mL | 15 | 1.7 | 0.4311 |
| P_Pool_4mL | 3 | **8.8** | 0.7228 |
| **Dunn's multiple comparisons test** | **Adjusted P Value** | | |
| NP_Pool_1mL vs. NP_Ind_1mL | <0.0001 | | |
| NP_Pool_1mL vs. P_Ind_1mL | 0.0411 | | |
| NP_Ind_1mL vs. NP_Pool_4mL | 0.0003 | | |
| NP_Ind_1mL vs. P_Pool_4mL | 0.0142 | | |
| **Comparison of %tRNA in all F samples extracted without dry-down** | | | |
| **All F samples without DD** | **Number of values** | **Mean** | **Std. Error of Mean** |
| ExoRNeasy | 6 | 23.6 | 0.5375 |
| miRCURY | 48 | 4.7 | 0.7852 |
| mAdv | 30 | 5.2 | 1.547 |
| Norgen | 12 | **34.3** | 8.725 |
| SeraMir | 4 | 5.8 | 2.099 |
| **Dunn's multiple comparisons test** | **Adjusted P Value** | | |
| ExoRNeasy vs. miRCURY | 0.0021 | | |
| ExoRNeasy vs. mAdv | 0.0044 | | |
| miRCURY vs. Norgen | 0.0006 | | |
| mAdv vs. Norgen | 0.0021 | | |

**Supplementary Table 3:** Summary of small and long RNA-Seq analyses along with cpm cutoff used in each comparison.

| Manuscript figure/Ta | Data | Samples | Purpose | Comparison | Sample type | n | n | Total miRNA read counts | RNAs | Package or type of analys | Filtering cutoff (cpm present in at least n) |
|---|---|---|---|---|---|---|---|---|---|---|---|
| Fig. 3 | Small RNA | All | Visualization | exRNA Isolation Kits for PCA analysis | Pools and individuals | 176 (F only) | | >1000 | 111 unique miRNAs, 16 overlap | Filtering using edgeR | ExoRNeasy: 2534cpm in at least 3 samples; |
| Fig. 4, Supp. Table 4 | | no DD | DE analysis | Total (Norgen) vs. Vesicular (miRCURY) | | | | | | | |
| | | | | | Non-pregnant pools | 12 miRCURY | 6 Norgen | >25,000 | 35 DE miRNAs (15↑ and 20↓ in vesicular exRNA isolated samples | TMM | 181cpm in at least 9 samples |
| | | | | | Pregnant 2nd trimester pools | 6 miRCURY | 6 Norgen | >25,000 | 48 DE miRNAs (25↑ and 23↓ in vesicular exRNA isolated samples | TMM | 170 cpm in at least 6 samples |
| Fig. 5, Supp. Table 5 | | no DD | DE analysis | miRCURY Pregnant vs. non-pregnant | | | | | | | |
| | | | | | Non-pregnant and 2nd trimester pregnant | 12 non-pregnant pools | 6 2nd trimester pregnant pools | >70,000 | 37 DE miRNAs (14↑ and 23↓ in 2nd trimester pregnant pools) | TMM | 67cpm in at least 9 samples |
| | | | | | Non-pregnant and 3rd trimester pregnant | 11 non-pregnant from 4 individuals | 14 3rd trimester pregnant from 5 individuals | >6000 | 6 DE miRNAs (3↑ and 3↓ in 3rd trimester pregnant individuals) | TMM | 640cpm in at least 12 samples |
| | | | | | Pregnant 2nd trimester pools and 3rd trim | 6 2nd trimester pregnant pools | 14 3rd trimester pregnant from 5 individuals | >20,000 | 17 DE miRNAs (7↑ and 10↓ in 3rd trimester pregnant individuals) | TMM | 146cpm in at least 10 samples |
| **Manuscript figure/Ta** | **Data** | **Samples** | **Purpose** | **Comparison** | **Sample type** | **n** | **n** | **Total mRNA read counts** | **RNAs** | **Package or type of analysis** | |
| Fig. 6 | Long RNA | no DD | Visualization | Library prep kits for PCA analysis | Pools and individuals | 19 Pico v2 | 67 SMARTSeq v4 | >20K | 1176 (Pico v2), 3738 (SMARTSeq) unique mRNAs, 455 overlap | Filtering using edgeR | Pico v2: 53 cpm in at least 10 samples, SMAR |
| Supp. Table 6 | | no DD | DE analysis | Total (Norgen) vs. Vesicular (miRCURY) SMARTSeq v4 | | | | | | | |
| | | | | | Non-pregnant pools | 14 miRCURY | 6 Norgen | >450K | 176 DE mRNAs (13↑ and 163↓ in vesicular exRNA isolated sampl | TMM, limma-voom | 10cpm in at least 10 samples |
| | | | | | Pregnant 2nd trimester pools | 6 miRCURY | 6 Norgen | >450K | 75 DE mRNAs (9↑ and 66↓ in vesicular exRNA isolated samples) | TMM, limma-voom | 3cpm in at least 6 samples |
| Supp. Table 7 | | no DD | DE analysis | miRCURY Pregnant vs. non-pregnant SMARTSeq v4 | | | | | | | |
| | | | | | Non-pregnant and 2nd trimester pregnant | 15 non-pregnant pools | 6 pregnant pools | >200K | 50 DE mRNAs (44↑ and 6↓ in 2nd trimester pregnant pools) | TMM, limma-voom | 21cpm in at least 11 samples |
| | | | | | Non-pregnant and 3rd trimester pregnant | 8 non-pregnant from 5 individuals | 10 samples from 4 individuals | >250K | 208 DE mRNAs (204↑ and 4↓ in 3rd trimester pregnant individual | TMM, limma-voom | 11cpm in at least 9 samples |
| | | | | | Pregnant 2nd trimester pools and 3rd trim | 6 non-pregnant pools | 10 samples from 4 individuals | >250K | 230 DE mRNAs (5↑ and 225↓ in 3rd trimester pregnant individuals | TMM, limma-voom | 8cpm in at least 8 samples |

Filtering cutoff calculated as (5/total readcounts of lowest complexity sample) x $10^6$ cpm present in at least 50% of samples in each analysis

**Supplementary Table 4:** MicroRNAs that were significantly enriched in vesicular (miRCURY) or total (Norgan) exRNA isolation methods that were unique to either pregnant or non-pregnant pools, as well as overlapping in both pools (FDR<0.05).

| miRNA | Log fold-change | FDR | Pool | Vesicular expression |
|---|---|---|---|---|
| **Enriched following vesicular exRNA isolation** | | | | |
| hsa-miR-192-5p | 0.57 | 0.0273 | Non-pregnant | ↑ |
| hsa-miR-363-3p | 0.82 | 0.0249 | Non-pregnant | ↑ |
| hsa-miR-92a-3p | 0.68 | 0.0344 | Non-pregnant | ↑ |
| hsa-miR-941 | 1.27 | 0.0206 | Non-pregnant | ↑ |
| hsa-miR-10a-5p | 0.68 | 0.0014 | Pregnant | ↑ |
| hsa-miR-10b-5p | 0.70 | 0.0205 | Pregnant | ↑ |
| hsa-miR-1180-3p | 1.57 | 0.0022 | Pregnant | ↑ |
| hsa-miR-125a-5p | 1.43 | 0.0022 | Pregnant | ↑ |
| hsa-miR-1307-3p | 1.76 | 0.0002 | Pregnant | ↑ |
| hsa-miR-193a-5p | 1.73 | 0.0119 | Pregnant | ↑ |
| hsa-miR-28-3p | 1.40 | 0.0006 | Pregnant | ↑ |
| hsa-miR-30a-5p | 0.58 | 0.0228 | Pregnant | ↑ |
| hsa-miR-30c-2-3p | 1.20 | 0.0303 | Pregnant | ↑ |
| hsa-miR-320a | 2.09 | 4.42E-06 | Pregnant | ↑ |
| hsa-miR-361-3p | 2.19 | 9.43E-07 | Pregnant | ↑ |
| hsa-miR-501-3p | 1.11 | 0.0258 | Pregnant | ↑ |
| hsa-miR-744-5p | 1.57 | 0.0048 | Pregnant | ↑ |
| hsa-miR-99b-3p | 1.41 | 0.0225 | Pregnant | ↑ |
| hsa-miR-100-5p | 0.85 | 0.0073 | Non-pregnant | ↑ |
| | 1.45 | 7.23E-07 | Pregnant | ↑ |
| hsa-miR-125b-5p | 0.84 | 0.0205 | Non-pregnant | ↑ |
| | 1.87 | 7.93E-06 | Pregnant | ↑ |
| hsa-miR-151a-3p | 0.94 | 0.0015 | Non-pregnant | ↑ |
| | 0.98 | 0.0003 | Pregnant | ↑ |
| hsa-miR-204-3p | 1.26 | 0.0062 | Non-pregnant | ↑ |
| | 2.65 | 2.79E-08 | Pregnant | ↑ |
| hsa-miR-204-5p | 0.96 | 0.0111 | Non-pregnant | ↑ |
| | 1.01 | 0.0135 | Pregnant | ↑ |
| hsa-miR-30d-5p | 0.82 | 1.33E-05 | Non-pregnant | ↑ |
| | 1.44 | 9.67E-15 | Pregnant | ↑ |
| hsa-miR-423-3p | 1.04 | 0.0009 | Non-pregnant | ↑ |
| | 1.44 | 3.40E-06 | Pregnant | ↑ |
| hsa-miR-423-5p | 1.29 | 0.0004 | Non-pregnant | ↑ |
| | 2.20 | 1.56E-05 | Pregnant | ↑ |
| hsa-miR-615-3p | 1.54 | 0.0475 | Non-pregnant | ↑ |
| | 2.81 | 0.0003 | Pregnant | ↑ |
| hsa-miR-99a-5p | 1.13 | 4.14E-06 | Non-pregnant | ↑ |

| | | | | |
|---|---|---|---|---|
| | 1.69 | 4.68E-10 | Pregnant | ↑ |
| hsa-miR-99b-5p | 1.32 | 0.0002 | Non-pregnant | ↑ |
| | 2.45 | 4.10E-21 | Pregnant | ↑ |
| **Enriched following total exRNA isolation (Decreased in vesicular exRNA isolation)** | | | | |
| hsa-miR-103a-3p | -1.47 | 0.0041 | Non-pregnant | ↓ |
| hsa-miR-22-3p | -0.91 | 0.0273 | Non-pregnant | ↓ |
| hsa-miR-23b-3p | -1.90 | 0.0148 | Non-pregnant | ↓ |
| hsa-miR-24-3p | -0.79 | 0.0037 | Non-pregnant | ↓ |
| hsa-miR-30b-5p | -1.50 | 0.0206 | Non-pregnant | ↓ |
| hsa-miR-30e-5p | -0.65 | 0.0344 | Non-pregnant | ↓ |
| hsa-miR-320b | -1.25 | 0.0206 | Non-pregnant | ↓ |
| hsa-miR-9-5p | -1.04 | 0.0080 | Non-pregnant | ↓ |
| hsa-let-7a-5p | -1.56 | 3.54E-09 | Pregnant | ↓ |
| hsa-let-7b-5p | -0.91 | 7.93E-06 | Pregnant | ↓ |
| hsa-let-7c-5p | -1.08 | 0.0007 | Pregnant | ↓ |
| hsa-miR-101-3p | -1.19 | 0.0061 | Pregnant | ↓ |
| hsa-miR-148a-3p | -0.90 | 0.0004 | Pregnant | ↓ |
| hsa-miR-196a-5p | -2.16 | 1.18E-05 | Pregnant | ↓ |
| hsa-miR-20a-5p | -2.83 | 0.0002 | Pregnant | ↓ |
| hsa-miR-224-5p | -1.52 | 0.0449 | Pregnant | ↓ |
| hsa-miR-30a-3p | -0.65 | 0.0240 | Pregnant | ↓ |
| hsa-miR-3182 | -2.89 | 2.05E-08 | Pregnant | ↓ |
| hsa-miR-629-5p | -2.03 | 2.20E-06 | Pregnant | ↓ |
| hsa-let-7f-5p | -1.22 | 0.0003 | Non-pregnant | ↓ |
| | -1.91 | 3.49E-11 | Pregnant | ↓ |
| hsa-let-7g-5p | -0.86 | 0.0115 | Non-pregnant | ↓ |
| | -1.40 | 0.0015 | Pregnant | ↓ |
| hsa-miR-200a-3p | -0.54 | 0.0126 | Non-pregnant | ↓ |
| | -0.63 | 0.0303 | Pregnant | ↓ |
| hsa-miR-203a-3p | -1.45 | 1.33E-05 | Non-pregnant | ↓ |
| | -1.96 | 2.13E-07 | Pregnant | ↓ |
| hsa-miR-21-5p | -0.58 | 0.0127 | Non-pregnant | ↓ |
| | -0.61 | 0.0090 | Pregnant | ↓ |
| hsa-miR-23a-3p | -1.58 | 0.0026 | Non-pregnant | ↓ |
| | -2.62 | 0.0090 | Pregnant | ↓ |
| hsa-miR-26a-5p | -0.84 | 0.0003 | Non-pregnant | ↓ |
| | -0.60 | 0.0341 | Pregnant | ↓ |
| hsa-miR-26b-5p | -1.39 | 9.88E-05 | Non-pregnant | ↓ |
| | -1.54 | 0.0008 | Pregnant | ↓ |
| hsa-miR-27a-3p | -2.11 | 1.90E-09 | Non-pregnant | ↓ |
| | -2.74 | 3.49E-11 | Pregnant | ↓ |
| hsa-miR-27b-3p | -1.45 | 2.43E-10 | Non-pregnant | ↓ |
| | -2.03 | 4.38E-12 | Pregnant | ↓ |

| | | | | |
|---|---|---|---|---|
| hsa-miR-429 | -1.26 | 0.00157 | Non-pregnant | ↓ |
| | -1.00 | 0.02289 | Pregnant | ↓ |
| hsa-miR-98-5p | -1.07 | 0.02002 | Non-pregnant | ↓ |
| | -1.57 | 0.00012 | Pregnant | ↓ |

**Supplementary Table 5:** MicroRNAs that were significantly differentially expressed in pregnant compared to non-pregnant urine in miRCURY-extracted samples (FDR<0.05).

| miRNA | Log fold-change | FDR | Expression in pregnant urine samples |
|---|---|---|---|
| ***Non-pregnant vs. 2nd trimester pregnant pools*** | | | |
| *hsa-miR-122-5p* | -3.22 | 0.0412 | Decreased in 2nd trimester pregnant pools |
| hsa-miR-30b-5p | -2.85 | 9.75E-06 | Decreased in 2nd trimester pregnant pools |
| hsa-miR-146a-5p | -2.73 | 2.70E-06 | Decreased in 2nd trimester pregnant pools |
| hsa-miR-660-5p | -2.67 | 0.0040 | Decreased in 2nd trimester pregnant pools |
| hsa-miR-197-3p | -2.12 | 0.0001 | Decreased in 2nd trimester pregnant pools |
| hsa-let-7e-5p | -1.72 | 0.0002 | Decreased in 2nd trimester pregnant pools |
| hsa-let-7d-5p | -1.69 | 0.0003 | Decreased in 2nd trimester pregnant pools |
| hsa-miR-141-3p | -1.59 | 0.0152 | Decreased in 2nd trimester pregnant pools |
| hsa-let-7a-5p | -1.56 | 3.90E-05 | Decreased in 2nd trimester pregnant pools |
| hsa-let-7f-5p | -1.15 | 0.0040 | Decreased in 2nd trimester pregnant pools |
| hsa-let-7c-5p | -1.10 | 8.15E-05 | Decreased in 2nd trimester pregnant pools |
| hsa-let-7b-5p | -1.05 | 9.12E-06 | Decreased in 2nd trimester pregnant pools |
| hsa-miR-27b-3p | -1.05 | 9.12E-06 | Decreased in 2nd trimester pregnant pools |
| hsa-miR-29a-3p | -1.02 | 0.0197 | Decreased in 2nd trimester pregnant pools |
| hsa-let-7g-5p | -0.99 | 7.70E-05 | Decreased in 2nd trimester pregnant pools |
| hsa-miR-30c-5p | -0.98 | 0.0003 | Decreased in 2nd trimester pregnant pools |
| hsa-miR-92a-3p | -0.94 | 0.0164 | Decreased in 2nd trimester pregnant pools |
| hsa-miR-192-5p | -0.80 | 0.0002 | Decreased in 2nd trimester pregnant pools |
| hsa-miR-30a-3p | -0.70 | 0.0019 | Decreased in 2nd trimester pregnant pools |
| hsa-miR-204-5p | -0.58 | 0.0197 | Decreased in 2nd trimester pregnant pools |
| hsa-miR-200a-3p | -0.57 | 0.0048 | Decreased in 2nd trimester pregnant pools |
| hsa-miR-10b-5p | -0.39 | 0.0326 | Decreased in 2nd trimester pregnant pools |
| hsa-miR-10a-5p | -0.37 | 0.0340 | Decreased in 2nd trimester pregnant pools |
| hsa-miR-1307-3p | 0.82 | 0.0316 | Increased in 2nd trimester pregnant pools |
| hsa-miR-203a-3p | 0.83 | 0.0041 | Increased in 2nd trimester pregnant pools |
| hsa-miR-361-3p | 0.84 | 0.0399 | Increased in 2nd trimester pregnant pools |
| hsa-miR-320b | 1.03 | 0.0340 | Increased in 2nd trimester pregnant pools |
| hsa-miR-320a | 1.18 | 0.0364 | Increased in 2nd trimester pregnant pools |
| hsa-miR-874-3p | 1.21 | 0.0326 | Increased in 2nd trimester pregnant pools |
| hsa-miR-193a-5p | 1.35 | 0.0341 | Increased in 2nd trimester pregnant pools |
| hsa-miR-125a-3p | 1.45 | 0.0273 | Increased in 2nd trimester pregnant pools |
| hsa-miR-664a-5p | 1.48 | 0.0268 | Increased in 2nd trimester pregnant pools |

| | | | |
|---|---|---|---|
| hsa-miR-320d | 1.54 | 0.0197 | Increased in 2nd trimester pregnant pools |
| hsa-miR-2110 | 1.63 | 0.0091 | Increased in 2nd trimester pregnant pools |
| hsa-miR-375 | 2.07 | 2.18E-06 | Increased in 2nd trimester pregnant pools |
| hsa-miR-3182 | 2.44 | 0.0011 | Increased in 2nd trimester pregnant pools |
| hsa-miR-3168 | 2.46 | 0.0197 | Increased in 2nd trimester pregnant pools |
| ***Non-pregnant individuals vs. 3rd trimester pregnant individuals*** | | | |
| hsa-miR-486-5p | -1.95 | 0.0002 | Decreased in pregnant individuals |
| *hsa-miR-122-5p* | -1.72 | 0.0012 | Decreased in pregnant individuals |
| hsa-miR-451a | -1.50 | 0.0012 | Decreased in pregnant individuals |
| hsa-miR-99a-5p | 0.77 | 0.0247 | Increased in pregnant individuals |
| hsa-miR-99b-5p | 1.02 | 0.0107 | Increased in pregnant individuals |
| hsa-miR-514a-3p | 1.38 | 0.0313 | Increased in pregnant individuals |
| ***2nd trimester pregnant pools vs. 3rd trimester pregnant individuals*** | | | |
| hsa-miR-874-3p | -2.22 | 0.0278 | Decreased in 3rd trimester pregnant individuals |
| hsa-miR-429 | -2.01 | 0.0031 | Decreased in 3rd trimester pregnant individuals |
| hsa-miR-30e-5p | -1.38 | 0.0007 | Decreased in 3rd trimester pregnant individuals |
| hsa-miR-26a-5p | -1.37 | 4.28346E-05 | Decreased in 3rd trimester pregnant individuals |
| hsa-miR-10b-5p | -1.19 | 5.8355E-06 | Decreased in 3rd trimester pregnant individuals |
| hsa-miR-125a-5p | -1.18 | 0.0046 | Decreased in 3rd trimester pregnant individuals |
| hsa-miR-10a-5p | -1.10 | 4.81559E-05 | Decreased in 3rd trimester pregnant individuals |
| hsa-miR-532-5p | -1.01 | 0.0029 | Decreased in 3rd trimester pregnant individuals |
| hsa-miR-200c-3p | -0.96 | 0.0147 | Decreased in 3rd trimester pregnant individuals |
| hsa-miR-30c-5p | -0.90 | 0.0273 | Decreased in 3rd trimester pregnant individuals |
| hsa-miR-21-5p | 0.88 | 0.0278 | Increased in 3rd trimester pregnant individuals |
| hsa-miR-192-5p | 0.99 | 0.0457 | Increased in 3rd trimester pregnant individuals |
| hsa-miR-514a-3p | 2.10 | 0.0019 | Increased in 3rd trimester pregnant individuals |
| hsa-miR-451a | 2.58 | 0.0200 | Increased in 3rd trimester pregnant individuals |
| hsa-miR-127-3p | 2.89 | 0.0148 | Increased in 3rd trimester pregnant individuals |
| hsa-miR-122-5p | 5.57 | 1.83E-14 | Increased in 3rd trimester pregnant individuals |
| hsa-miR-516b-5p | 11.00 | 9.1583E-08 | Increased in 3rd trimester pregnant individuals |

**Supplementary Table 6:** Protein-coding mRNAs that were significantly enriched following vesicular or total exRNA isolation that were unique to either nonpregnant or pregnant pools, as well as overlapping in both pools (FDR<0.05).

| ENSEMBL Gene ID | Gene description | HGNC symbol | Log fold-change (Total-Vesicular) | FDR | Pool | Vesicular expression |
|---|---|---|---|---|---|---|
| ***Enriched following vesicular exRNA isolation (miRCURY)*** | | | | | | |
| ENSG00000178761 | family with sequence similarity 219 member B [Source:HGNC Symbol;Acc:HGNC:24695] | *FAM219B* | -3.81 | 0.0367 | Non-pregnant | ↑ |
| ENSG00000063241 | isochorismatase domain containing 2 [Source:HGNC Symbol;Acc:HGNC:26278] | *ISOC2* | -3.21 | 0.0329 | Non-pregnant | ↑ |
| ENSG00000144035 | N-acetyltransferase 8 (putative) [Source:HGNC Symbol;Acc:HGNC:18069] | *NAT8* | -2.77 | 0.0343 | Non-pregnant | ↑ |
| ENSG00000165516 | kelch domain containing 2 [Source:HGNC Symbol;Acc:HGNC:20231] | *KLHDC2* | -2.67 | 0.0343 | Non-pregnant | ↑ |
| ENSG00000103316 | crystallin mu [Source:HGNC Symbol;Acc:HGNC:2418] | *CRYM* | -2.61 | 0.0353 | Non-pregnant | ↑ |
| ENSG00000214049 | urothelial cancer associated 1 [Source:HGNC Symbol;Acc:HGNC:37126] | *UCA1* | -2.46 | 0.0343 | Non-pregnant | ↑ |
| ENSG00000109919 | mitochondrial carrier 2 [Source:HGNC Symbol;Acc:HGNC:17587] | *MTCH2* | -1.95 | 0.0488 | Non-pregnant | ↑ |
| ENSG00000105552 | branched chain amino acid transaminase 2 [Source:HGNC Symbol;Acc:HGNC:977] | *BCAT2* | -1.79 | 0.0227 | Non-pregnant | ↑ |
| ENSG00000172663 | transmembrane protein 134 [Source:HGNC Symbol;Acc:HGNC:26142] | *TMEM134* | -1.68 | 0.0352 | Non-pregnant | ↑ |
| ENSG00000165678 | growth hormone inducible transmembrane protein [Source:HGNC Symbol;Acc:HGNC:17281] | *GHITM* | -1.25 | 0.0496 | Non-pregnant | ↑ |
| ENSG00000211445 | glutathione peroxidase 3 [Source:HGNC Symbol;Acc:HGNC:4555] | *GPX3* | -0.85 | 0.0218 | Non-pregnant | ↑ |
| ENSG00000188536 | hemoglobin subunit alpha 2 [Source:HGNC Symbol;Acc:HGNC:4824] | *HBA2* | -4.94 | 0.0092 | Pregnant | ↑ |
| ENSG00000162129 | caseinolytic mitochondrial matrix peptidase chaperone subunit B [Source:HGNC Symbol;Acc:HGNC:30664] | *CLPB* | -4.70 | 0.0496 | Pregnant | ↑ |
| ENSG00000163736 | pro-platelet basic protein [Source:HGNC Symbol;Acc:HGNC:9240] | *PPBP* | -4.59 | 0.0399 | Pregnant | ↑ |
| ENSG00000244734 | hemoglobin subunit beta [Source:HGNC Symbol;Acc:HGNC:4827] | *HBB* | -4.13 | 0.0032 | Pregnant | ↑ |

| | | | | | | |
|---|---|---|---|---|---|---|
| ENSG00000174132 | family with sequence similarity 174 member A [Source:HGNC Symbol;Acc:HGNC:24943] | *FAM174A* | -3.89 | 0.0500 | Pregnant | ↑ |
| ENSG00000127884 | enoyl-CoA hydratase, short chain 1 [Source:HGNC Symbol;Acc:HGNC:3151] | *ECHS1* | -1.26 | 0.0462 | Pregnant | ↑ |
| ENSG00000087086 | ferritin light chain [Source:HGNC Symbol;Acc:HGNC:3999] | *FTL* | -0.70 | 0.0005 | Pregnant | ↑ |
| ENSG00000158710 | transgelin 2 [Source:HGNC Symbol;Acc:HGNC:11554] | *TAGLN2* | -1.55 | 0.0312 | Non-pregnant | ↑ |
| | | | -2.22 | 0.0376 | Pregnant | ↑ |
| ENSG00000075624 | actin beta [Source:HGNC Symbol;Acc:HGNC:132] | *ACTB* | -1.31 | 0.0500 | Pregnant | ↑ |
| | | | -1.01 | 0.0343 | Non-pregnant | ↑ |
| ***Enriched following total exRNA isolation (Norgen)*** | | | | | | |
| ENSG00000143546 | S100 calcium binding protein A8 [Source:HGNC Symbol;Acc:HGNC:10498] | *S100A8* | 0.60 | 0.0352 | Non-pregnant | ↓ |
| ENSG00000071082 | ribosomal protein L31 [Source:HGNC Symbol;Acc:HGNC:10334] | *RPL31* | 0.74 | 0.0279 | Non-pregnant | ↓ |
| ENSG00000107485 | GATA binding protein 3 [Source:HGNC Symbol;Acc:HGNC:4172] | *GATA3* | 0.75 | 0.0318 | Non-pregnant | ↓ |
| ENSG00000133112 | tumor protein, translationally-controlled 1 [Source:HGNC Symbol;Acc:HGNC:12022] | *TPT1* | 0.77 | 0.0267 | Non-pregnant | ↓ |
| ENSG00000109475 | ribosomal protein L34 [Source:HGNC Symbol;Acc:HGNC:10340] | *RPL34* | 0.80 | 0.0343 | Non-pregnant | ↓ |
| ENSG00000131174 | cytochrome c oxidase subunit 7B [Source:HGNC Symbol;Acc:HGNC:2291] | *COX7B* | 0.84 | 0.0081 | Non-pregnant | ↓ |
| ENSG00000166441 | ribosomal protein L27a [Source:HGNC Symbol;Acc:HGNC:10329] | *RPL27A* | 0.84 | 0.0352 | Non-pregnant | ↓ |
| ENSG00000198034 | ribosomal protein S4 X-linked [Source:HGNC Symbol;Acc:HGNC:10424] | *RPS4X* | 0.85 | 0.0372 | Non-pregnant | ↓ |
| ENSG00000161016 | ribosomal protein L8 [Source:HGNC Symbol;Acc:HGNC:10368] | *RPL8* | 0.86 | 0.0341 | Non-pregnant | ↓ |
| ENSG00000149273 | ribosomal protein S3 [Source:HGNC Symbol;Acc:HGNC:10420] | *RPS3* | 0.90 | 0.0343 | Non-pregnant | ↓ |
| ENSG00000112306 | ribosomal protein S12 [Source:HGNC Symbol;Acc:HGNC:10385] | *RPS12* | 0.92 | 0.0318 | Non-pregnant | ↓ |
| ENSG00000122026 | ribosomal protein L21 [Source:HGNC Symbol;Acc:HGNC:10313] | *RPL21* | 0.95 | 0.0399 | Non-pregnant | ↓ |

| | | | | | | |
|---|---|---|---|---|---|---|
| ENSG00000174444 | ribosomal protein L4 [Source:HGNC Symbol;Acc:HGNC:10353] | *RPL4* | 0.95 | 0.0343 | Non-pregnant | ↓ |
| ENSG00000162522 | KIAA1522 [Source:HGNC Symbol;Acc:HGNC:29301] | *KIAA1522* | 0.97 | 0.0318 | Non-pregnant | ↓ |
| ENSG00000138326 | ribosomal protein S24 [Source:HGNC Symbol;Acc:HGNC:10411] | *RPS24* | 0.98 | 0.0486 | Non-pregnant | ↓ |
| ENSG00000198755 | ribosomal protein L10a [Source:HGNC Symbol;Acc:HGNC:10299] | *RPL10A* | 1.00 | 0.0458 | Non-pregnant | ↓ |
| ENSG00000177954 | ribosomal protein S27 [Source:HGNC Symbol;Acc:HGNC:10416] | *RPS27* | 1.01 | 0.0353 | Non-pregnant | ↓ |
| ENSG00000213741 | ribosomal protein S29 [Source:HGNC Symbol;Acc:HGNC:10419] | *RPS29* | 1.02 | 0.0181 | Non-pregnant | ↓ |
| ENSG00000182774 | ribosomal protein S17 [Source:HGNC Symbol;Acc:HGNC:10397] | *RPS17* | 1.06 | 0.0472 | Non-pregnant | ↓ |
| ENSG00000118898 | periplakin [Source:HGNC Symbol;Acc:HGNC:9273] | *PPL* | 1.06 | 0.0262 | Non-pregnant | ↓ |
| ENSG00000105193 | ribosomal protein S16 [Source:HGNC Symbol;Acc:HGNC:10396] | *RPS16* | 1.06 | 0.0369 | Non-pregnant | ↓ |
| ENSG00000156467 | ubiquinol-cytochrome c reductase binding protein [Source:HGNC Symbol;Acc:HGNC:12582] | *UQCRB* | 1.07 | 0.0267 | Non-pregnant | ↓ |
| ENSG00000126878 | allograft inflammatory factor 1 like [Source:HGNC Symbol;Acc:HGNC:28904] | *AIF1L* | 1.10 | 0.0455 | Non-pregnant | ↓ |
| ENSG00000148303 | ribosomal protein L7a [Source:HGNC Symbol;Acc:HGNC:10364] | *RPL7A* | 1.14 | 0.0343 | Non-pregnant | ↓ |
| ENSG00000080503 | SWI/SNF related, matrix associated, actin dependent regulator of chromatin, subfamily a, member 2 [Source:HGNC Symbol;Acc:HGNC:11098] | *SMARCA2* | 1.15 | 0.0399 | Non-pregnant | ↓ |
| ENSG00000163682 | ribosomal protein L9 [Source:HGNC Symbol;Acc:HGNC:10369] | *RPL9* | 1.18 | 0.0262 | Non-pregnant | ↓ |
| ENSG00000115306 | spectrin beta, non-erythrocytic 1 [Source:HGNC Symbol;Acc:HGNC:11275] | *SPTBN1* | 1.19 | 0.0352 | Non-pregnant | ↓ |
| ENSG00000161970 | ribosomal protein L26 [Source:HGNC Symbol;Acc:HGNC:10327] | *RPL26* | 1.19 | 0.0341 | Non-pregnant | ↓ |
| ENSG00000060237 | WNK lysine deficient protein kinase 1 [Source:HGNC Symbol;Acc:HGNC:14540] | *WNK1* | 1.24 | 0.0232 | Non-pregnant | ↓ |
| ENSG00000241251 | ribosomal protein L21 (RPL21) pseudogene | *ENSG00000241251* | 1.26 | 0.0487 | Non-pregnant | ↓ |
| ENSG00000100280 | adaptor related protein complex 1 subunit beta 1 [Source:HGNC Symbol;Acc:HGNC:554] | *AP1B1* | 1.27 | 0.0437 | Non-pregnant | ↓ |

| | | | | | | |
|---|---|---|---|---|---|---|
| ENSG00000113645 | WW and C2 domain containing 1 [Source:HGNC Symbol;Acc:HGNC:29435] | *WWC1* | 1.33 | 0.0159 | Non-pregnant | ↓ |
| ENSG00000104419 | N-myc downstream regulated 1 [Source:HGNC Symbol;Acc:HGNC:7679] | *NDRG1* | 1.42 | 0.0198 | Non-pregnant | ↓ |
| ENSG00000155506 | La ribonucleoprotein 1, translational regulator [Source:HGNC Symbol;Acc:HGNC:29531] | *LARP1* | 1.47 | 0.0399 | Non-pregnant | ↓ |
| ENSG00000158195 | WASP family member 2 [Source:HGNC Symbol;Acc:HGNC:12733] | *WASF2* | 1.48 | 0.0372 | Non-pregnant | ↓ |
| ENSG00000204387 | small nucleolar RNA host gene 32 [Source:HGNC Symbol;Acc:HGNC:19078] | *SNHG32* | 1.49 | 0.0449 | Non-pregnant | ↓ |
| ENSG00000167880 | envoplakin [Source:HGNC Symbol;Acc:HGNC:3503] | *EVPL* | 1.49 | 0.0006 | Non-pregnant | ↓ |
| ENSG00000163202 | late cornified envelope 3D [Source:HGNC Symbol;Acc:HGNC:16615] | *LCE3D* | 1.51 | 0.0006 | Non-pregnant | ↓ |
| ENSG00000275832 | Rho GTPase activating protein 23 [Source:HGNC Symbol;Acc:HGNC:29293] | *ARHGAP23* | 1.52 | 0.0357 | Non-pregnant | ↓ |
| ENSG00000170921 | tetratricopeptide repeat, ankyrin repeat and coiled-coil containing 2 [Source:HGNC Symbol;Acc:HGNC:30212] | *TANC2* | 1.52 | 0.0465 | Non-pregnant | ↓ |
| ENSG00000119185 | integrin subunit beta 1 binding protein 1 [Source:HGNC Symbol;Acc:HGNC:23927] | *ITGB1BP1* | 1.52 | 0.0263 | Non-pregnant | ↓ |
| ENSG00000132849 | PATJ crumbs cell polarity complex component [Source:HGNC Symbol;Acc:HGNC:28881] | *PATJ* | 1.60 | 0.0175 | Non-pregnant | ↓ |
| ENSG00000112379 | ARFGEF family member 3 [Source:HGNC Symbol;Acc:HGNC:21213] | *ARFGEF3* | 1.62 | 0.0455 | Non-pregnant | ↓ |
| ENSG00000173465 | zinc ribbon domain containing 2 [Source:HGNC Symbol;Acc:HGNC:11328] | *ZNRD2* | 1.65 | 0.0399 | Non-pregnant | ↓ |
| ENSG00000103342 | G1 to S phase transition 1 [Source:HGNC Symbol;Acc:HGNC:4621] | *GSPT1* | 1.70 | 0.0354 | Non-pregnant | ↓ |
| ENSG00000117713 | AT-rich interaction domain 1A [Source:HGNC Symbol;Acc:HGNC:11110] | *ARID1A* | 1.72 | 0.0339 | Non-pregnant | ↓ |
| ENSG00000163586 | fatty acid binding protein 1 [Source:HGNC Symbol;Acc:HGNC:3555] | *FABP1* | 1.74 | 0.0018 | Non-pregnant | ↓ |
| ENSG00000069020 | microtubule associated serine/threonine kinase family member 4 [Source:HGNC Symbol;Acc:HGNC:19037] | *MAST4* | 1.75 | 0.0353 | Non-pregnant | ↓ |
| ENSG00000169871 | tripartite motif containing 56 [Source:HGNC Symbol;Acc:HGNC:19028] | *TRIM56* | 1.77 | 0.0330 | Non-pregnant | ↓ |
| ENSG00000198938 | mitochondrially encoded cytochrome c oxidase III [Source:HGNC Symbol;Acc:HGNC:7422] | *MT-CO3* | 1.78 | 0.0353 | Non-pregnant | ↓ |

| | | | | | | |
|---|---|---|---|---|---|---|
| ENSG00000184831 | apolipoprotein O [Source:HGNC Symbol;Acc:HGNC:28727] | *APOO* | 1.80 | 0.0456 | Non-pregnant | ↓ |
| ENSG00000105698 | upstream transcription factor 2, c-fos interacting [Source:HGNC Symbol;Acc:HGNC:12594] | *USF2* | 1.81 | 0.0372 | Non-pregnant | ↓ |
| ENSG00000143153 | ATPase Na+/K+ transporting subunit beta 1 [Source:HGNC Symbol;Acc:HGNC:804] | *ATP1B1* | 1.84 | 0.0343 | Non-pregnant | ↓ |
| ENSG00000197879 | myosin IC [Source:HGNC Symbol;Acc:HGNC:7597] | *MYO1C* | 1.85 | 0.0085 | Non-pregnant | ↓ |
| ENSG00000038274 | methionine adenosyltransferase 2B [Source:HGNC Symbol;Acc:HGNC:6905] | *MAT2B* | 1.88 | 0.0417 | Non-pregnant | ↓ |
| ENSG00000131037 | EPS8 like 1 [Source:HGNC Symbol;Acc:HGNC:21295] | *EPS8L1* | 1.92 | 0.0195 | Non-pregnant | ↓ |
| ENSG00000069275 | nuclear casein kinase and cyclin dependent kinase substrate 1 [Source:HGNC Symbol;Acc:HGNC:29923] | *NUCKS1* | 1.94 | 0.0367 | Non-pregnant | ↓ |
| ENSG00000177885 | growth factor receptor bound protein 2 [Source:HGNC Symbol;Acc:HGNC:4566] | *GRB2* | 1.95 | 0.0318 | Non-pregnant | ↓ |
| ENSG00000143850 | pleckstrin homology domain containing A6 [Source:HGNC Symbol;Acc:HGNC:17053] | *PLEKHA6* | 1.98 | 0.0354 | Non-pregnant | ↓ |
| ENSG00000015171 | zinc finger MYND-type containing 11 [Source:HGNC Symbol;Acc:HGNC:16966] | *ZMYND11* | 2.00 | 0.0267 | Non-pregnant | ↓ |
| ENSG00000154380 | ENAH actin regulator [Source:HGNC Symbol;Acc:HGNC:18271] | *ENAH* | 2.06 | 0.0195 | Non-pregnant | ↓ |
| ENSG00000135631 | RAB11 family interacting protein 5 [Source:HGNC Symbol;Acc:HGNC:24845] | *RAB11FIP5* | 2.08 | 0.0343 | Non-pregnant | ↓ |
| ENSG00000130158 | dedicator of cytokinesis 6 [Source:HGNC Symbol;Acc:HGNC:19189] | *DOCK6* | 2.08 | 0.0343 | Non-pregnant | ↓ |
| ENSG00000170423 | keratin 78 [Source:HGNC Symbol;Acc:HGNC:28926] | *KRT78* | 2.10 | 0.0048 | Non-pregnant | ↓ |
| ENSG00000116871 | MAP7 domain containing 1 [Source:HGNC Symbol;Acc:HGNC:25514] | *MAP7D1* | 2.11 | 0.0249 | Non-pregnant | ↓ |
| ENSG00000198899 | mitochondrially encoded ATP synthase membrane subunit 6 [Source:HGNC Symbol;Acc:HGNC:7414] | *MT-ATP6* | 2.13 | 0.0496 | Non-pregnant | ↓ |
| ENSG00000153822 | potassium inwardly rectifying channel subfamily J member 16 [Source:HGNC Symbol;Acc:HGNC:6262] | *KCNJ16* | 2.14 | 0.0343 | Non-pregnant | ↓ |
| ENSG00000198886 | mitochondrially encoded NADH:ubiquinone oxidoreductase core subunit 4 [Source:HGNC Symbol;Acc:HGNC:7459] | *MT-ND4* | 2.16 | 0.0232 | Non-pregnant | ↓ |
| ENSG00000187867 | paralemmin 3 [Source:HGNC Symbol;Acc:HGNC:33274] | *PALM3* | 2.18 | 0.0262 | Non-pregnant | ↓ |
| ENSG00000198804 | mitochondrially encoded cytochrome c oxidase I [Source:HGNC Symbol;Acc:HGNC:7419] | *MT-CO1* | 2.19 | 0.0050 | Non-pregnant | ↓ |

| | | | | | | |
|---|---|---|---|---|---|---|
| ENSG00000168264 | interferon regulatory factor 2 binding protein 2 [Source:HGNC Symbol;Acc:HGNC:21729] | *IRF2BP2* | 2.21 | 0.0330 | Non-pregnant | ↓ |
| ENSG00000015133 | coiled-coil domain containing 88C [Source:HGNC Symbol;Acc:HGNC:19967] | *CCDC88C* | 2.27 | 0.0195 | Non-pregnant | ↓ |
| ENSG00000173273 | tankyrase [Source:HGNC Symbol;Acc:HGNC:11941] | *TNKS* | 2.31 | 0.0352 | Non-pregnant | ↓ |
| ENSG00000169925 | bromodomain containing 3 [Source:HGNC Symbol;Acc:HGNC:1104] | *BRD3* | 2.34 | 0.0353 | Non-pregnant | ↓ |
| ENSG00000089280 | FUS RNA binding protein [Source:HGNC Symbol;Acc:HGNC:4010] | *FUS* | 2.37 | 0.0352 | Non-pregnant | ↓ |
| ENSG00000103876 | fumarylacetoacetate hydrolase [Source:HGNC Symbol;Acc:HGNC:3579] | *FAH* | 2.40 | 0.0455 | Non-pregnant | ↓ |
| ENSG00000114302 | protein kinase cAMP-dependent type II regulatory subunit alpha [Source:HGNC Symbol;Acc:HGNC:9391] | *PRKAR2A* | 2.40 | 0.0341 | Non-pregnant | ↓ |
| ENSG00000164754 | RAD21 cohesin complex component [Source:HGNC Symbol;Acc:HGNC:9811] | *RAD21* | 2.41 | 0.0367 | Non-pregnant | ↓ |
| ENSG00000197928 | zinc finger protein 677 [Source:HGNC Symbol;Acc:HGNC:28730] | *ZNF677* | 2.45 | 0.0352 | Non-pregnant | ↓ |
| ENSG00000148396 | SEC16 homolog A, endoplasmic reticulum export factor [Source:HGNC Symbol;Acc:HGNC:29006] | *SEC16A* | 2.47 | 0.0343 | Non-pregnant | ↓ |
| ENSG00000107816 | leucine zipper tumor suppressor 2 [Source:HGNC Symbol;Acc:HGNC:29381] | *LZTS2* | 2.47 | 0.0488 | Non-pregnant | ↓ |
| ENSG00000108839 | arachidonate 12-lipoxygenase, 12S type [Source:HGNC Symbol;Acc:HGNC:429] | *ALOX12* | 2.50 | 0.0343 | Non-pregnant | ↓ |
| ENSG00000168066 | splicing factor 1 [Source:HGNC Symbol;Acc:HGNC:12950] | *SF1* | 2.55 | 0.0069 | Non-pregnant | ↓ |
| ENSG00000127481 | ubiquitin protein ligase E3 component n-recognin 4 [Source:HGNC Symbol;Acc:HGNC:30313] | *UBR4* | 2.56 | 0.0465 | Non-pregnant | ↓ |
| ENSG00000160305 | disco interacting protein 2 homolog A [Source:HGNC Symbol;Acc:HGNC:17217] | *DIP2A* | 2.56 | 0.0352 | Non-pregnant | ↓ |
| ENSG00000105397 | tyrosine kinase 2 [Source:HGNC Symbol;Acc:HGNC:12440] | *TYK2* | 2.57 | 0.0330 | Non-pregnant | ↓ |
| ENSG00000103550 | lysine rich nucleolar protein 1 [Source:HGNC Symbol;Acc:HGNC:34404] | *KNOP1* | 2.57 | 0.0341 | Non-pregnant | ↓ |
| ENSG00000076555 | acetyl-CoA carboxylase beta [Source:HGNC Symbol;Acc:HGNC:85] | *ACACB* | 2.63 | 0.0323 | Non-pregnant | ↓ |
| ENSG00000125741 | outer mitochondrial membrane lipid metabolism regulator OPA3 [Source:HGNC Symbol;Acc:HGNC:8142] | *OPA3* | 2.69 | 0.0353 | Non-pregnant | ↓ |

| | | | | | | |
|---|---|---|---|---|---|---|
| ENSG00000274523 | RCC1 like [Source:HGNC Symbol;Acc:HGNC:14948] | *RCC1L* | 2.69 | 0.0437 | Non-pregnant | ↓ |
| ENSG00000072786 | serine/threonine kinase 10 [Source:HGNC Symbol;Acc:HGNC:11388] | *STK10* | 2.77 | 0.0279 | Non-pregnant | ↓ |
| ENSG00000164631 | zinc finger protein 12 [Source:HGNC Symbol;Acc:HGNC:12902] | *ZNF12* | 2.77 | 0.0352 | Non-pregnant | ↓ |
| ENSG00000110917 | malectin [Source:HGNC Symbol;Acc:HGNC:28973] | *MLEC* | 2.79 | 0.0472 | Non-pregnant | ↓ |
| ENSG00000158321 | activator of transcription and developmental regulator AUTS2 [Source:HGNC Symbol;Acc:HGNC:14262] | *AUTS2* | 2.79 | 0.0488 | Non-pregnant | ↓ |
| ENSG00000224078 | small nucleolar RNA host gene 14 [Source:HGNC Symbol;Acc:HGNC:37462] | *SNHG14* | 2.81 | 0.0372 | Non-pregnant | ↓ |
| ENSG00000150457 | large tumor suppressor kinase 2 [Source:HGNC Symbol;Acc:HGNC:6515] | *LATS2* | 2.82 | 0.0352 | Non-pregnant | ↓ |
| ENSG00000064393 | homeodomain interacting protein kinase 2 [Source:HGNC Symbol;Acc:HGNC:14402] | *HIPK2* | 2.85 | 0.0120 | Non-pregnant | ↓ |
| ENSG00000248527 | MT-ATP6 pseudogene 1 [Source:HGNC Symbol;Acc:HGNC:44575] | *MTATP6P1* | 2.85 | 0.0449 | Non-pregnant | ↓ |
| ENSG00000103043 | VAC14 component of PIKFYVE complex [Source:HGNC Symbol;Acc:HGNC:25507] | *VAC14* | 2.87 | 0.0267 | Non-pregnant | ↓ |
| ENSG00000179023 | kelch domain containing 7A [Source:HGNC Symbol;Acc:HGNC:26791] | *KLHDC7A* | 2.90 | 0.0488 | Non-pregnant | ↓ |
| ENSG00000088888 | mitochondrial antiviral signaling protein [Source:HGNC Symbol;Acc:HGNC:29233] | *MAVS* | 2.90 | 0.0092 | Non-pregnant | ↓ |
| ENSG00000172831 | carboxylesterase 2 [Source:HGNC Symbol;Acc:HGNC:1864] | *CES2* | 2.92 | 0.0352 | Non-pregnant | ↓ |
| ENSG00000047617 | anoctamin 2 [Source:HGNC Symbol;Acc:HGNC:1183] | *ANO2* | 3.00 | 0.0279 | Non-pregnant | ↓ |
| ENSG00000280304 | TEC | *ENSG00000280304* | 3.07 | 0.0002 | Non-pregnant | ↓ |
| ENSG00000166016 | ankyrin repeat and BTB domain containing 2 [Source:HGNC Symbol;Acc:HGNC:23842] | *ABTB2* | 3.08 | 0.0262 | Non-pregnant | ↓ |
| ENSG00000108219 | tetraspanin 14 [Source:HGNC Symbol;Acc:HGNC:23303] | *TSPAN14* | 3.10 | 0.0006 | Non-pregnant | ↓ |
| ENSG00000141338 | ATP binding cassette subfamily A member 8 [Source:HGNC Symbol;Acc:HGNC:38] | *ABCA8* | 3.12 | 0.0120 | Non-pregnant | ↓ |
| ENSG00000205903 | zinc finger protein 316 [Source:HGNC Symbol;Acc:HGNC:13843] | *ZNF316* | 3.18 | 0.0343 | Non-pregnant | ↓ |
| ENSG00000054965 | family with sequence similarity 168 member A [Source:HGNC Symbol;Acc:HGNC:28999] | *FAM168A* | 3.21 | 0.0230 | Non-pregnant | ↓ |

| | | | | | | |
|---|---|---|---|---|---|---|
| ENSG00000181045 | solute carrier family 26 member 11 [Source:HGNC Symbol;Acc:HGNC:14471] | *SLC26A11* | 3.22 | 0.0369 | Non-pregnant | ↓ |
| ENSG00000188488 | serpin family A member 5 [Source:HGNC Symbol;Acc:HGNC:8723] | *SERPINA5* | 3.27 | 0.0159 | Non-pregnant | ↓ |
| ENSG00000171408 | phosphodiesterase 7B [Source:HGNC Symbol;Acc:HGNC:8792] | *PDE7B* | 3.28 | 0.0343 | Non-pregnant | ↓ |
| ENSG00000154914 | ubiquitin specific peptidase 43 [Source:HGNC Symbol;Acc:HGNC:20072] | *USP43* | 3.33 | 0.0417 | Non-pregnant | ↓ |
| ENSG00000145362 | ankyrin 2 [Source:HGNC Symbol;Acc:HGNC:493] | *ANK2* | 3.34 | 0.0330 | Non-pregnant | ↓ |
| ENSG00000188315 | chromosome 3 open reading frame 62 [Source:HGNC Symbol;Acc:HGNC:24771] | *C3orf62* | 3.34 | 0.0424 | Non-pregnant | ↓ |
| ENSG00000166508 | minichromosome maintenance complex component 7 [Source:HGNC Symbol;Acc:HGNC:6950] | *MCM7* | 3.34 | 0.0343 | Non-pregnant | ↓ |
| ENSG00000101966 | X-linked inhibitor of apoptosis [Source:HGNC Symbol;Acc:HGNC:592] | *XIAP* | 3.35 | 0.0069 | Non-pregnant | ↓ |
| ENSG00000125812 | GDNF inducible zinc finger protein 1 [Source:HGNC Symbol;Acc:HGNC:15808] | *GZF1* | 3.37 | 0.0352 | Non-pregnant | ↓ |
| ENSG00000214753 | heterogeneous nuclear ribonucleoprotein U like 2 [Source:HGNC Symbol;Acc:HGNC:25451] | *HNRNPUL2* | 3.38 | 0.0318 | Non-pregnant | ↓ |
| ENSG00000164080 | RAD54 like 2 [Source:HGNC Symbol;Acc:HGNC:29123] | *RAD54L2* | 3.38 | 0.0367 | Non-pregnant | ↓ |
| ENSG00000143756 | F-box protein 28 [Source:HGNC Symbol;Acc:HGNC:29046] | *FBXO28* | 3.41 | 0.0279 | Non-pregnant | ↓ |
| ENSG00000036672 | ubiquitin specific peptidase 2 [Source:HGNC Symbol;Acc:HGNC:12618] | *USP2* | 3.42 | 0.0038 | Non-pregnant | ↓ |
| ENSG00000148158 | sorting nexin family member 30 [Source:HGNC Symbol;Acc:HGNC:23685] | *SNX30* | 3.43 | 0.0372 | Non-pregnant | ↓ |
| ENSG00000155363 | Mov10 RISC complex RNA helicase [Source:HGNC Symbol;Acc:HGNC:7200] | *MOV10* | 3.44 | 0.0343 | Non-pregnant | ↓ |
| ENSG00000087250 | metallothionein 3 [Source:HGNC Symbol;Acc:HGNC:7408] | *MT3* | 3.44 | 0.0372 | Non-pregnant | ↓ |
| ENSG00000108262 | GIT ArfGAP 1 [Source:HGNC Symbol;Acc:HGNC:4272] | *GIT1* | 3.44 | 0.0367 | Non-pregnant | ↓ |
| ENSG00000120899 | protein tyrosine kinase 2 beta [Source:HGNC Symbol;Acc:HGNC:9612] | *PTK2B* | 3.45 | 0.0452 | Non-pregnant | ↓ |
| ENSG00000182308 | DDB1 and CUL4 associated factor 4 like 1 [Source:HGNC Symbol;Acc:HGNC:27723] | *DCAF4L1* | 3.45 | 0.0000 | Non-pregnant | ↓ |
| ENSG00000099864 | paralemmin [Source:HGNC Symbol;Acc:HGNC:8594] | *PALM* | 3.47 | 0.0137 | Non-pregnant | ↓ |

| | | | | | | |
|---|---|---|---|---|---|---|
| ENSG00000110237 | Rho guanine nucleotide exchange factor 17 [Source:HGNC Symbol;Acc:HGNC:21726] | *ARHGEF17* | 3.53 | 0.0279 | Non-pregnant | ↓ |
| ENSG00000189319 | family with sequence similarity 53 member B [Source:HGNC Symbol;Acc:HGNC:28968] | *FAM53B* | 3.54 | 0.0343 | Non-pregnant | ↓ |
| ENSG00000196365 | lon peptidase 1, mitochondrial [Source:HGNC Symbol;Acc:HGNC:9479] | *LONP1* | 3.57 | 0.0326 | Non-pregnant | ↓ |
| ENSG00000177133 | PRDM16 divergent transcript [Source:HGNC Symbol;Acc:HGNC:48664] | *PRDM16-DT* | 3.68 | 0.0055 | Non-pregnant | ↓ |
| ENSG00000158863 | FHF complex subunit HOOK interacting protein 2B [Source:HGNC Symbol;Acc:HGNC:16492] | *FHIP2B* | 3.68 | 0.0326 | Non-pregnant | ↓ |
| ENSG00000106266 | sorting nexin 8 [Source:HGNC Symbol;Acc:HGNC:14972] | *SNX8* | 3.73 | 0.0267 | Non-pregnant | ↓ |
| ENSG00000185818 | N-acetyltransferase 8 like [Source:HGNC Symbol;Acc:HGNC:26742] | *NAT8L* | 3.76 | 0.0434 | Non-pregnant | ↓ |
| ENSG00000124507 | protein kinase C and casein kinase substrate in neurons 1 [Source:HGNC Symbol;Acc:HGNC:8570] | *PACSIN1* | 3.78 | 0.0159 | Non-pregnant | ↓ |
| ENSG00000005810 | MYC binding protein 2 [Source:HGNC Symbol;Acc:HGNC:23386] | *MYCBP2* | 3.82 | 0.0000 | Non-pregnant | ↓ |
| ENSG00000133114 | GPALPP motifs containing 1 [Source:HGNC Symbol;Acc:HGNC:20298] | *GPALPP1* | 3.82 | 0.0100 | Non-pregnant | ↓ |
| ENSG00000087088 | BCL2 associated X, apoptosis regulator [Source:HGNC Symbol;Acc:HGNC:959] | *BAX* | 3.83 | 0.0001 | Non-pregnant | ↓ |
| ENSG00000261116 | novel transcript, overlapping to FAM83B | *ENSG00000261116* | 3.83 | 0.0341 | Non-pregnant | ↓ |
| ENSG00000129116 | palladin, cytoskeletal associated protein [Source:HGNC Symbol;Acc:HGNC:17068] | *PALLD* | 3.90 | 0.0092 | Non-pregnant | ↓ |
| ENSG00000071242 | ribosomal protein S6 kinase A2 [Source:HGNC Symbol;Acc:HGNC:10431] | *RPS6KA2* | 3.94 | 0.0067 | Non-pregnant | ↓ |
| ENSG00000137171 | kinesin light chain 4 [Source:HGNC Symbol;Acc:HGNC:21624] | *KLC4* | 3.94 | 0.0323 | Non-pregnant | ↓ |
| ENSG00000144645 | oxysterol binding protein like 10 [Source:HGNC Symbol;Acc:HGNC:16395] | *OSBPL10* | 3.98 | 0.0048 | Non-pregnant | ↓ |
| ENSG00000155657 | titin [Source:HGNC Symbol;Acc:HGNC:12403] | *TTN* | 4.00 | 0.0001 | Non-pregnant | ↓ |
| ENSG00000182544 | major facilitator superfamily domain containing 5 [Source:HGNC Symbol;Acc:HGNC:28156] | *MFSD5* | 4.03 | 0.0232 | Non-pregnant | ↓ |
| ENSG00000267106 | ZNF561 antisense RNA 1 (head to head) [Source:HGNC Symbol;Acc:HGNC:27613] | *ZNF561-AS1* | 4.04 | 0.0232 | Non-pregnant | ↓ |

| | | | | | | |
|---|---|---|---|---|---|---|
| ENSG00000153975 | zinc finger containing ubiquitin peptidase 1 [Source:HGNC Symbol;Acc:HGNC:21224] | *ZUP1* | 4.05 | 0.0343 | Non-pregnant | ↓ |
| ENSG00000140297 | glucosaminyl (N-acetyl) transferase 3, mucin type [Source:HGNC Symbol;Acc:HGNC:4205] | *GCNT3* | 4.07 | 0.0000 | Non-pregnant | ↓ |
| ENSG00000182749 | progestin and adipoQ receptor family member 7 [Source:HGNC Symbol;Acc:HGNC:23146] | *PAQR7* | 4.23 | 0.0092 | Non-pregnant | ↓ |
| ENSG00000257704 | InaF motif containing 1 [Source:HGNC Symbol;Acc:HGNC:27406] | *INAFM1* | 4.41 | 0.0120 | Non-pregnant | ↓ |
| ENSG00000122126 | OCRL inositol polyphosphate-5-phosphatase [Source:HGNC Symbol;Acc:HGNC:8108] | *OCRL* | 4.47 | 0.0040 | Non-pregnant | ↓ |
| ENSG00000280287 | novel transcript | *ENSG0000028 0287* | 4.69 | 0.0000 | Non-pregnant | ↓ |
| ENSG00000089050 | RB binding protein 9, serine hydrolase [Source:HGNC Symbol;Acc:HGNC:9892] | *RBBP9* | 5.09 | 0.0013 | Non-pregnant | ↓ |
| ENSG00000047662 | family with sequence similarity 184 member B [Source:HGNC Symbol;Acc:HGNC:29235] | *FAM184B* | 5.86 | 0.0000 | Non-pregnant | ↓ |
| ENSG00000249689 | chromosome 1 open reading frame 57 pseudogene | *ENSG0000024 9689* | 7.84 | 0.0000 | Non-pregnant | ↓ |
| ENSG00000203396 | WDR45-like (WDR45L) pseudogene | *ENSG0000020 3396* | 2.15 | 0.0013 | Pregnant | ↓ |
| ENSG00000170088 | transmembrane protein 192 [Source:HGNC Symbol;Acc:HGNC:26775] | *TMEM192* | 3.23 | 0.0500 | Pregnant | ↓ |
| ENSG00000156958 | galactokinase 2 [Source:HGNC Symbol;Acc:HGNC:4119] | *GALK2* | 3.74 | 0.0487 | Pregnant | ↓ |
| ENSG00000180543 | TSPY like 5 [Source:HGNC Symbol;Acc:HGNC:29367] | *TSPYL5* | 3.83 | 0.0290 | Pregnant | ↓ |
| ENSG00000145868 | F-box protein 38 [Source:HGNC Symbol;Acc:HGNC:28844] | *FBXO38* | 3.95 | 0.0404 | Pregnant | ↓ |
| ENSG00000143367 | tuftelin 1 [Source:HGNC Symbol;Acc:HGNC:12422] | *TUFT1* | 3.95 | 0.0404 | Pregnant | ↓ |
| ENSG00000144802 | NFKB inhibitor zeta [Source:HGNC Symbol;Acc:HGNC:29805] | *NFKBIZ* | 4.01 | 0.0399 | Pregnant | ↓ |
| ENSG00000110888 | caprin family member 2 [Source:HGNC Symbol;Acc:HGNC:21259] | *CAPRIN2* | 4.02 | 0.0450 | Pregnant | ↓ |
| ENSG00000214765 | septin 7 pseudogene 2 [Source:HGNC Symbol;Acc:HGNC:32339] | *SEPTIN7P2* | 4.12 | 0.0336 | Pregnant | ↓ |
| ENSG00000069702 | transforming growth factor beta receptor 3 [Source:HGNC Symbol;Acc:HGNC:11774] | *TGFBR3* | 4.14 | 0.0423 | Pregnant | ↓ |
| ENSG00000204839 | maestro heat like repeat family member 6 [Source:HGNC Symbol;Acc:HGNC:27814] | *MROH6* | 4.20 | 0.0404 | Pregnant | ↓ |

| | | | | | | |
|---|---|---|---|---|---|---|
| ENSG00000140948 | zinc finger CCHC-type containing 14 [Source:HGNC Symbol;Acc:HGNC:24134] | *ZCCHC14* | 4.23 | 0.0462 | Pregnant | ↓ |
| ENSG00000155252 | phosphatidylinositol 4-kinase type 2 alpha [Source:HGNC Symbol;Acc:HGNC:30031] | *PI4K2A* | 4.24 | 0.0404 | Pregnant | ↓ |
| ENSG00000214960 | CDP-L-ribitol pyrophosphorylase A [Source:HGNC Symbol;Acc:HGNC:37276] | *CRPPA* | 4.25 | 0.0500 | Pregnant | ↓ |
| ENSG00000248240 | novel transcript | *ENSG00000248240* | 4.31 | 0.0404 | Pregnant | ↓ |
| ENSG00000227835 | coactivator associated arginine methyltransferase 1 pseudogene 1 [Source:HGNC Symbol;Acc:HGNC:23392] | *CARM1P1* | 4.33 | 0.0404 | Pregnant | ↓ |
| ENSG00000122912 | solute carrier family 25 member 16 [Source:HGNC Symbol;Acc:HGNC:10986] | *SLC25A16* | 4.33 | 0.0325 | Pregnant | ↓ |
| ENSG00000277265 | RNA, 7SL, cytoplasmic 556, pseudogene [Source:HGNC Symbol;Acc:HGNC:46572] | *RN7SL556P* | 4.33 | 0.0404 | Pregnant | ↓ |
| ENSG00000279881 | TEC | *ENSG00000279881* | 4.36 | 0.0496 | Pregnant | ↓ |
| ENSG00000135587 | sphingomyelin phosphodiesterase 2 [Source:HGNC Symbol;Acc:HGNC:11121] | *SMPD2* | 4.37 | 0.0496 | Pregnant | ↓ |
| ENSG00000146676 | purine rich element binding protein B [Source:HGNC Symbol;Acc:HGNC:9702] | *PURB* | 4.37 | 0.0476 | Pregnant | ↓ |
| ENSG00000051825 | M-phase phosphoprotein 9 [Source:HGNC Symbol;Acc:HGNC:7215] | *MPHOSPH9* | 4.41 | 0.0476 | Pregnant | ↓ |
| ENSG00000154305 | MIA SH3 domain ER export factor 3 [Source:HGNC Symbol;Acc:HGNC:24008] | *MIA3* | 4.42 | 0.0399 | Pregnant | ↓ |
| ENSG00000232696 | novel transcript | *ENSG00000232696* | 4.43 | 0.0496 | Pregnant | ↓ |
| ENSG00000165192 | ankyrin repeat and SOCS box containing 11 [Source:HGNC Symbol;Acc:HGNC:17186] | *ASB11* | 4.44 | 0.0292 | Pregnant | ↓ |
| ENSG00000154839 | spindle and kinetochore associated complex subunit 1 [Source:HGNC Symbol;Acc:HGNC:28109] | *SKA1* | 4.45 | 0.0399 | Pregnant | ↓ |
| ENSG00000125885 | minichromosome maintenance 8 homologous recombination repair factor [Source:HGNC Symbol;Acc:HGNC:16147] | *MCM8* | 4.46 | 0.0404 | Pregnant | ↓ |
| ENSG00000241956 | novel transcript | *ENSG00000241956* | 4.46 | 0.0299 | Pregnant | ↓ |
| ENSG00000125449 | armadillo repeat containing 7 [Source:HGNC Symbol;Acc:HGNC:26168] | *ARMC7* | 4.50 | 0.0404 | Pregnant | ↓ |

| | | | | | | |
|---|---|---|---|---|---|---|
| ENSG00000129295 | dynein axonemal assembly factor 11 [Source:HGNC Symbol;Acc:HGNC:16725] | *DNAAF11* | 4.51 | 0.0428 | Pregnant | ↓ |
| ENSG00000143398 | phosphatidylinositol-4-phosphate 5-kinase type 1 alpha [Source:HGNC Symbol;Acc:HGNC:8994] | *PIP5K1A* | 4.53 | 0.0346 | Pregnant | ↓ |
| ENSG00000223125 | RNA, U2 small nuclear 32, pseudogene [Source:HGNC Symbol;Acc:HGNC:48525] | *RNU2-32P* | 4.55 | 0.0404 | Pregnant | ↓ |
| ENSG00000271811 | novel transcript | *ENSG00000271811* | 4.57 | 0.0476 | Pregnant | ↓ |
| ENSG00000270885 | RAS like family 10 member B [Source:HGNC Symbol;Acc:HGNC:30295] | *RASL10B* | 4.57 | 0.0446 | Pregnant | ↓ |
| ENSG00000267060 | prostaglandin E synthase 3 like [Source:HGNC Symbol;Acc:HGNC:43943] | *PTGES3L* | 4.58 | 0.0399 | Pregnant | ↓ |
| ENSG00000088543 | chromosome 3 open reading frame 18 [Source:HGNC Symbol;Acc:HGNC:24837] | *C3orf18* | 4.63 | 0.0290 | Pregnant | ↓ |
| ENSG00000279043 | TEC | *ENSG00000279043* | 4.67 | 0.0299 | Pregnant | ↓ |
| ENSG00000096093 | EF-hand domain containing 1 [Source:HGNC Symbol;Acc:HGNC:16406] | *EFHC1* | 4.67 | 0.0292 | Pregnant | ↓ |
| ENSG00000226356 | ribosomal protein S6 pseudogene 20 [Source:HGNC Symbol;Acc:HGNC:36940] | *RPS6P20* | 4.71 | 0.0325 | Pregnant | ↓ |
| ENSG00000243297 | ribosomal protein L31 pseudogene 61 [Source:HGNC Symbol;Acc:HGNC:35635] | *RPL31P61* | 4.75 | 0.0399 | Pregnant | ↓ |
| ENSG00000254605 | novel transcript | *ENSG00000254605* | 4.82 | 0.0273 | Pregnant | ↓ |
| ENSG00000279345 | novel transcript, antisense to BRD1 | *ENSG00000279345* | 4.82 | 0.0290 | Pregnant | ↓ |
| ENSG00000214369 | cytochrome b5 type A pseudogene 2 [Source:HGNC Symbol;Acc:HGNC:2573] | *CYB5AP2* | 4.86 | 0.0290 | Pregnant | ↓ |
| ENSG00000105122 | RAS protein activator like 3 [Source:HGNC Symbol;Acc:HGNC:26129] | *RASAL3* | 4.87 | 0.0336 | Pregnant | ↓ |
| ENSG00000264618 | RNA, 7SL, cytoplasmic 732, pseudogene [Source:HGNC Symbol;Acc:HGNC:46748] | *RN7SL732P* | 4.94 | 0.0280 | Pregnant | ↓ |
| ENSG00000134899 | ERCC excision repair 5, endonuclease [Source:HGNC Symbol;Acc:HGNC:3437] | *ERCC5* | 4.95 | 0.0325 | Pregnant | ↓ |
| ENSG00000144837 | phospholipase A1 member A [Source:HGNC Symbol;Acc:HGNC:17661] | *PLA1A* | 4.96 | 0.0292 | Pregnant | ↓ |
| ENSG00000279948 | TEC | *ENSG00000279948* | 4.97 | 0.0177 | Pregnant | ↓ |

| | | | | | | |
|---|---|---|---|---|---|---|
| ENSG00000257027 | novel transcript | *ENSG0000025 7027* | 5.02 | 0.0368 | Pregnant | ↓ |
| ENSG00000254726 | mex-3 RNA binding family member A [Source:HGNC Symbol;Acc:HGNC:33482] | *MEX3A* | 5.07 | 0.0282 | Pregnant | ↓ |
| ENSG00000244532 | RNA, 7SL, cytoplasmic 380, pseudogene [Source:HGNC Symbol;Acc:HGNC:46396] | *RN7SL380P* | 5.11 | 0.0249 | Pregnant | ↓ |
| ENSG00000265813 | RNA, 7SL, cytoplasmic 300, pseudogene [Source:HGNC Symbol;Acc:HGNC:46316] | *RN7SL300P* | 5.12 | 0.0159 | Pregnant | ↓ |
| **ENSG00000182752*** | **pappalysin 1 [Source:HGNC Symbol;Acc:HGNC:8602]** | **PAPPA** | **5.13** | **0.0153** | **Pregnant** | ↓ |
| **ENSG00000169550*** | **mucin 15, cell surface associated [Source:HGNC Symbol;Acc:HGNC:14956]** | **MUC15** | **5.14** | **0.0301** | **Pregnant** | ↓ |
| ENSG00000261795 | novel transcript | *ENSG0000026 1795* | 5.15 | 0.0325 | Pregnant | ↓ |
| ENSG00000107165 | tyrosinase related protein 1 [Source:HGNC Symbol;Acc:HGNC:12450] | *TYRP1* | 5.19 | 0.0292 | Pregnant | ↓ |
| ENSG00000163393 | solute carrier family 22 member 15 [Source:HGNC Symbol;Acc:HGNC:20301] | *SLC22A15* | 5.25 | 0.0290 | Pregnant | ↓ |
| ENSG00000158683 | polycystin 1 like 1, transient receptor potential channel interacting [Source:HGNC Symbol;Acc:HGNC:18053] | *PKD1L1* | 5.26 | 0.0346 | Pregnant | ↓ |
| **ENSG00000116183*** | **pappalysin 2 [Source:HGNC Symbol;Acc:HGNC:14615]** | **PAPPA2** | **5.62** | **0.0126** | **Pregnant** | ↓ |
| ENSG00000124459 | zinc finger protein 45 [Source:HGNC Symbol;Acc:HGNC:13111] | *ZNF45* | 6.40 | 0.0038 | Pregnant | ↓ |
| ENSG00000143226 | Fc fragment of IgG receptor IIa [Source:HGNC Symbol;Acc:HGNC:3616] | *FCGR2A* | 2.41 | 0.0048 | Non-pregnant | ↓ |
| | | | 2.37 | 0.0032 | Pregnant | ↓ |
| ENSG00000158079 | protein tyrosine phosphatase domain containing 1 [Source:HGNC Symbol;Acc:HGNC:30184] | *PTPDC1* | 5.55 | 0.0000 | Non-pregnant | ↓ |
| | | | 2.43 | 0.0159 | Pregnant | ↓ |
| ENSG00000160551 | TAO kinase 1 [Source:HGNC Symbol;Acc:HGNC:29259] | *TAOK1* | 2.06 | 0.0007 | Non-pregnant | ↓ |
| | | | 2.02 | 0.0071 | Pregnant | ↓ |
| ENSG00000182218 | HHIP like 1 [Source:HGNC Symbol;Acc:HGNC:19710] | *HHIPL1* | 5.97 | 0.0000 | Non-pregnant | ↓ |
| | | | 2.74 | 0.0014 | Pregnant | ↓ |

| | | | | | | |
|---|---|---|---|---|---|---|
| ENSG00000198695 | mitochondrially encoded NADH:ubiquinone oxidoreductase core subunit 6 [Source:HGNC Symbol;Acc:HGNC:7462] | *MT-ND6* | 2.19 | 0.0022 | Non-pregnant | ↓ |
| | | | 2.57 | 0.0159 | Pregnant | ↓ |
| ENSG00000274211 | suppressor of cytokine signaling 7 [Source:HGNC Symbol;Acc:HGNC:29846] | *SOCS7* | 2.34 | 0.0007 | Non-pregnant | ↓ |
| | | | 2.42 | 0.0005 | Pregnant | ↓ |

***Enriched in placental tissue compared to other tissue types**

**Supplementary table 7:** Protein-coding mRNAs that were significantly differentially expressed in pregnant compared to non-pregnant urine samples (FDR<0.05).

| **ENSEMBL Gene ID** | **Gene description** | **HGNC symbol** | **Log fold-change** | **FDR** | **Expression in pregnant urine samples** |
|---|---|---|---|---|---|
| ***Non-pregnant pools vs. 2nd trimester pregnant pools*** | | | | | |
| ENSG00000160991 | ORAI calcium release-activated calcium modulator 2 [Source:HGNC Symbol;Acc:HGNC:21667] | *ORAI2* | -3.30 | 0.0224 | Decreased in 2nd trimester pregnant pools |
| ENSG00000137831 | uveal autoantigen with coiled-coil domains and ankyrin repeats [Source:HGNC Symbol;Acc:HGNC:15947] | *UACA* | -2.45 | 0.0028 | Decreased in 2nd trimester pregnant pools |
| ENSG00000214049 | urothelial cancer associated 1 [Source:HGNC Symbol;Acc:HGNC:37126] | *UCA1* | -2.39 | 0.0073 | Decreased in 2nd trimester pregnant pools |
| ENSG00000165685 | transmembrane protein 52B [Source:HGNC Symbol;Acc:HGNC:26438] | *TMEM52B* | -1.60 | 0.0029 | Decreased in 2nd trimester pregnant pools |
| ENSG00000104267 | carbonic anhydrase 2 [Source:HGNC Symbol;Acc:HGNC:1373] | *CA2* | -1.42 | 0.0431 | Decreased in 2nd trimester pregnant pools |

| | | | | | |
|---|---|---|---|---|---|
| ENSG00000211445 | glutathione peroxidase 3 [Source:HGNC Symbol;Acc:HGNC:4555] | *GPX3* | -0.87 | 0.0002 | Decreased in 2nd trimester pregnant pools |
| ENSG00000163216 | small proline rich protein 2D [Source:HGNC Symbol;Acc:HGNC:11264] | *SPRR2D* | 0.92 | 0.0277 | Increased in 2nd trimester pregnant pools |
| ENSG00000115468 | EF-hand domain family member D1 [Source:HGNC Symbol;Acc:HGNC:29556] | *EFHD1* | 1.01 | 0.0377 | Increased in 2nd trimester pregnant pools |
| ENSG00000118898 | periplakin [Source:HGNC Symbol;Acc:HGNC:9273] | *PPL* | 1.05 | 0.0070 | Increased in 2nd trimester pregnant pools |
| ENSG00000172005 | mal, T cell differentiation protein [Source:HGNC Symbol;Acc:HGNC:6817] | *MAL* | 1.08 | 0.0006 | Increased in 2nd trimester pregnant pools |
| ENSG00000169474 | small proline rich protein 1A [Source:HGNC Symbol;Acc:HGNC:11259] | *SPRR1A* | 1.19 | 0.0023 | Increased in 2nd trimester pregnant pools |
| ENSG00000160213 | cystatin B [Source:HGNC Symbol;Acc:HGNC:2482] | *CSTB* | 1.21 | 0.0000 | Increased in 2nd trimester pregnant pools |
| ENSG00000116717 | growth arrest and DNA damage inducible alpha [Source:HGNC Symbol;Acc:HGNC:4095] | *GADD45A* | 1.26 | 0.0310 | Increased in 2nd trimester pregnant pools |
| ENSG00000169469 | small proline rich protein 1B [Source:HGNC Symbol;Acc:HGNC:11260] | *SPRR1B* | 1.33 | 0.0000 | Increased in 2nd trimester pregnant pools |
| ENSG00000241794 | small proline rich protein 2A [Source:HGNC Symbol;Acc:HGNC:11261] | *SPRR2A* | 1.36 | 0.0000 | Increased in 2nd trimester pregnant pools |
| ENSG00000134184 | glutathione S-transferase mu 1 [Source:HGNC Symbol;Acc:HGNC:4632] | *GSTM1* | 1.39 | 0.0225 | Increased in 2nd trimester pregnant pools |
| ENSG00000143546 | S100 calcium binding protein A8 [Source:HGNC Symbol;Acc:HGNC:10498] | *S100A8* | 1.46 | 5.75E-08 | Increased in 2nd trimester pregnant pools |
| ENSG00000166165 | creatine kinase B [Source:HGNC Symbol;Acc:HGNC:1991] | *CKB* | 1.47 | 0.0169 | Increased in 2nd trimester pregnant pools |

| | | | | | |
|---|---|---|---|---|---|
| ENSG00000148346 | lipocalin 2 [Source:HGNC Symbol;Acc:HGNC:6526] | *LCN2* | 1.48 | 0.0005 | Increased in 2nd trimester pregnant pools |
| ENSG00000121552 | cystatin A [Source:HGNC Symbol;Acc:HGNC:2481] | *CSTA* | 1.50 | 0.0034 | Increased in 2nd trimester pregnant pools |
| ENSG00000188100 | family with sequence similarity 25 member A [Source:HGNC Symbol;Acc:HGNC:23436] | *FAM25A* | 1.61 | 0.0015 | Increased in 2nd trimester pregnant pools |
| ENSG00000140519 | Rh family C glycoprotein [Source:HGNC Symbol;Acc:HGNC:18140] | *RHCG* | 1.61 | 0.0005 | Increased in 2nd trimester pregnant pools |
| ENSG00000105427 | cornifelin [Source:HGNC Symbol;Acc:HGNC:30183] | *CNFN* | 1.62 | 0.0000 | Increased in 2nd trimester pregnant pools |
| ENSG00000135046 | annexin A1 [Source:HGNC Symbol;Acc:HGNC:533] | *ANXA1* | 1.63 | 0.0000 | Increased in 2nd trimester pregnant pools |
| ENSG00000163209 | small proline rich protein 3 [Source:HGNC Symbol;Acc:HGNC:11268] | *SPRR3* | 1.64 | 4.05E-08 | Increased in 2nd trimester pregnant pools |
| ENSG00000143536 | cornulin [Source:HGNC Symbol;Acc:HGNC:1230] | *CRNN* | 1.65 | 0.0000 | Increased in 2nd trimester pregnant pools |
| ENSG00000197191 | cysteine rich tail 1 [Source:HGNC Symbol;Acc:HGNC:30529] | *CYSRT1* | 1.67 | 0.0001 | Increased in 2nd trimester pregnant pools |
| ENSG00000169509 | cysteine rich C-terminal 1 [Source:HGNC Symbol;Acc:HGNC:29875] | *CRCT1* | 1.69 | 0.0000 | Increased in 2nd trimester pregnant pools |
| ENSG00000163220 | S100 calcium binding protein A9 [Source:HGNC Symbol;Acc:HGNC:10499] | *S100A9* | 1.70 | 4.05E-08 | Increased in 2nd trimester pregnant pools |
| ENSG00000171401 | keratin 13 [Source:HGNC Symbol;Acc:HGNC:6415] | *KRT13* | 1.71 | 0.0000 | Increased in 2nd trimester pregnant pools |
| ENSG00000124107 | secretory leukocyte peptidase inhibitor [Source:HGNC Symbol;Acc:HGNC:11092] | *SLPI* | 1.73 | 0.0277 | Increased in 2nd trimester pregnant pools |

| | | | | | |
|---|---|---|---|---|---|
| ENSG00000185966 | late cornified envelope 3E [Source:HGNC Symbol;Acc:HGNC:29463] | *LCE3E* | 1.83 | 0.0254 | Increased in 2nd trimester pregnant pools |
| ENSG00000108839 | arachidonate 12-lipoxygenase, 12S type [Source:HGNC Symbol;Acc:HGNC:429] | *ALOX12* | 2.10 | 0.0436 | Increased in 2nd trimester pregnant pools |
| ENSG00000095383 | TBC1 domain family member 2 [Source:HGNC Symbol;Acc:HGNC:18026] | *TBC1D2* | 2.12 | 0.0254 | Increased in 2nd trimester pregnant pools |
| ENSG00000166920 | chromosome 15 open reading frame 48 [Source:HGNC Symbol;Acc:HGNC:29898] | *C15orf48* | 2.14 | 0.0003 | Increased in 2nd trimester pregnant pools |
| ENSG00000198682 | 3'-phosphoadenosine 5'-phosphosulfate synthase 2 [Source:HGNC Symbol;Acc:HGNC:8604] | *PAPSS2* | 2.21 | 0.0358 | Increased in 2nd trimester pregnant pools |
| ENSG00000118579 | mediator complex subunit 28 [Source:HGNC Symbol;Acc:HGNC:24628] | *MED28* | 2.22 | 0.0466 | Increased in 2nd trimester pregnant pools |
| ENSG00000145879 | serine peptidase inhibitor Kazal type 7 [Source:HGNC Symbol;Acc:HGNC:24643] | *SPINK7* | 2.24 | 0.0000 | Increased in 2nd trimester pregnant pools |
| ENSG00000165949 | interferon alpha inducible protein 27 [Source:HGNC Symbol;Acc:HGNC:5397] | *IFI27* | 2.29 | 0.0254 | Increased in 2nd trimester pregnant pools |
| ENSG00000170423 | keratin 78 [Source:HGNC Symbol;Acc:HGNC:28926] | *KRT78* | 2.32 | 0.0048 | Increased in 2nd trimester pregnant pools |
| ENSG00000151704 | potassium inwardly rectifying channel subfamily J member 1 [Source:HGNC Symbol;Acc:HGNC:6255] | *KCNJ1* | 2.35 | 0.0377 | Increased in 2nd trimester pregnant pools |
| ENSG00000109854 | HIV-1 Tat interactive protein 2 [Source:HGNC Symbol;Acc:HGNC:16637] | *HTATIP2* | 2.42 | 0.0104 | Increased in 2nd trimester pregnant pools |
| ENSG00000064652 | sorting nexin 24 [Source:HGNC Symbol;Acc:HGNC:21533] | *SNX24* | 2.56 | 0.0252 | Increased in 2nd trimester pregnant pools |
| ENSG00000088002 | sulfotransferase family 2B member 1 [Source:HGNC Symbol;Acc:HGNC:11459] | *SULT2B1* | 2.60 | 0.0113 | Increased in 2nd trimester pregnant pools |

| | | | | | |
|---|---|---|---|---|---|
| ENSG00000125834 | serine/threonine kinase 35 [Source:HGNC Symbol;Acc:HGNC:16254] | *STK35* | 2.61 | 0.0301 | Increased in 2nd trimester pregnant pools |
| ENSG00000019144 | pleckstrin homology like domain family B member 1 [Source:HGNC Symbol;Acc:HGNC:23697] | *PHLDB1* | 2.63 | 0.0029 | Increased in 2nd trimester pregnant pools |
| ENSG00000163202 | late cornified envelope 3D [Source:HGNC Symbol;Acc:HGNC:16615] | *LCE3D* | 3.03 | 1.61E-09 | Increased in 2nd trimester pregnant pools |
| ENSG00000171747 | galectin 4 [Source:HGNC Symbol;Acc:HGNC:6565] | *LGALS4* | 3.13 | 0.0491 | Increased in 2nd trimester pregnant pools |
| ENSG00000188001 | tumor protein p63 regulated 1 [Source:HGNC Symbol;Acc:HGNC:24759] | *TPRG1* | 3.26 | 0.0224 | Increased in 2nd trimester pregnant pools |
| ENSG00000267106 | ZNF561 antisense RNA 1 (head to head) [Source:HGNC Symbol;Acc:HGNC:27613] | *ZNF561-AS1* | 3.54 | 0.0087 | Increased in 2nd trimester pregnant pools |
| ***Non-pregnant individuals vs. 3rd trimester pregnant individuals*** | | | | | |
| ENSG00000169727 | G protein pathway suppressor 1 [Source:HGNC Symbol;Acc:HGNC:4549] | *GPS1* | 7.35 | 1.3042E-06 | Increased in 3rd trimester pregnant individuals |
| ENSG00000124762 | cyclin dependent kinase inhibitor 1A [Source:HGNC Symbol;Acc:HGNC:1784] | *CDKN1A* | 6.78 | 0.0001 | Increased in 3rd trimester pregnant individuals |
| ENSG00000143320 | cellular retinoic acid binding protein 2 [Source:HGNC Symbol;Acc:HGNC:2339] | *CRABP2* | 5.99 | 0.0040 | Increased in 3rd trimester pregnant individuals |
| ENSG00000166797 | cytosolic iron-sulfur assembly component 2A [Source:HGNC Symbol;Acc:HGNC:26235] | *CIAO2A* | 5.84 | 0.0016 | Increased in 3rd trimester pregnant individuals |
| ENSG00000103316 | crystallin mu [Source:HGNC Symbol;Acc:HGNC:2418] | *CRYM* | 5.84 | 0.0064 | Increased in 3rd trimester pregnant individuals |
| ENSG00000124596 | O-acyl-ADP-ribose deacylase 1 [Source:HGNC Symbol;Acc:HGNC:21257] | *OARD1* | 5.81 | 0.0020 | Increased in 3rd trimester pregnant individuals |

| | | | | | |
|---|---|---|---|---|---|
| ENSG00000075239 | acetyl-CoA acetyltransferase 1 [Source:HGNC Symbol;Acc:HGNC:93] | *ACAT1* | 5.78 | 0.0053 | Increased in 3rd trimester pregnant individuals |
| ENSG00000182054 | isocitrate dehydrogenase (NADP(+)) 2 [Source:HGNC Symbol;Acc:HGNC:5383] | *IDH2* | 5.73 | 0.0057 | Increased in 3rd trimester pregnant individuals |
| ENSG00000171988 | jumonji domain containing 1C [Source:HGNC Symbol;Acc:HGNC:12313] | *JMJD1C* | 5.69 | 0.0026 | Increased in 3rd trimester pregnant individuals |
| ENSG00000084093 | RE1 silencing transcription factor [Source:HGNC Symbol;Acc:HGNC:9966] | *REST* | 5.66 | 0.0058 | Increased in 3rd trimester pregnant individuals |
| ENSG00000085832 | epidermal growth factor receptor pathway substrate 15 [Source:HGNC Symbol;Acc:HGNC:3419] | *EPS15* | 5.59 | 0.0047 | Increased in 3rd trimester pregnant individuals |
| ENSG00000171475 | WAS/WASL interacting protein family member 2 [Source:HGNC Symbol;Acc:HGNC:30923] | *WIPF2* | 5.53 | 0.0026 | Increased in 3rd trimester pregnant individuals |
| ENSG00000124226 | ring finger protein 114 [Source:HGNC Symbol;Acc:HGNC:13094] | *RNF114* | 5.52 | 0.0031 | Increased in 3rd trimester pregnant individuals |
| ENSG00000119541 | vacuolar protein sorting 4 homolog B [Source:HGNC Symbol;Acc:HGNC:10895] | *VPS4B* | 5.48 | 0.0065 | Increased in 3rd trimester pregnant individuals |
| ENSG00000107521 | HPS1 biogenesis of lysosomal organelles complex 3 subunit 1 [Source:HGNC Symbol;Acc:HGNC:5163] | *HPS1* | 5.44 | 0.0058 | Increased in 3rd trimester pregnant individuals |
| ENSG00000161714 | phospholipase C delta 3 [Source:HGNC Symbol;Acc:HGNC:9061] | *PLCD3* | 5.44 | 0.0047 | Increased in 3rd trimester pregnant individuals |
| ENSG00000013583 | heme binding protein 1 [Source:HGNC Symbol;Acc:HGNC:17176] | *HEBP1* | 5.43 | 0.0091 | Increased in 3rd trimester pregnant individuals |
| ENSG00000196230 | tubulin beta class I [Source:HGNC Symbol;Acc:HGNC:20778] | *TUBB* | 5.40 | 0.0020 | Increased in 3rd trimester pregnant individuals |
| ENSG00000188313 | phospholipid scramblase 1 [Source:HGNC Symbol;Acc:HGNC:9092] | *PLSCR1* | 5.39 | 0.0160 | Increased in 3rd trimester pregnant individuals |

| | | | | | |
|---|---|---|---|---|---|
| ENSG00000004478 | FKBP prolyl isomerase 4 [Source:HGNC Symbol;Acc:HGNC:3720] | *FKBP4* | 5.37 | 0.0054 | Increased in 3rd trimester pregnant individuals |
| ENSG00000185627 | proteasome 26S subunit, non-ATPase 13 [Source:HGNC Symbol;Acc:HGNC:9558] | *PSMD13* | 5.31 | 0.0136 | Increased in 3rd trimester pregnant individuals |
| ENSG00000126581 | beclin 1 [Source:HGNC Symbol;Acc:HGNC:1034] | *BECN1* | 5.31 | 0.0094 | Increased in 3rd trimester pregnant individuals |
| ENSG00000144659 | solute carrier family 25 member 38 [Source:HGNC Symbol;Acc:HGNC:26054] | *SLC25A38* | 5.31 | 0.0219 | Increased in 3rd trimester pregnant individuals |
| ENSG00000115306 | spectrin beta, non-erythrocytic 1 [Source:HGNC Symbol;Acc:HGNC:11275] | *SPTBN1* | 5.28 | 0.0090 | Increased in 3rd trimester pregnant individuals |
| ENSG00000130522 | JunD proto-oncogene, AP-1 transcription factor subunit [Source:HGNC Symbol;Acc:HGNC:6206] | *JUND* | 5.28 | 0.0058 | Increased in 3rd trimester pregnant individuals |
| ENSG00000101004 | ninein like [Source:HGNC Symbol;Acc:HGNC:29163] | *NINL* | 5.26 | 0.0064 | Increased in 3rd trimester pregnant individuals |
| ENSG00000172380 | G protein subunit gamma 12 [Source:HGNC Symbol;Acc:HGNC:19663] | *GNG12* | 5.25 | 0.0164 | Increased in 3rd trimester pregnant individuals |
| ENSG00000107438 | PDZ and LIM domain 1 [Source:HGNC Symbol;Acc:HGNC:2067] | *PDLIM1* | 5.24 | 0.0228 | Increased in 3rd trimester pregnant individuals |
| ENSG00000142546 | nitric oxide synthase interacting protein [Source:HGNC Symbol;Acc:HGNC:17946] | *NOSIP* | 5.22 | 0.0411 | Increased in 3rd trimester pregnant individuals |
| ENSG00000143575 | HCLS1 associated protein X-1 [Source:HGNC Symbol;Acc:HGNC:16915] | *HAX1* | 5.18 | 0.0179 | Increased in 3rd trimester pregnant individuals |
| ENSG00000087274 | adducin 1 [Source:HGNC Symbol;Acc:HGNC:243] | *ADD1* | 5.16 | 0.0139 | Increased in 3rd trimester pregnant individuals |
| ENSG00000105220 | glucose-6-phosphate isomerase [Source:HGNC Symbol;Acc:HGNC:4458] | *GPI* | 5.14 | 0.0245 | Increased in 3rd trimester pregnant individuals |

| | | | | | |
|---|---|---|---|---|---|
| ENSG00000157500 | adaptor protein, phosphotyrosine interacting with PH domain and leucine zipper 1 [Source:HGNC Symbol;Acc:HGNC:24035] | *APPL1* | 5.13 | 0.0064 | Increased in 3rd trimester pregnant individuals |
| ENSG00000051108 | homocysteine inducible ER protein with ubiquitin like domain 1 [Source:HGNC Symbol;Acc:HGNC:13744] | *HERPUD1* | 5.13 | 0.0094 | Increased in 3rd trimester pregnant individuals |
| ENSG00000170759 | kinesin family member 5B [Source:HGNC Symbol;Acc:HGNC:6324] | *KIF5B* | 5.12 | 0.0064 | Increased in 3rd trimester pregnant individuals |
| ENSG00000091039 | oxysterol binding protein like 8 [Source:HGNC Symbol;Acc:HGNC:16396] | *OSBPL8* | 5.10 | 0.0064 | Increased in 3rd trimester pregnant individuals |
| ENSG00000137876 | ribosomal L24 domain containing 1 [Source:HGNC Symbol;Acc:HGNC:18479] | *RSL24D1* | 5.07 | 0.0058 | Increased in 3rd trimester pregnant individuals |
| ENSG00000115762 | pleckstrin homology domain containing B2 [Source:HGNC Symbol;Acc:HGNC:19236] | *PLEKHB2* | 5.06 | 0.0058 | Increased in 3rd trimester pregnant individuals |
| ENSG00000073350 | LLGL scribble cell polarity complex component 2 [Source:HGNC Symbol;Acc:HGNC:6629] | *LLGL2* | 5.04 | 0.0136 | Increased in 3rd trimester pregnant individuals |
| ENSG00000158528 | protein phosphatase 1 regulatory subunit 9A [Source:HGNC Symbol;Acc:HGNC:14946] | *PPP1R9A* | 5.03 | 0.0467 | Increased in 3rd trimester pregnant individuals |
| ENSG00000156587 | ubiquitin conjugating enzyme E2 L6 [Source:HGNC Symbol;Acc:HGNC:12490] | *UBE2L6* | 5.02 | 0.0179 | Increased in 3rd trimester pregnant individuals |
| ENSG00000188938 | family with sequence similarity 120A opposite strand [Source:HGNC Symbol;Acc:HGNC:23389] | *FAM120AOS* | 5.02 | 0.0365 | Increased in 3rd trimester pregnant individuals |
| ENSG00000005469 | carnitine O-octanoyltransferase [Source:HGNC Symbol;Acc:HGNC:2366] | *CROT* | 5.01 | 0.0058 | Increased in 3rd trimester pregnant individuals |
| ENSG00000138629 | ubiquitin like 7 [Source:HGNC Symbol;Acc:HGNC:28221] | *UBL7* | 5.00 | 0.0088 | Increased in 3rd trimester pregnant individuals |
| ENSG00000100823 | apurinic/apyrimidinic endodeoxyribonuclease 1 [Source:HGNC Symbol;Acc:HGNC:587] | *APEX1* | 4.99 | 0.0062 | Increased in 3rd trimester pregnant individuals |

| | | | | | |
|---|---|---|---|---|---|
| ENSG00000198682 | 3'-phosphoadenosine 5'-phosphosulfate synthase 2 [Source:HGNC Symbol;Acc:HGNC:8604] | *PAPSS2* | 4.99 | 0.0236 | Increased in 3rd trimester pregnant individuals |
| ENSG00000132002 | DnaJ heat shock protein family (Hsp40) member B1 [Source:HGNC Symbol;Acc:HGNC:5270] | *DNAJB1* | 4.99 | 0.0086 | Increased in 3rd trimester pregnant individuals |
| ENSG00000052795 | folliculin interacting protein 2 [Source:HGNC Symbol;Acc:HGNC:29280] | *FNIP2* | 4.98 | 0.0065 | Increased in 3rd trimester pregnant individuals |
| ENSG00000100897 | DDB1 and CUL4 associated factor 11 [Source:HGNC Symbol;Acc:HGNC:20258] | *DCAF11* | 4.98 | 0.0063 | Increased in 3rd trimester pregnant individuals |
| ENSG00000057149 | serpin family B member 3 [Source:HGNC Symbol;Acc:HGNC:10569] | *SERPINB3* | 4.97 | 0.0064 | Increased in 3rd trimester pregnant individuals |
| ENSG00000128789 | proteasome assembly chaperone 2 [Source:HGNC Symbol;Acc:HGNC:24929] | *PSMG2* | 4.97 | 0.0246 | Increased in 3rd trimester pregnant individuals |
| ENSG00000100603 | SNW domain containing 1 [Source:HGNC Symbol;Acc:HGNC:16696] | *SNW1* | 4.95 | 0.0188 | Increased in 3rd trimester pregnant individuals |
| ENSG00000111907 | TPD52 like 1 [Source:HGNC Symbol;Acc:HGNC:12006] | *TPD52L1* | 4.95 | 0.0239 | Increased in 3rd trimester pregnant individuals |
| ENSG00000198130 | 3-hydroxyisobutyryl-CoA hydrolase [Source:HGNC Symbol;Acc:HGNC:4908] | *HIBCH* | 4.94 | 0.0058 | Increased in 3rd trimester pregnant individuals |
| ENSG00000147400 | centrin 2 [Source:HGNC Symbol;Acc:HGNC:1867] | *CETN2* | 4.92 | 0.0139 | Increased in 3rd trimester pregnant individuals |
| ENSG00000230067 | heat shock protein family D (Hsp60) member 1 pseudogene 6 [Source:HGNC Symbol;Acc:HGNC:5267] | *HSPD1P6* | 4.91 | 0.0179 | Increased in 3rd trimester pregnant individuals |
| ENSG00000175220 | Rho GTPase activating protein 1 [Source:HGNC Symbol;Acc:HGNC:673] | *ARHGAP1* | 4.90 | 0.0065 | Increased in 3rd trimester pregnant individuals |
| ENSG00000122042 | ubiquitin like 3 [Source:HGNC Symbol;Acc:HGNC:12504] | *UBL3* | 4.89 | 0.0065 | Increased in 3rd trimester pregnant individuals |

| | | | | | |
|---|---|---|---|---|---|
| ENSG00000139055 | endoplasmic reticulum protein 27 [Source:HGNC Symbol;Acc:HGNC:26495] | *ERP27* | 4.88 | 0.0094 | Increased in 3rd trimester pregnant individuals |
| ENSG00000103254 | adenine nucleotide translocase lysine methyltransferase [Source:HGNC Symbol;Acc:HGNC:14152] | *ANTKMT* | 4.87 | 0.0228 | Increased in 3rd trimester pregnant individuals |
| ENSG00000140307 | general transcription factor IIA subunit 2 [Source:HGNC Symbol;Acc:HGNC:4647] | *GTF2A2* | 4.87 | 0.0133 | Increased in 3rd trimester pregnant individuals |
| ENSG00000023191 | ribonuclease/angiogenin inhibitor 1 [Source:HGNC Symbol;Acc:HGNC:10074] | *RNH1* | 4.86 | 0.0136 | Increased in 3rd trimester pregnant individuals |
| ENSG00000120705 | eukaryotic translation termination factor 1 [Source:HGNC Symbol;Acc:HGNC:3477] | *ETF1* | 4.85 | 0.0059 | Increased in 3rd trimester pregnant individuals |
| ENSG00000091164 | thioredoxin like 1 [Source:HGNC Symbol;Acc:HGNC:12436] | *TXNL1* | 4.85 | 0.0090 | Increased in 3rd trimester pregnant individuals |
| ENSG00000167699 | glyoxalase domain containing 4 [Source:HGNC Symbol;Acc:HGNC:14111] | *GLOD4* | 4.85 | 0.0136 | Increased in 3rd trimester pregnant individuals |
| ENSG00000167797 | cyclin dependent kinase 2 associated protein 2 [Source:HGNC Symbol;Acc:HGNC:30833] | *CDK2AP2* | 4.83 | 0.0139 | Increased in 3rd trimester pregnant individuals |
| ENSG00000146729 | nipsnap homolog 2 [Source:HGNC Symbol;Acc:HGNC:4179] | *NIPSNAP2* | 4.83 | 0.0480 | Increased in 3rd trimester pregnant individuals |
| ENSG00000139168 | zinc finger CCHC-type and RNA binding motif containing 1 [Source:HGNC Symbol;Acc:HGNC:29620] | *ZCRB1* | 4.82 | 0.0100 | Increased in 3rd trimester pregnant individuals |
| ENSG00000143799 | poly(ADP-ribose) polymerase 1 [Source:HGNC Symbol;Acc:HGNC:270] | *PARP1* | 4.81 | 0.0088 | Increased in 3rd trimester pregnant individuals |
| ENSG00000164654 | meiosis regulator for oocyte development [Source:HGNC Symbol;Acc:HGNC:21905] | *MIOS* | 4.81 | 0.0258 | Increased in 3rd trimester pregnant individuals |
| ENSG00000135245 | hypoxia inducible lipid droplet associated [Source:HGNC Symbol;Acc:HGNC:28859] | *HILPDA* | 4.79 | 0.0440 | Increased in 3rd trimester pregnant individuals |

| | | | | | |
|---|---|---|---|---|---|
| ENSG00000166562 | SEC11 homolog C, signal peptidase complex subunit [Source:HGNC Symbol;Acc:HGNC:23400] | *SEC11C* | 4.78 | 0.0179 | Increased in 3rd trimester pregnant individuals |
| ENSG00000185187 | single Ig and TIR domain containing [Source:HGNC Symbol;Acc:HGNC:30575] | *SIGIRR* | 4.77 | 0.0290 | Increased in 3rd trimester pregnant individuals |
| ENSG00000278053 | DExD-box helicase 52 [Source:HGNC Symbol;Acc:HGNC:20038] | *DDX52* | 4.77 | 0.0090 | Increased in 3rd trimester pregnant individuals |
| ENSG00000106399 | replication protein A3 [Source:HGNC Symbol;Acc:HGNC:10291] | *RPA3* | 4.77 | 0.0136 | Increased in 3rd trimester pregnant individuals |
| ENSG00000143321 | heparin binding growth factor [Source:HGNC Symbol;Acc:HGNC:4856] | *HDGF* | 4.76 | 0.0094 | Increased in 3rd trimester pregnant individuals |
| ENSG00000151500 | thymocyte nuclear protein 1 [Source:HGNC Symbol;Acc:HGNC:29560] | *THYN1* | 4.72 | 0.0133 | Increased in 3rd trimester pregnant individuals |
| ENSG00000065526 | spen family transcriptional repressor [Source:HGNC Symbol;Acc:HGNC:17575] | *SPEN* | 4.72 | 0.0344 | Increased in 3rd trimester pregnant individuals |
| ENSG00000177889 | ubiquitin conjugating enzyme E2 N [Source:HGNC Symbol;Acc:HGNC:12492] | *UBE2N* | 4.70 | 0.0114 | Increased in 3rd trimester pregnant individuals |
| ENSG00000011114 | BTB domain containing 7 [Source:HGNC Symbol;Acc:HGNC:18269] | *BTBD7* | 4.70 | 0.0116 | Increased in 3rd trimester pregnant individuals |
| ENSG00000197930 | endoplasmic reticulum oxidoreductase 1 alpha [Source:HGNC Symbol;Acc:HGNC:13280] | *ERO1A* | 4.70 | 0.0364 | Increased in 3rd trimester pregnant individuals |
| ENSG00000116977 | galectin 8 [Source:HGNC Symbol;Acc:HGNC:6569] | *LGALS8* | 4.68 | 0.0133 | Increased in 3rd trimester pregnant individuals |
| ENSG00000243449 | chromosome 4 open reading frame 48 [Source:HGNC Symbol;Acc:HGNC:34437] | *C4orf48* | 4.66 | 0.0136 | Increased in 3rd trimester pregnant individuals |
| ENSG00000099330 | occludin/ELL domain containing 1 [Source:HGNC Symbol;Acc:HGNC:26221] | *OCEL1* | 4.66 | 0.0364 | Increased in 3rd trimester pregnant individuals |

| | | | | | |
|---|---|---|---|---|---|
| ENSG00000215021 | prohibitin 2 [Source:HGNC Symbol;Acc:HGNC:30306] | *PHB2* | 4.65 | 0.0188 | Increased in 3rd trimester pregnant individuals |
| ENSG00000049283 | epsin 3 [Source:HGNC Symbol;Acc:HGNC:18235] | *EPN3* | 4.64 | 0.0463 | Increased in 3rd trimester pregnant individuals |
| ENSG00000184432 | COPI coat complex subunit beta 2 [Source:HGNC Symbol;Acc:HGNC:2232] | *COPB2* | 4.62 | 0.0235 | Increased in 3rd trimester pregnant individuals |
| ENSG00000146038 | doublecortin domain containing 2 [Source:HGNC Symbol;Acc:HGNC:18141] | *DCDC2* | 4.61 | 0.0139 | Increased in 3rd trimester pregnant individuals |
| ENSG00000126870 | dynein 2 intermediate chain 1 [Source:HGNC Symbol;Acc:HGNC:21862] | *DYNC2I1* | 4.59 | 0.0323 | Increased in 3rd trimester pregnant individuals |
| ENSG00000128626 | mitochondrial ribosomal protein S12 [Source:HGNC Symbol;Acc:HGNC:10380] | *MRPS12* | 4.58 | 0.0164 | Increased in 3rd trimester pregnant individuals |
| ENSG00000169032 | mitogen-activated protein kinase kinase 1 [Source:HGNC Symbol;Acc:HGNC:6840] | *MAP2K1* | 4.58 | 0.0154 | Increased in 3rd trimester pregnant individuals |
| ENSG00000128487 | sperm antigen with calponin homology and coiled-coil domains 1 [Source:HGNC Symbol;Acc:HGNC:30615] | *SPECC1* | 4.58 | 0.0136 | Increased in 3rd trimester pregnant individuals |
| ENSG00000188529 | serine and arginine rich splicing factor 10 [Source:HGNC Symbol;Acc:HGNC:16713] | *SRSF10* | 4.57 | 0.0332 | Increased in 3rd trimester pregnant individuals |
| ENSG00000160570 | death effector domain containing 2 [Source:HGNC Symbol;Acc:HGNC:24450] | *DEDD2* | 4.57 | 0.0377 | Increased in 3rd trimester pregnant individuals |
| ENSG00000087077 | thyroid hormone receptor interactor 6 [Source:HGNC Symbol;Acc:HGNC:12311] | *TRIP6* | 4.56 | 0.0445 | Increased in 3rd trimester pregnant individuals |
| ENSG00000170291 | elongator acetyltransferase complex subunit 5 [Source:HGNC Symbol;Acc:HGNC:30617] | *ELP5* | 4.55 | 0.0476 | Increased in 3rd trimester pregnant individuals |
| ENSG00000122203 | KIAA1191 [Source:HGNC Symbol;Acc:HGNC:29209] | *KIAA1191* | 4.54 | 0.0136 | Increased in 3rd trimester pregnant individuals |

| | | | | | |
|---|---|---|---|---|---|
| ENSG00000076864 | RAP1 GTPase activating protein [Source:HGNC Symbol;Acc:HGNC:9858] | *RAP1GAP* | 4.54 | 0.0188 | Increased in 3rd trimester pregnant individuals |
| **ENSG00000007944*** | **myosin regulatory light chain interacting protein [Source:HGNC Symbol;Acc:HGNC:21155]** | **MYLIP** | **4.53** | **0.0365** | **Increased in 3rd trimester pregnant individuals** |
| ENSG00000116574 | ras homolog family member U [Source:HGNC Symbol;Acc:HGNC:17794] | *RHOU* | 4.53 | 0.0100 | Increased in 3rd trimester pregnant individuals |
| ENSG00000198730 | CTR9 homolog, Paf1/RNA polymerase II complex component [Source:HGNC Symbol;Acc:HGNC:16850] | *CTR9* | 4.53 | 0.0195 | Increased in 3rd trimester pregnant individuals |
| ENSG00000139726 | density regulated re-initiation and release factor [Source:HGNC Symbol;Acc:HGNC:2769] | *DENR* | 4.51 | 0.0439 | Increased in 3rd trimester pregnant individuals |
| ENSG00000136758 | YME1 like 1 ATPase [Source:HGNC Symbol;Acc:HGNC:12843] | *YME1L1* | 4.50 | 0.0179 | Increased in 3rd trimester pregnant individuals |
| ENSG00000007520 | TSR3 ribosome maturation factor [Source:HGNC Symbol;Acc:HGNC:14175] | *TSR3* | 4.50 | 0.0229 | Increased in 3rd trimester pregnant individuals |
| ENSG00000165322 | Rho GTPase activating protein 12 [Source:HGNC Symbol;Acc:HGNC:16348] | *ARHGAP12* | 4.49 | 0.0179 | Increased in 3rd trimester pregnant individuals |
| ENSG00000089737 | DEAD-box helicase 24 [Source:HGNC Symbol;Acc:HGNC:13266] | *DDX24* | 4.48 | 0.0159 | Increased in 3rd trimester pregnant individuals |
| **ENSG00000108244*** | **keratin 23 [Source:HGNC Symbol;Acc:HGNC:6438]** | **KRT23** | **4.47** | **0.0452** | **Increased in 3rd trimester pregnant individuals** |
| ENSG00000165813 | coiled-coil domain containing 186 [Source:HGNC Symbol;Acc:HGNC:24349] | *CCDC186* | 4.46 | 0.0179 | Increased in 3rd trimester pregnant individuals |
| ENSG00000126878 | allograft inflammatory factor 1 like [Source:HGNC Symbol;Acc:HGNC:28904] | *AIF1L* | 4.45 | 0.0150 | Increased in 3rd trimester pregnant individuals |
| ENSG00000044115 | catenin alpha 1 [Source:HGNC Symbol;Acc:HGNC:2509] | *CTNNA1* | 4.44 | 0.0237 | Increased in 3rd trimester pregnant individuals |

| | | | | | |
|---|---|---|---|---|---|
| ENSG00000119878 | CXXC repeat containing interactor of PDZ3 domain [Source:HGNC Symbol;Acc:HGNC:14312] | *CRIPT* | 4.43 | 0.0159 | Increased in 3rd trimester pregnant individuals |
| ENSG00000100865 | cyclin dependent kinase 2 interacting protein [Source:HGNC Symbol;Acc:HGNC:23789] | *CINP* | 4.43 | 0.0250 | Increased in 3rd trimester pregnant individuals |
| ENSG00000126254 | RNA binding motif protein 42 [Source:HGNC Symbol;Acc:HGNC:28117] | *RBM42* | 4.42 | 0.0467 | Increased in 3rd trimester pregnant individuals |
| ENSG00000122545 | septin 7 [Source:HGNC Symbol;Acc:HGNC:1717] | *SEPTIN7* | 4.41 | 0.0154 | Increased in 3rd trimester pregnant individuals |
| ENSG00000104964 | TLE family member 5, transcriptional modulator [Source:HGNC Symbol;Acc:HGNC:307] | *TLE5* | 4.40 | 0.0179 | Increased in 3rd trimester pregnant individuals |
| ENSG00000119431 | haloacid dehalogenase like hydrolase domain containing 3 [Source:HGNC Symbol;Acc:HGNC:28171] | *HDHD3* | 4.40 | 0.0463 | Increased in 3rd trimester pregnant individuals |
| ENSG00000135447 | protein phosphatase 1 regulatory inhibitor subunit 1A [Source:HGNC Symbol;Acc:HGNC:9286] | *PPP1R1A* | 4.40 | 0.0252 | Increased in 3rd trimester pregnant individuals |
| ENSG00000151893 | CDK2 associated cullin domain 1 [Source:HGNC Symbol;Acc:HGNC:23727] | *CACUL1* | 4.38 | 0.0476 | Increased in 3rd trimester pregnant individuals |
| ENSG00000153130 | short coiled-coil protein [Source:HGNC Symbol;Acc:HGNC:20335] | *SCOC* | 4.38 | 0.0370 | Increased in 3rd trimester pregnant individuals |
| ENSG00000168092 | platelet activating factor acetylhydrolase 1b catalytic subunit 2 [Source:HGNC Symbol;Acc:HGNC:8575] | *PAFAH1B2* | 4.37 | 0.0323 | Increased in 3rd trimester pregnant individuals |
| ENSG00000128849 | cingulin like 1 [Source:HGNC Symbol;Acc:HGNC:25931] | *CGNL1* | 4.36 | 0.0219 | Increased in 3rd trimester pregnant individuals |
| ENSG00000101182 | proteasome 20S subunit alpha 7 [Source:HGNC Symbol;Acc:HGNC:9536] | *PSMA7* | 4.36 | 0.0225 | Increased in 3rd trimester pregnant individuals |
| ENSG00000182606 | trafficking kinesin protein 1 [Source:HGNC Symbol;Acc:HGNC:29947] | *TRAK1* | 4.35 | 0.0164 | Increased in 3rd trimester pregnant individuals |

| | | | | | |
|---|---|---|---|---|---|
| ENSG00000167986 | damage specific DNA binding protein 1 [Source:HGNC Symbol;Acc:HGNC:2717] | *DDB1* | 4.34 | 0.0477 | Increased in 3rd trimester pregnant individuals |
| ENSG00000242372 | eukaryotic translation initiation factor 6 [Source:HGNC Symbol;Acc:HGNC:6159] | *EIF6* | 4.34 | 0.0170 | Increased in 3rd trimester pregnant individuals |
| ENSG00000109332 | ubiquitin conjugating enzyme E2 D3 [Source:HGNC Symbol;Acc:HGNC:12476] | *UBE2D3* | 4.34 | 0.0143 | Increased in 3rd trimester pregnant individuals |
| ENSG00000116560 | splicing factor proline and glutamine rich [Source:HGNC Symbol;Acc:HGNC:10774] | *SFPQ* | 4.34 | 0.0224 | Increased in 3rd trimester pregnant individuals |
| ENSG00000119938 | protein phosphatase 1 regulatory subunit 3C [Source:HGNC Symbol;Acc:HGNC:9293] | *PPP1R3C* | 4.33 | 0.0170 | Increased in 3rd trimester pregnant individuals |
| ENSG00000023228 | NADH:ubiquinone oxidoreductase core subunit S1 [Source:HGNC Symbol;Acc:HGNC:7707] | *NDUFS1* | 4.32 | 0.0323 | Increased in 3rd trimester pregnant individuals |
| ENSG00000113312 | tetratricopeptide repeat domain 1 [Source:HGNC Symbol;Acc:HGNC:12391] | *TTC1* | 4.31 | 0.0252 | Increased in 3rd trimester pregnant individuals |
| ENSG00000169398 | protein tyrosine kinase 2 [Source:HGNC Symbol;Acc:HGNC:9611] | *PTK2* | 4.29 | 0.0179 | Increased in 3rd trimester pregnant individuals |
| ENSG00000146731 | chaperonin containing TCP1 subunit 6A [Source:HGNC Symbol;Acc:HGNC:1620] | *CCT6A* | 4.29 | 0.0421 | Increased in 3rd trimester pregnant individuals |
| ENSG00000154174 | translocase of outer mitochondrial membrane 70 [Source:HGNC Symbol;Acc:HGNC:11985] | *TOMM70* | 4.29 | 0.0496 | Increased in 3rd trimester pregnant individuals |
| ENSG00000143222 | ubiquitin-fold modifier conjugating enzyme 1 [Source:HGNC Symbol;Acc:HGNC:26941] | *UFC1* | 4.29 | 0.0159 | Increased in 3rd trimester pregnant individuals |
| ENSG00000106628 | DNA polymerase delta 2, accessory subunit [Source:HGNC Symbol;Acc:HGNC:9176] | *POLD2* | 4.27 | 0.0239 | Increased in 3rd trimester pregnant individuals |
| ENSG00000175581 | mitochondrial ribosomal protein L48 [Source:HGNC Symbol;Acc:HGNC:16653] | *MRPL48* | 4.27 | 0.0448 | Increased in 3rd trimester pregnant individuals |

| | | | | | |
|---|---|---|---|---|---|
| ENSG00000167770 | OTU deubiquitinase, ubiquitin aldehyde binding 1 [Source:HGNC Symbol;Acc:HGNC:23077] | *OTUB1* | 4.27 | 0.0344 | Increased in 3rd trimester pregnant individuals |
| ENSG00000126756 | ubiquitously expressed prefoldin like chaperone [Source:HGNC Symbol;Acc:HGNC:12641] | *UXT* | 4.26 | 0.0219 | Increased in 3rd trimester pregnant individuals |
| ENSG00000164134 | N-alpha-acetyltransferase 15, NatA auxiliary subunit [Source:HGNC Symbol;Acc:HGNC:30782] | *NAA15* | 4.25 | 0.0421 | Increased in 3rd trimester pregnant individuals |
| ENSG00000226950 | differentiation antagonizing non-protein coding RNA [Source:HGNC Symbol;Acc:HGNC:28964] | *DANCR* | 4.25 | 0.0195 | Increased in 3rd trimester pregnant individuals |
| ENSG00000120686 | ubiquitin fold modifier 1 [Source:HGNC Symbol;Acc:HGNC:20597] | *UFM1* | 4.24 | 0.0188 | Increased in 3rd trimester pregnant individuals |
| ENSG00000107960 | STN1 subunit of CST complex [Source:HGNC Symbol;Acc:HGNC:26200] | *STN1* | 4.24 | 0.0179 | Increased in 3rd trimester pregnant individuals |
| ENSG00000104969 | small glutamine rich tetratricopeptide repeat co-chaperone alpha [Source:HGNC Symbol;Acc:HGNC:10819] | *SGTA* | 4.24 | 0.0435 | Increased in 3rd trimester pregnant individuals |
| ENSG00000165525 | nuclear export mediator factor [Source:HGNC Symbol;Acc:HGNC:10663] | *NEMF* | 4.24 | 0.0496 | Increased in 3rd trimester pregnant individuals |
| ENSG00000198791 | CCR4-NOT transcription complex subunit 7 [Source:HGNC Symbol;Acc:HGNC:14101] | *CNOT7* | 4.22 | 0.0148 | Increased in 3rd trimester pregnant individuals |
| ENSG00000137574 | trimethylguanosine synthase 1 [Source:HGNC Symbol;Acc:HGNC:17843] | *TGS1* | 4.22 | 0.0323 | Increased in 3rd trimester pregnant individuals |
| ENSG00000107929 | La ribonucleoprotein 4B [Source:HGNC Symbol;Acc:HGNC:28987] | *LARP4B* | 4.22 | 0.0477 | Increased in 3rd trimester pregnant individuals |
| ENSG00000102054 | RB binding protein 7, chromatin remodeling factor [Source:HGNC Symbol;Acc:HGNC:9890] | *RBBP7* | 4.22 | 0.0320 | Increased in 3rd trimester pregnant individuals |
| ENSG00000166266 | cullin 5 [Source:HGNC Symbol;Acc:HGNC:2556] | *CUL5* | 4.21 | 0.0433 | Increased in 3rd trimester pregnant individuals |

| | | | | | |
|---|---|---|---|---|---|
| ENSG00000065427 | lysyl-tRNA synthetase 1 [Source:HGNC Symbol;Acc:HGNC:6215] | *KARS1* | 4.20 | 0.0439 | Increased in 3rd trimester pregnant individuals |
| ENSG00000102572 | serine/threonine kinase 24 [Source:HGNC Symbol;Acc:HGNC:11403] | *STK24* | 4.20 | 0.0250 | Increased in 3rd trimester pregnant individuals |
| ENSG00000090013 | biliverdin reductase B [Source:HGNC Symbol;Acc:HGNC:1063] | *BLVRB* | 4.18 | 0.0341 | Increased in 3rd trimester pregnant individuals |
| ENSG00000272398 | CD24 molecule [Source:HGNC Symbol;Acc:HGNC:1645] | *CD24* | 4.17 | 0.0323 | Increased in 3rd trimester pregnant individuals |
| ENSG00000141401 | inositol monophosphatase 2 [Source:HGNC Symbol;Acc:HGNC:6051] | *IMPA2* | 4.13 | 0.0440 | Increased in 3rd trimester pregnant individuals |
| ENSG00000144224 | UBX domain protein 4 [Source:HGNC Symbol;Acc:HGNC:14860] | *UBXN4* | 4.13 | 0.0344 | Increased in 3rd trimester pregnant individuals |
| ENSG00000099260 | palmdelphin [Source:HGNC Symbol;Acc:HGNC:15846] | *PALMD* | 4.13 | 0.0298 | Increased in 3rd trimester pregnant individuals |
| ENSG00000151276 | membrane associated guanylate kinase, WW and PDZ domain containing 1 [Source:HGNC Symbol;Acc:HGNC:946] | *MAGI1* | 4.12 | 0.0287 | Increased in 3rd trimester pregnant individuals |
| ENSG00000100697 | dicer 1, ribonuclease III [Source:HGNC Symbol;Acc:HGNC:17098] | *DICER1* | 4.12 | 0.0317 | Increased in 3rd trimester pregnant individuals |
| ENSG00000143499 | SET and MYND domain containing 2 [Source:HGNC Symbol;Acc:HGNC:20982] | *SMYD2* | 4.12 | 0.0416 | Increased in 3rd trimester pregnant individuals |
| ENSG00000213719 | chloride intracellular channel 1 [Source:HGNC Symbol;Acc:HGNC:2062] | *CLIC1* | 4.11 | 0.0237 | Increased in 3rd trimester pregnant individuals |
| ENSG00000189060 | H1.0 linker histone [Source:HGNC Symbol;Acc:HGNC:4714] | *H1-0* | 4.10 | 0.0440 | Increased in 3rd trimester pregnant individuals |
| ENSG00000138175 | ADP ribosylation factor like GTPase 3 [Source:HGNC Symbol;Acc:HGNC:694] | *ARL3* | 4.10 | 0.0476 | Increased in 3rd trimester pregnant individuals |

| | | | | | |
|---|---|---|---|---|---|
| ENSG00000115364 | mitochondrial ribosomal protein L19 [Source:HGNC Symbol;Acc:HGNC:14052] | *MRPL19* | 4.09 | 0.0195 | Increased in 3rd trimester pregnant individuals |
| ENSG00000110048 | oxysterol binding protein [Source:HGNC Symbol;Acc:HGNC:8503] | *OSBP* | 4.08 | 0.0448 | Increased in 3rd trimester pregnant individuals |
| ENSG00000119720 | NRDE-2, necessary for RNA interference, domain containing [Source:HGNC Symbol;Acc:HGNC:20186] | *NRDE2* | 4.07 | 0.0298 | Increased in 3rd trimester pregnant individuals |
| ENSG00000125863 | MKKS centrosomal shuttling protein [Source:HGNC Symbol;Acc:HGNC:7108] | *MKKS* | 4.07 | 0.0364 | Increased in 3rd trimester pregnant individuals |
| ENSG00000124098 | family with sequence similarity 210 member B [Source:HGNC Symbol;Acc:HGNC:16102] | *FAM210B* | 4.05 | 0.0401 | Increased in 3rd trimester pregnant individuals |
| ENSG00000077684 | jade family PHD finger 1 [Source:HGNC Symbol;Acc:HGNC:30027] | *JADE1* | 4.05 | 0.0468 | Increased in 3rd trimester pregnant individuals |
| ENSG00000134744 | terminal uridylyl transferase 4 [Source:HGNC Symbol;Acc:HGNC:28981] | *TUT4* | 4.03 | 0.0159 | Increased in 3rd trimester pregnant individuals |
| ENSG00000175701 | mitoregulin [Source:HGNC Symbol;Acc:HGNC:27339] | *MTLN* | 4.02 | 0.0434 | Increased in 3rd trimester pregnant individuals |
| ENSG00000166750 | schlafen family member 5 [Source:HGNC Symbol;Acc:HGNC:28286] | *SLFN5* | 4.01 | 0.0179 | Increased in 3rd trimester pregnant individuals |
| ENSG00000122482 | zinc finger protein 644 [Source:HGNC Symbol;Acc:HGNC:29222] | *ZNF644* | 3.98 | 0.0496 | Increased in 3rd trimester pregnant individuals |
| ENSG00000116754 | serine and arginine rich splicing factor 11 [Source:HGNC Symbol;Acc:HGNC:10782] | *SRSF11* | 3.97 | 0.0344 | Increased in 3rd trimester pregnant individuals |
| ENSG00000065882 | TBC1 domain family member 1 [Source:HGNC Symbol;Acc:HGNC:11578] | *TBC1D1* | 3.92 | 0.0496 | Increased in 3rd trimester pregnant individuals |
| ENSG00000122705 | clathrin light chain A [Source:HGNC Symbol;Acc:HGNC:2090] | *CLTA* | 3.91 | 0.0380 | Increased in 3rd trimester pregnant individuals |

| | | | | | |
|---|---|---|---|---|---|
| ENSG00000117118 | succinate dehydrogenase complex iron sulfur subunit B [Source:HGNC Symbol;Acc:HGNC:10681] | *SDHB* | 3.91 | 0.0406 | Increased in 3rd trimester pregnant individuals |
| ENSG00000115128 | splicing factor 3b subunit 6 [Source:HGNC Symbol;Acc:HGNC:30096] | *SF3B6* | 3.90 | 0.0427 | Increased in 3rd trimester pregnant individuals |
| ENSG00000124802 | eukaryotic translation elongation factor 1 epsilon 1 [Source:HGNC Symbol;Acc:HGNC:3212] | *EEF1E1* | 3.84 | 0.0358 | Increased in 3rd trimester pregnant individuals |
| ENSG00000163170 | bolA family member 3 [Source:HGNC Symbol;Acc:HGNC:24415] | *BOLA3* | 3.84 | 0.0496 | Increased in 3rd trimester pregnant individuals |
| ENSG00000105357 | myosin heavy chain 14 [Source:HGNC Symbol;Acc:HGNC:23212] | *MYH14* | 3.84 | 0.0365 | Increased in 3rd trimester pregnant individuals |
| ENSG00000124380 | small nuclear ribonucleoprotein U4/U6.U5 subunit 27 [Source:HGNC Symbol;Acc:HGNC:30240] | *SNRNP27* | 3.82 | 0.0346 | Increased in 3rd trimester pregnant individuals |
| ENSG00000173473 | SWI/SNF related, matrix associated, actin dependent regulator of chromatin subfamily c member 1 [Source:HGNC Symbol;Acc:HGNC:11104] | *SMARCC1* | 3.82 | 0.0377 | Increased in 3rd trimester pregnant individuals |
| ENSG00000102144 | phosphoglycerate kinase 1 [Source:HGNC Symbol;Acc:HGNC:8896] | *PGK1* | 3.81 | 0.0404 | Increased in 3rd trimester pregnant individuals |
| ENSG00000141753 | insulin like growth factor binding protein 4 [Source:HGNC Symbol;Acc:HGNC:5473] | *IGFBP4* | 3.80 | 0.0492 | Increased in 3rd trimester pregnant individuals |
| ENSG00000138385 | small RNA binding exonuclease protection factor La [Source:HGNC Symbol;Acc:HGNC:11316] | *SSB* | 3.79 | 0.0411 | Increased in 3rd trimester pregnant individuals |
| ENSG00000090989 | exocyst complex component 1 [Source:HGNC Symbol;Acc:HGNC:30380] | *EXOC1* | 3.78 | 0.0439 | Increased in 3rd trimester pregnant individuals |
| ENSG00000197312 | DNA damage inducible 1 homolog 2 [Source:HGNC Symbol;Acc:HGNC:24578] | *DDI2* | 3.77 | 0.0402 | Increased in 3rd trimester pregnant individuals |
| ENSG00000131051 | RNA binding motif protein 39 [Source:HGNC Symbol;Acc:HGNC:15923] | *RBM39* | 3.71 | 0.0463 | Increased in 3rd trimester pregnant individuals |

| | | | | | |
|---|---|---|---|---|---|
| ENSG00000137288 | ubiquinol-cytochrome c reductase complex assembly factor 2 [Source:HGNC Symbol;Acc:HGNC:21237] | *UQCC2* | 3.70 | 0.0496 | Increased in 3rd trimester pregnant individuals |
| ENSG00000242485 | mitochondrial ribosomal protein L20 [Source:HGNC Symbol;Acc:HGNC:14478] | *MRPL20* | 3.70 | 0.0476 | Increased in 3rd trimester pregnant individuals |
| ENSG00000179271 | GADD45G interacting protein 1 [Source:HGNC Symbol;Acc:HGNC:29996] | *GADD45GIP1* | 3.69 | 0.0439 | Increased in 3rd trimester pregnant individuals |
| ENSG00000161533 | acyl-CoA oxidase 1 [Source:HGNC Symbol;Acc:HGNC:119] | *ACOX1* | 3.69 | 0.0317 | Increased in 3rd trimester pregnant individuals |
| ENSG00000278311 | gametogenetin binding protein 2 [Source:HGNC Symbol;Acc:HGNC:19357] | *GGNBP2* | 3.69 | 0.0277 | Increased in 3rd trimester pregnant individuals |
| ENSG00000084090 | StAR related lipid transfer domain containing 7 [Source:HGNC Symbol;Acc:HGNC:18063] | *STARD7* | 3.68 | 0.0496 | Increased in 3rd trimester pregnant individuals |
| ENSG00000165119 | heterogeneous nuclear ribonucleoprotein K [Source:HGNC Symbol;Acc:HGNC:5044] | *HNRNPK* | 3.67 | 0.0345 | Increased in 3rd trimester pregnant individuals |
| ENSG00000069020 | microtubule associated serine/threonine kinase family member 4 [Source:HGNC Symbol;Acc:HGNC:19037] | *MAST4* | 3.62 | 0.0476 | Increased in 3rd trimester pregnant individuals |
| ENSG00000114439 | BBX high mobility group box domain containing [Source:HGNC Symbol;Acc:HGNC:14422] | *BBX* | 3.56 | 0.0290 | Increased in 3rd trimester pregnant individuals |
| ENSG00000224877 | NADH:ubiquinone oxidoreductase complex assembly factor 8 [Source:HGNC Symbol;Acc:HGNC:33551] | *NDUFAF8* | 3.47 | 0.0331 | Increased in 3rd trimester pregnant individuals |
| ENSG00000110958 | prostaglandin E synthase 3 [Source:HGNC Symbol;Acc:HGNC:16049] | *PTGES3* | 3.42 | 0.0476 | Increased in 3rd trimester pregnant individuals |
| ENSG00000153006 | SREK1 interacting protein 1 [Source:HGNC Symbol;Acc:HGNC:26716] | *SREK1IP1* | 3.32 | 0.0411 | Increased in 3rd trimester pregnant individuals |
| ENSG00000174720 | La ribonucleoprotein 7, transcriptional regulator [Source:HGNC Symbol;Acc:HGNC:24912] | *LARP7* | 3.27 | 0.0440 | Increased in 3rd trimester pregnant individuals |

| | | | | | |
|---|---|---|---|---|---|
| ENSG00000051620 | heme binding protein 2 [Source:HGNC Symbol;Acc:HGNC:15716] | *HEBP2* | 3.25 | 0.0439 | Increased in 3rd trimester pregnant individuals |
| ENSG00000198168 | small VCP interacting protein [Source:HGNC Symbol;Acc:HGNC:25238] | *SVIP* | 3.09 | 0.0496 | Increased in 3rd trimester pregnant individuals |
| ENSG00000089220 | phosphatidylethanolamine binding protein 1 [Source:HGNC Symbol;Acc:HGNC:8630] | *PEBP1* | 0.81 | 0.0411 | Increased in 3rd trimester pregnant individuals |
| ENSG00000169738 | dicarbonyl and L-xylulose reductase [Source:HGNC Symbol;Acc:HGNC:18985] | *DCXR* | -1.23 | 0.0298 | Decreased in 3rd trimester pregnant individuals |
| ENSG00000087086 | ferritin light chain [Source:HGNC Symbol;Acc:HGNC:3999] | *FTL* | -1.48 | 3.37E-36 | Decreased in 3rd trimester pregnant individuals |
| ENSG00000203396 | WDR45-like (WDR45L) pseudogene | *ENSG00000203396* | -2.54 | 7.29E-16 | Decreased in 3rd trimester pregnant individuals |
| ENSG00000132535 | discs large MAGUK scaffold protein 4 [Source:HGNC Symbol;Acc:HGNC:2903] | *DLG4* | -3.00 | 0.0377 | Decreased in 3rd trimester pregnant individuals |
| ***2nd trimester pregnant pools vs. 3rd trimester pregnant individuals*** | | | | | |
| ENSG00000213995 | NAD(P)HX dehydratase [Source:HGNC Symbol;Acc:HGNC:25576] | *NAXD* | -5.90 | 0.0022 | Decreased in 3rd trimester pregnant individuals |
| ENSG00000177674 | angiotensin II receptor associated protein [Source:HGNC Symbol;Acc:HGNC:13539] | *AGTRAP* | -5.78 | 0.0072 | Decreased in 3rd trimester pregnant individuals |
| ENSG00000125618 | paired box 8 [Source:HGNC Symbol;Acc:HGNC:8622] | *PAX8* | -5.75 | 0.0072 | Decreased in 3rd trimester pregnant individuals |
| ENSG00000179178 | transmembrane protein 125 [Source:HGNC Symbol;Acc:HGNC:28275] | *TMEM125* | -5.72 | 0.0104 | Decreased in 3rd trimester pregnant individuals |
| ENSG00000117676 | ribosomal protein S6 kinase A1 [Source:HGNC Symbol;Acc:HGNC:10430] | *RPS6KA1* | -5.67 | 0.0100 | Decreased in 3rd trimester pregnant individuals |

| | | | | | |
|---|---|---|---|---|---|
| ENSG00000108592 | FtsJ RNA 2'-O-methyltransferase 3 [Source:HGNC Symbol;Acc:HGNC:17136] | *FTSJ3* | -5.67 | 0.0100 | Decreased in 3rd trimester pregnant individuals |
| ENSG00000244734 | hemoglobin subunit beta [Source:HGNC Symbol;Acc:HGNC:4827] | *HBB* | -5.61 | 0.0169 | Decreased in 3rd trimester pregnant individuals |
| ENSG00000131779 | peroxisomal biogenesis factor 11 beta [Source:HGNC Symbol;Acc:HGNC:8853] | *PEX11B* | -5.59 | 0.0100 | Decreased in 3rd trimester pregnant individuals |
| ENSG00000032389 | EARP complex and GARP complex interacting protein 1 [Source:HGNC Symbol;Acc:HGNC:12383] | *EIPR1* | -5.54 | 0.0072 | Decreased in 3rd trimester pregnant individuals |
| ENSG00000182810 | DEAD-box helicase 28 [Source:HGNC Symbol;Acc:HGNC:17330] | *DDX28* | -5.51 | 0.0104 | Decreased in 3rd trimester pregnant individuals |
| ENSG00000110063 | decapping enzyme, scavenger [Source:HGNC Symbol;Acc:HGNC:29812] | *DCPS* | -5.42 | 0.0100 | Decreased in 3rd trimester pregnant individuals |
| ENSG00000110931 | calcium/calmodulin dependent protein kinase kinase 2 [Source:HGNC Symbol;Acc:HGNC:1470] | *CAMKK2* | -5.42 | 0.0100 | Decreased in 3rd trimester pregnant individuals |
| ENSG00000175287 | phytanoyl-CoA dioxygenase domain containing 1 [Source:HGNC Symbol;Acc:HGNC:23396] | *PHYHD1* | -5.39 | 0.0100 | Decreased in 3rd trimester pregnant individuals |
| ENSG00000169696 | ASPSCR1 tether for SLC2A4, UBX domain containing [Source:HGNC Symbol;Acc:HGNC:13825] | *ASPSCR1* | -5.36 | 0.0072 | Decreased in 3rd trimester pregnant individuals |
| ENSG00000184903 | inner mitochondrial membrane peptidase subunit 2 [Source:HGNC Symbol;Acc:HGNC:14598] | *IMMP2L* | -5.35 | 0.0100 | Decreased in 3rd trimester pregnant individuals |
| ENSG00000105854 | paraoxonase 2 [Source:HGNC Symbol;Acc:HGNC:9205] | *PON2* | -5.24 | 0.0127 | Decreased in 3rd trimester pregnant individuals |
| ENSG00000054611 | TBC1 domain family member 22A [Source:HGNC Symbol;Acc:HGNC:1309] | *TBC1D22A* | -5.21 | 0.0124 | Decreased in 3rd trimester pregnant individuals |
| ENSG00000182541 | LIM domain kinase 2 [Source:HGNC Symbol;Acc:HGNC:6614] | *LIMK2* | -5.14 | 0.0072 | Decreased in 3rd trimester pregnant individuals |

| | | | | | |
|---|---|---|---|---|---|
| ENSG00000164978 | nudix hydrolase 2 [Source:HGNC Symbol;Acc:HGNC:8049] | *NUDT2* | -5.14 | 0.0169 | Decreased in 3rd trimester pregnant individuals |
| ENSG00000176533 | G protein subunit gamma 7 [Source:HGNC Symbol;Acc:HGNC:4410] | *GNG7* | -5.11 | 0.0100 | Decreased in 3rd trimester pregnant individuals |
| ENSG00000116141 | microtubule affinity regulating kinase 1 [Source:HGNC Symbol;Acc:HGNC:6896] | *MARK1* | -5.09 | 0.0100 | Decreased in 3rd trimester pregnant individuals |
| ENSG00000100612 | dehydrogenase/reductase 7 [Source:HGNC Symbol;Acc:HGNC:21524] | *DHRS7* | -5.08 | 0.0196 | Decreased in 3rd trimester pregnant individuals |
| ENSG00000177105 | ras homolog family member G [Source:HGNC Symbol;Acc:HGNC:672] | *RHOG* | -5.08 | 0.0100 | Decreased in 3rd trimester pregnant individuals |
| ENSG00000124164 | VAMP associated protein B and C [Source:HGNC Symbol;Acc:HGNC:12649] | *VAPB* | -5.08 | 0.0100 | Decreased in 3rd trimester pregnant individuals |
| ENSG00000110871 | coenzyme Q5, methyltransferase [Source:HGNC Symbol;Acc:HGNC:28722] | *COQ5* | -5.07 | 0.0121 | Decreased in 3rd trimester pregnant individuals |
| ENSG00000100139 | MICAL like 1 [Source:HGNC Symbol;Acc:HGNC:29804] | *MICALL1* | -5.05 | 0.0100 | Decreased in 3rd trimester pregnant individuals |
| ENSG00000108798 | ABI family member 3 [Source:HGNC Symbol;Acc:HGNC:29859] | *ABI3* | -5.05 | 0.0100 | Decreased in 3rd trimester pregnant individuals |
| ENSG00000155465 | solute carrier family 7 member 7 [Source:HGNC Symbol;Acc:HGNC:11065] | *SLC7A7* | -5.03 | 0.0111 | Decreased in 3rd trimester pregnant individuals |
| ENSG00000110514 | MAP kinase activating death domain [Source:HGNC Symbol;Acc:HGNC:6766] | *MADD* | -5.03 | 0.0196 | Decreased in 3rd trimester pregnant individuals |
| ENSG00000151502 | VPS26, retromer complex component B [Source:HGNC Symbol;Acc:HGNC:28119] | *VPS26B* | -5.02 | 0.0100 | Decreased in 3rd trimester pregnant individuals |
| ENSG00000099282 | tetraspanin 15 [Source:HGNC Symbol;Acc:HGNC:23298] | *TSPAN15* | -5.02 | 0.0191 | Decreased in 3rd trimester pregnant individuals |

| | | | | | |
|---|---|---|---|---|---|
| ENSG00000172482 | alanine--glyoxylate and serine--pyruvate aminotransferase [Source:HGNC Symbol;Acc:HGNC:341] | *AGXT* | -4.99 | 0.0356 | Decreased in 3rd trimester pregnant individuals |
| ENSG00000105518 | transmembrane protein 205 [Source:HGNC Symbol;Acc:HGNC:29631] | *TMEM205* | -4.99 | 0.0111 | Decreased in 3rd trimester pregnant individuals |
| ENSG00000132823 | oxidative stress responsive serine rich 1 [Source:HGNC Symbol;Acc:HGNC:16105] | *OSER1* | -4.99 | 0.0153 | Decreased in 3rd trimester pregnant individuals |
| ENSG00000101457 | deoxynucleotidyltransferase terminal interacting protein 1 [Source:HGNC Symbol;Acc:HGNC:16160] | *DNTTIP1* | -4.96 | 0.0169 | Decreased in 3rd trimester pregnant individuals |
| ENSG00000138442 | WD repeat domain 12 [Source:HGNC Symbol;Acc:HGNC:14098] | *WDR12* | -4.94 | 0.0100 | Decreased in 3rd trimester pregnant individuals |
| ENSG00000188536 | hemoglobin subunit alpha 2 [Source:HGNC Symbol;Acc:HGNC:4824] | *HBA2* | -4.93 | 0.0092 | Decreased in 3rd trimester pregnant individuals |
| ENSG00000102178 | ubiquitin like 4A [Source:HGNC Symbol;Acc:HGNC:12505] | *UBL4A* | -4.92 | 0.0265 | Decreased in 3rd trimester pregnant individuals |
| ENSG00000129484 | poly(ADP-ribose) polymerase 2 [Source:HGNC Symbol;Acc:HGNC:272] | *PARP2* | -4.92 | 0.0100 | Decreased in 3rd trimester pregnant individuals |
| ENSG00000188488 | serpin family A member 5 [Source:HGNC Symbol;Acc:HGNC:8723] | *SERPINA5* | -4.90 | 0.0169 | Decreased in 3rd trimester pregnant individuals |
| ENSG00000011243 | A-kinase anchoring protein 8 like [Source:HGNC Symbol;Acc:HGNC:29857] | *AKAP8L* | -4.90 | 0.0122 | Decreased in 3rd trimester pregnant individuals |
| ENSG00000123130 | acyl-CoA thioesterase 9 [Source:HGNC Symbol;Acc:HGNC:17152] | *ACOT9* | -4.90 | 0.0254 | Decreased in 3rd trimester pregnant individuals |
| ENSG00000109107 | aldolase, fructose-bisphosphate C [Source:HGNC Symbol;Acc:HGNC:418] | *ALDOC* | -4.89 | 0.0145 | Decreased in 3rd trimester pregnant individuals |
| ENSG00000131966 | actin related protein 10 [Source:HGNC Symbol;Acc:HGNC:17372] | *ACTR10* | -4.87 | 0.0109 | Decreased in 3rd trimester pregnant individuals |

| | | | | | |
|---|---|---|---|---|---|
| ENSG00000108039 | X-prolyl aminopeptidase 1 [Source:HGNC Symbol;Acc:HGNC:12822] | *XPNPEP1* | -4.85 | 0.0165 | Decreased in 3rd trimester pregnant individuals |
| ENSG00000128965 | ChaC glutathione specific gamma-glutamylcyclotransferase 1 [Source:HGNC Symbol;Acc:HGNC:28680] | *CHAC1* | -4.81 | 0.0227 | Decreased in 3rd trimester pregnant individuals |
| ENSG00000234608 | MAPKAPK5 antisense RNA 1 [Source:HGNC Symbol;Acc:HGNC:24091] | *MAPKAPK5-AS1* | -4.81 | 0.0260 | Decreased in 3rd trimester pregnant individuals |
| ENSG00000162542 | transmembrane and coiled-coil domains 4 [Source:HGNC Symbol;Acc:HGNC:27393] | *TMCO4* | -4.80 | 0.0121 | Decreased in 3rd trimester pregnant individuals |
| ENSG00000181666 | zinc finger protein 875 [Source:HGNC Symbol;Acc:HGNC:4928] | *ZNF875* | -4.80 | 0.0254 | Decreased in 3rd trimester pregnant individuals |
| ENSG00000179163 | alpha-L-fucosidase 1 [Source:HGNC Symbol;Acc:HGNC:4006] | *FUCA1* | -4.76 | 0.0263 | Decreased in 3rd trimester pregnant individuals |
| ENSG00000198805 | purine nucleoside phosphorylase [Source:HGNC Symbol;Acc:HGNC:7892] | *PNP* | -4.74 | 0.0237 | Decreased in 3rd trimester pregnant individuals |
| ENSG00000175348 | TMEM9 domain family member B [Source:HGNC Symbol;Acc:HGNC:1168] | *TMEM9B* | -4.73 | 0.0169 | Decreased in 3rd trimester pregnant individuals |
| ENSG00000248144 | alcohol dehydrogenase 1C (class I), gamma polypeptide [Source:HGNC Symbol;Acc:HGNC:251] | *ADH1C* | -4.72 | 0.0203 | Decreased in 3rd trimester pregnant individuals |
| ENSG00000105655 | inositol-3-phosphate synthase 1 [Source:HGNC Symbol;Acc:HGNC:29821] | *ISYNA1* | -4.72 | 0.0260 | Decreased in 3rd trimester pregnant individuals |
| ENSG00000135241 | patatin like phospholipase domain containing 8 [Source:HGNC Symbol;Acc:HGNC:28900] | *PNPLA8* | -4.70 | 0.0260 | Decreased in 3rd trimester pregnant individuals |
| ENSG00000115825 | protein kinase D3 [Source:HGNC Symbol;Acc:HGNC:9408] | *PRKD3* | -4.70 | 0.0100 | Decreased in 3rd trimester pregnant individuals |
| ENSG00000104915 | syntaxin 10 [Source:HGNC Symbol;Acc:HGNC:11428] | *STX10* | -4.66 | 0.0263 | Decreased in 3rd trimester pregnant individuals |

| | | | | | |
|---|---|---|---|---|---|
| ENSG00000134283 | periphilin 1 [Source:HGNC Symbol;Acc:HGNC:19369] | *PPHLN1* | -4.66 | 0.0169 | Decreased in 3rd trimester pregnant individuals |
| ENSG00000053900 | anaphase promoting complex subunit 4 [Source:HGNC Symbol;Acc:HGNC:19990] | *ANAPC4* | -4.66 | 0.0383 | Decreased in 3rd trimester pregnant individuals |
| ENSG00000068903 | sirtuin 2 [Source:HGNC Symbol;Acc:HGNC:10886] | *SIRT2* | -4.65 | 0.0193 | Decreased in 3rd trimester pregnant individuals |
| ENSG00000068394 | G-patch domain and KOW motifs [Source:HGNC Symbol;Acc:HGNC:30677] | *GPKOW* | -4.65 | 0.0386 | Decreased in 3rd trimester pregnant individuals |
| ENSG00000184470 | thioredoxin reductase 2 [Source:HGNC Symbol;Acc:HGNC:18155] | *TXNRD2* | -4.65 | 0.0355 | Decreased in 3rd trimester pregnant individuals |
| ENSG00000153561 | required for meiotic nuclear division 5 homolog A [Source:HGNC Symbol;Acc:HGNC:25850] | *RMND5A* | -4.65 | 0.0165 | Decreased in 3rd trimester pregnant individuals |
| ENSG00000170854 | ribosomal oxygenase 2 [Source:HGNC Symbol;Acc:HGNC:19441] | *RIOX2* | -4.65 | 0.0263 | Decreased in 3rd trimester pregnant individuals |
| ENSG00000141258 | small G protein signaling modulator 2 [Source:HGNC Symbol;Acc:HGNC:29026] | *SGSM2* | -4.64 | 0.0196 | Decreased in 3rd trimester pregnant individuals |
| ENSG00000275700 | apoptosis antagonizing transcription factor [Source:HGNC Symbol;Acc:HGNC:19235] | *AATF* | -4.63 | 0.0100 | Decreased in 3rd trimester pregnant individuals |
| ENSG00000067836 | rogdi atypical leucine zipper [Source:HGNC Symbol;Acc:HGNC:29478] | *ROGDI* | -4.60 | 0.0263 | Decreased in 3rd trimester pregnant individuals |
| ENSG00000267106 | ZNF561 antisense RNA 1 (head to head) [Source:HGNC Symbol;Acc:HGNC:27613] | *ZNF561-AS1* | -4.60 | 0.0206 | Decreased in 3rd trimester pregnant individuals |
| ENSG00000135972 | mitochondrial ribosomal protein S9 [Source:HGNC Symbol;Acc:HGNC:14501] | *MRPS9* | -4.59 | 0.0206 | Decreased in 3rd trimester pregnant individuals |
| ENSG00000156502 | Suv3 like RNA helicase [Source:HGNC Symbol;Acc:HGNC:11471] | *SUPV3L1* | -4.59 | 0.0169 | Decreased in 3rd trimester pregnant individuals |

| | | | | | |
|---|---|---|---|---|---|
| ENSG00000132274 | tripartite motif containing 22 [Source:HGNC Symbol;Acc:HGNC:16379] | *TRIM22* | -4.59 | 0.0122 | Decreased in 3rd trimester pregnant individuals |
| ENSG00000185966 | late cornified envelope 3E [Source:HGNC Symbol;Acc:HGNC:29463] | *LCE3E* | -4.58 | 0.0190 | Decreased in 3rd trimester pregnant individuals |
| ENSG00000147155 | EBP cholestenol delta-isomerase [Source:HGNC Symbol;Acc:HGNC:3133] | *EBP* | -4.58 | 0.0260 | Decreased in 3rd trimester pregnant individuals |
| ENSG00000065268 | WD repeat domain 18 [Source:HGNC Symbol;Acc:HGNC:17956] | *WDR18* | -4.57 | 0.0347 | Decreased in 3rd trimester pregnant individuals |
| ENSG00000132581 | stromal cell derived factor 2 [Source:HGNC Symbol;Acc:HGNC:10675] | *SDF2* | -4.56 | 0.0263 | Decreased in 3rd trimester pregnant individuals |
| ENSG00000132612 | vacuolar protein sorting 4 homolog A [Source:HGNC Symbol;Acc:HGNC:13488] | *VPS4A* | -4.55 | 0.0254 | Decreased in 3rd trimester pregnant individuals |
| ENSG00000120948 | TAR DNA binding protein [Source:HGNC Symbol;Acc:HGNC:11571] | *TARDBP* | -4.55 | 0.0169 | Decreased in 3rd trimester pregnant individuals |
| ENSG00000244462 | RNA binding motif protein 12 [Source:HGNC Symbol;Acc:HGNC:9898] | *RBM12* | -4.55 | 0.0240 | Decreased in 3rd trimester pregnant individuals |
| ENSG00000033800 | protein inhibitor of activated STAT 1 [Source:HGNC Symbol;Acc:HGNC:2752] | *PIAS1* | -4.55 | 0.0301 | Decreased in 3rd trimester pregnant individuals |
| ENSG00000173545 | zinc finger protein 622 [Source:HGNC Symbol;Acc:HGNC:30958] | *ZNF622* | -4.54 | 0.0201 | Decreased in 3rd trimester pregnant individuals |
| ENSG00000173171 | metaxin 1 [Source:HGNC Symbol;Acc:HGNC:7504] | *MTX1* | -4.53 | 0.0207 | Decreased in 3rd trimester pregnant individuals |
| ENSG00000157551 | potassium inwardly rectifying channel subfamily J member 15 [Source:HGNC Symbol;Acc:HGNC:6261] | *KCNJ15* | -4.52 | 0.0323 | Decreased in 3rd trimester pregnant individuals |
| ENSG00000159111 | mitochondrial ribosomal protein L10 [Source:HGNC Symbol;Acc:HGNC:14055] | *MRPL10* | -4.52 | 0.0206 | Decreased in 3rd trimester pregnant individuals |

| | | | | | |
|---|---|---|---|---|---|
| ENSG00000148334 | prostaglandin E synthase 2 [Source:HGNC Symbol;Acc:HGNC:17822] | *PTGES2* | -4.50 | 0.0428 | Decreased in 3rd trimester pregnant individuals |
| ENSG00000260549 | metallothionein 1L, pseudogene [Source:HGNC Symbol;Acc:HGNC:7404] | *MT1L* | -4.48 | 0.0240 | Decreased in 3rd trimester pregnant individuals |
| ENSG00000197647 | zinc finger protein 433 [Source:HGNC Symbol;Acc:HGNC:20811] | *ZNF433* | -4.48 | 0.0366 | Decreased in 3rd trimester pregnant individuals |
| ENSG00000167700 | major facilitator superfamily domain containing 3 [Source:HGNC Symbol;Acc:HGNC:25157] | *MFSD3* | -4.48 | 0.0392 | Decreased in 3rd trimester pregnant individuals |
| ENSG00000101843 | proteasome 26S subunit, non-ATPase 10 [Source:HGNC Symbol;Acc:HGNC:9555] | *PSMD10* | -4.47 | 0.0263 | Decreased in 3rd trimester pregnant individuals |
| ENSG00000276023 | dual specificity phosphatase 14 [Source:HGNC Symbol;Acc:HGNC:17007] | *DUSP14* | -4.46 | 0.0323 | Decreased in 3rd trimester pregnant individuals |
| ENSG00000110442 | COMM domain containing 9 [Source:HGNC Symbol;Acc:HGNC:25014] | *COMMD9* | -4.46 | 0.0240 | Decreased in 3rd trimester pregnant individuals |
| ENSG00000034152 | mitogen-activated protein kinase kinase 3 [Source:HGNC Symbol;Acc:HGNC:6843] | *MAP2K3* | -4.46 | 0.0193 | Decreased in 3rd trimester pregnant individuals |
| ENSG00000187778 | microspherule protein 1 [Source:HGNC Symbol;Acc:HGNC:6960] | *MCRS1* | -4.45 | 0.0287 | Decreased in 3rd trimester pregnant individuals |
| ENSG00000262919 | cyclin Q [Source:HGNC Symbol;Acc:HGNC:28434] | *CCNQ* | -4.44 | 0.0136 | Decreased in 3rd trimester pregnant individuals |
| ENSG00000070961 | ATPase plasma membrane Ca2+ transporting 1 [Source:HGNC Symbol;Acc:HGNC:814] | *ATP2B1* | -4.42 | 0.0169 | Decreased in 3rd trimester pregnant individuals |
| ENSG00000100439 | abhydrolase domain containing 4, N-acyl phospholipase B [Source:HGNC Symbol;Acc:HGNC:20154] | *ABHD4* | -4.42 | 0.0287 | Decreased in 3rd trimester pregnant individuals |
| ENSG00000162496 | dehydrogenase/reductase 3 [Source:HGNC Symbol;Acc:HGNC:17693] | *DHRS3* | -4.41 | 0.0206 | Decreased in 3rd trimester pregnant individuals |

| | | | | | |
|---|---|---|---|---|---|
| ENSG00000110002 | von Willebrand factor A domain containing 5A [Source:HGNC Symbol;Acc:HGNC:6658] | *VWA5A* | -4.40 | 0.0355 | Decreased in 3rd trimester pregnant individuals |
| ENSG00000119986 | arginine vasopressin induced 1 [Source:HGNC Symbol;Acc:HGNC:30898] | *AVPI1* | -4.38 | 0.0265 | Decreased in 3rd trimester pregnant individuals |
| ENSG00000106771 | transmembrane protein 245 [Source:HGNC Symbol;Acc:HGNC:1363] | *TMEM245* | -4.37 | 0.0494 | Decreased in 3rd trimester pregnant individuals |
| ENSG00000118579 | mediator complex subunit 28 [Source:HGNC Symbol;Acc:HGNC:24628] | *MED28* | -4.37 | 0.0201 | Decreased in 3rd trimester pregnant individuals |
| ENSG00000136875 | pre-mRNA processing factor 4 [Source:HGNC Symbol;Acc:HGNC:17349] | *PRPF4* | -4.37 | 0.0277 | Decreased in 3rd trimester pregnant individuals |
| ENSG00000165819 | methyltransferase 3, N6-adenosine-methyltransferase complex catalytic subunit [Source:HGNC Symbol;Acc:HGNC:17563] | *METTL3* | -4.37 | 0.0264 | Decreased in 3rd trimester pregnant individuals |
| ENSG00000109794 | family with sequence similarity 149 member A [Source:HGNC Symbol;Acc:HGNC:24527] | *FAM149A* | -4.37 | 0.0251 | Decreased in 3rd trimester pregnant individuals |
| ENSG00000156398 | sideroflexin 2 [Source:HGNC Symbol;Acc:HGNC:16086] | *SFXN2* | -4.36 | 0.0386 | Decreased in 3rd trimester pregnant individuals |
| ENSG00000104332 | secreted frizzled related protein 1 [Source:HGNC Symbol;Acc:HGNC:10776] | *SFRP1* | -4.36 | 0.0181 | Decreased in 3rd trimester pregnant individuals |
| ENSG00000170619 | COMM domain containing 5 [Source:HGNC Symbol;Acc:HGNC:17902] | *COMMD5* | -4.35 | 0.0263 | Decreased in 3rd trimester pregnant individuals |
| ENSG00000164896 | Fas activated serine/threonine kinase [Source:HGNC Symbol;Acc:HGNC:24676] | *FASTK* | -4.35 | 0.0263 | Decreased in 3rd trimester pregnant individuals |
| ENSG00000104695 | protein phosphatase 2 catalytic subunit beta [Source:HGNC Symbol;Acc:HGNC:9300] | *PPP2CB* | -4.34 | 0.0169 | Decreased in 3rd trimester pregnant individuals |
| ENSG00000183605 | sideroflexin 4 [Source:HGNC Symbol;Acc:HGNC:16088] | *SFXN4* | -4.32 | 0.0345 | Decreased in 3rd trimester pregnant individuals |

| | | | | | |
|---|---|---|---|---|---|
| ENSG00000153094 | BCL2 like 11 [Source:HGNC Symbol;Acc:HGNC:994] | *BCL2L11* | -4.32 | 0.0169 | Decreased in 3rd trimester pregnant individuals |
| ENSG00000205250 | E2F transcription factor 4 [Source:HGNC Symbol;Acc:HGNC:3118] | *E2F4* | -4.32 | 0.0345 | Decreased in 3rd trimester pregnant individuals |
| ENSG00000127955 | G protein subunit alpha i1 [Source:HGNC Symbol;Acc:HGNC:4384] | *GNAI1* | -4.30 | 0.0301 | Decreased in 3rd trimester pregnant individuals |
| ENSG00000151657 | Kin17 DNA and RNA binding protein [Source:HGNC Symbol;Acc:HGNC:6327] | *KIN* | -4.30 | 0.0447 | Decreased in 3rd trimester pregnant individuals |
| ENSG00000176444 | CDC like kinase 2 [Source:HGNC Symbol;Acc:HGNC:2069] | *CLK2* | -4.30 | 0.0392 | Decreased in 3rd trimester pregnant individuals |
| ENSG00000185324 | cyclin dependent kinase 10 [Source:HGNC Symbol;Acc:HGNC:1770] | *CDK10* | -4.29 | 0.0467 | Decreased in 3rd trimester pregnant individuals |
| ENSG00000197165 | sulfotransferase family 1A member 2 [Source:HGNC Symbol;Acc:HGNC:11454] | *SULT1A2* | -4.29 | 0.0347 | Decreased in 3rd trimester pregnant individuals |
| ENSG00000136877 | folylpolyglutamate synthase [Source:HGNC Symbol;Acc:HGNC:3824] | *FPGS* | -4.29 | 0.0387 | Decreased in 3rd trimester pregnant individuals |
| ENSG00000092439 | transient receptor potential cation channel subfamily M member 7 [Source:HGNC Symbol;Acc:HGNC:17994] | *TRPM7* | -4.29 | 0.0165 | Decreased in 3rd trimester pregnant individuals |
| ENSG00000105472 | C-type lectin domain containing 11A [Source:HGNC Symbol;Acc:HGNC:10576] | *CLEC11A* | -4.29 | 0.0263 | Decreased in 3rd trimester pregnant individuals |
| ENSG00000109861 | cathepsin C [Source:HGNC Symbol;Acc:HGNC:2528] | *CTSC* | -4.26 | 0.0392 | Decreased in 3rd trimester pregnant individuals |
| ENSG00000134684 | tyrosyl-tRNA synthetase 1 [Source:HGNC Symbol;Acc:HGNC:12840] | *YARS1* | -4.24 | 0.0329 | Decreased in 3rd trimester pregnant individuals |
| ENSG00000125834 | serine/threonine kinase 35 [Source:HGNC Symbol;Acc:HGNC:16254] | *STK35* | -4.24 | 0.0206 | Decreased in 3rd trimester pregnant individuals |

| | | | | | |
|---|---|---|---|---|---|
| ENSG00000131437 | kinesin family member 3A [Source:HGNC Symbol;Acc:HGNC:6319] | *KIF3A* | -4.24 | 0.0337 | Decreased in 3rd trimester pregnant individuals |
| ENSG00000173039 | RELA proto-oncogene, NF-kB subunit [Source:HGNC Symbol;Acc:HGNC:9955] | *RELA* | -4.23 | 0.0111 | Decreased in 3rd trimester pregnant individuals |
| ENSG00000165943 | modulator of apoptosis 1 [Source:HGNC Symbol;Acc:HGNC:16658] | *MOAP1* | -4.23 | 0.0428 | Decreased in 3rd trimester pregnant individuals |
| ENSG00000172466 | zinc finger protein 24 [Source:HGNC Symbol;Acc:HGNC:13032] | *ZNF24* | -4.22 | 0.0254 | Decreased in 3rd trimester pregnant individuals |
| ENSG00000130147 | SH3 domain binding protein 4 [Source:HGNC Symbol;Acc:HGNC:10826] | *SH3BP4* | -4.22 | 0.0491 | Decreased in 3rd trimester pregnant individuals |
| ENSG00000188176 | smoothelin like 2 [Source:HGNC Symbol;Acc:HGNC:24764] | *SMTNL2* | -4.22 | 0.0323 | Decreased in 3rd trimester pregnant individuals |
| ENSG00000103152 | N-methylpurine DNA glycosylase [Source:HGNC Symbol;Acc:HGNC:7211] | *MPG* | -4.21 | 0.0386 | Decreased in 3rd trimester pregnant individuals |
| ENSG00000156284 | claudin 8 [Source:HGNC Symbol;Acc:HGNC:2050] | *CLDN8* | -4.21 | 0.0415 | Decreased in 3rd trimester pregnant individuals |
| ENSG00000110080 | ST3 beta-galactoside alpha-2,3-sialyltransferase 4 [Source:HGNC Symbol;Acc:HGNC:10864] | *ST3GAL4* | -4.21 | 0.0400 | Decreased in 3rd trimester pregnant individuals |
| ENSG00000151612 | zinc finger protein 827 [Source:HGNC Symbol;Acc:HGNC:27193] | *ZNF827* | -4.20 | 0.0260 | Decreased in 3rd trimester pregnant individuals |
| ENSG00000158234 | Fas apoptotic inhibitory molecule [Source:HGNC Symbol;Acc:HGNC:18703] | *FAIM* | -4.19 | 0.0323 | Decreased in 3rd trimester pregnant individuals |
| ENSG00000185591 | Sp1 transcription factor [Source:HGNC Symbol;Acc:HGNC:11205] | *SP1* | -4.17 | 0.0254 | Decreased in 3rd trimester pregnant individuals |
| ENSG00000123739 | phospholipase A2 group XIIA [Source:HGNC Symbol;Acc:HGNC:18554] | *PLA2G12A* | -4.16 | 0.0317 | Decreased in 3rd trimester pregnant individuals |

| | | | | | |
|---|---|---|---|---|---|
| ENSG00000178802 | mannose phosphate isomerase [Source:HGNC Symbol;Acc:HGNC:7216] | *MPI* | -4.16 | 0.0495 | Decreased in 3rd trimester pregnant individuals |
| ENSG00000124702 | kelch domain containing 3 [Source:HGNC Symbol;Acc:HGNC:20704] | *KLHDC3* | -4.16 | 0.0263 | Decreased in 3rd trimester pregnant individuals |
| ENSG00000109046 | WD repeat and SOCS box containing 1 [Source:HGNC Symbol;Acc:HGNC:19221] | *WSB1* | -4.15 | 0.0263 | Decreased in 3rd trimester pregnant individuals |
| ENSG00000163617 | coiled-coil domain containing 191 [Source:HGNC Symbol;Acc:HGNC:29272] | *CCDC191* | -4.14 | 0.0452 | Decreased in 3rd trimester pregnant individuals |
| ENSG00000064703 | DEAD-box helicase 20 [Source:HGNC Symbol;Acc:HGNC:2743] | *DDX20* | -4.13 | 0.0432 | Decreased in 3rd trimester pregnant individuals |
| ENSG00000115468 | EF-hand domain family member D1 [Source:HGNC Symbol;Acc:HGNC:29556] | *EFHD1* | -4.12 | 0.0181 | Decreased in 3rd trimester pregnant individuals |
| ENSG00000188895 | MSL complex subunit 1 [Source:HGNC Symbol;Acc:HGNC:27905] | *MSL1* | -4.12 | 0.0289 | Decreased in 3rd trimester pregnant individuals |
| ENSG00000143153 | ATPase Na+/K+ transporting subunit beta 1 [Source:HGNC Symbol;Acc:HGNC:804] | *ATP1B1* | -4.12 | 0.0196 | Decreased in 3rd trimester pregnant individuals |
| ENSG00000135966 | transforming growth factor beta receptor associated protein 1 [Source:HGNC Symbol;Acc:HGNC:16836] | *TGFBRAP1* | -4.11 | 0.0406 | Decreased in 3rd trimester pregnant individuals |
| ENSG00000102871 | TNFRSF1A associated via death domain [Source:HGNC Symbol;Acc:HGNC:12030] | *TRADD* | -4.11 | 0.0355 | Decreased in 3rd trimester pregnant individuals |
| ENSG00000213445 | signal-induced proliferation-associated 1 [Source:HGNC Symbol;Acc:HGNC:10885] | *SIPA1* | -4.10 | 0.0452 | Decreased in 3rd trimester pregnant individuals |
| ENSG00000163644 | protein phosphatase, Mg2+/Mn2+ dependent 1K [Source:HGNC Symbol;Acc:HGNC:25415] | *PPM1K* | -4.10 | 0.0263 | Decreased in 3rd trimester pregnant individuals |
| ENSG00000076555 | acetyl-CoA carboxylase beta [Source:HGNC Symbol;Acc:HGNC:85] | *ACACB* | -4.09 | 0.0104 | Decreased in 3rd trimester pregnant individuals |

| | | | | | |
|---|---|---|---|---|---|
| ENSG00000196712 | neurofibromin 1 [Source:HGNC Symbol;Acc:HGNC:7765] | *NF1* | -4.09 | 0.0413 | Decreased in 3rd trimester pregnant individuals |
| ENSG00000160216 | 1-acylglycerol-3-phosphate O-acyltransferase 3 [Source:HGNC Symbol;Acc:HGNC:326] | *AGPAT3* | -4.09 | 0.0317 | Decreased in 3rd trimester pregnant individuals |
| ENSG00000165688 | peptidase, mitochondrial processing subunit alpha [Source:HGNC Symbol;Acc:HGNC:18667] | *PMPCA* | -4.09 | 0.0290 | Decreased in 3rd trimester pregnant individuals |
| ENSG00000169155 | zinc finger and BTB domain containing 43 [Source:HGNC Symbol;Acc:HGNC:17908] | *ZBTB43* | -4.09 | 0.0381 | Decreased in 3rd trimester pregnant individuals |
| ENSG00000103260 | meteorin, glial cell differentiation regulator [Source:HGNC Symbol;Acc:HGNC:14151] | *METRN* | -4.08 | 0.0462 | Decreased in 3rd trimester pregnant individuals |
| ENSG00000131669 | ninjurin 1 [Source:HGNC Symbol;Acc:HGNC:7824] | *NINJ1* | -4.08 | 0.0263 | Decreased in 3rd trimester pregnant individuals |
| ENSG00000078070 | methylcrotonyl-CoA carboxylase subunit 1 [Source:HGNC Symbol;Acc:HGNC:6936] | *MCCC1* | -4.07 | 0.0366 | Decreased in 3rd trimester pregnant individuals |
| ENSG00000111371 | solute carrier family 38 member 1 [Source:HGNC Symbol;Acc:HGNC:13447] | *SLC38A1* | -4.07 | 0.0323 | Decreased in 3rd trimester pregnant individuals |
| ENSG00000112357 | peroxisomal biogenesis factor 7 [Source:HGNC Symbol;Acc:HGNC:8860] | *PEX7* | -4.06 | 0.0490 | Decreased in 3rd trimester pregnant individuals |
| ENSG00000136848 | DAB2 interacting protein [Source:HGNC Symbol;Acc:HGNC:17294] | *DAB2IP* | -4.06 | 0.0413 | Decreased in 3rd trimester pregnant individuals |
| ENSG00000086598 | transmembrane p24 trafficking protein 2 [Source:HGNC Symbol;Acc:HGNC:16996] | *TMED2* | -4.06 | 0.0315 | Decreased in 3rd trimester pregnant individuals |
| ENSG00000120833 | suppressor of cytokine signaling 2 [Source:HGNC Symbol;Acc:HGNC:19382] | *SOCS2* | -4.05 | 0.0190 | Decreased in 3rd trimester pregnant individuals |
| ENSG00000063978 | ring finger protein 4 [Source:HGNC Symbol;Acc:HGNC:10067] | *RNF4* | -4.05 | 0.0254 | Decreased in 3rd trimester pregnant individuals |

| | | | | | |
|---|---|---|---|---|---|
| ENSG00000197555 | signal induced proliferation associated 1 like 1 [Source:HGNC Symbol;Acc:HGNC:20284] | *SIPA1L1* | -4.04 | 0.0306 | Decreased in 3rd trimester pregnant individuals |
| ENSG00000084234 | amyloid beta precursor like protein 2 [Source:HGNC Symbol;Acc:HGNC:598] | *APLP2* | -4.02 | 0.0263 | Decreased in 3rd trimester pregnant individuals |
| ENSG00000154640 | BTG anti-proliferation factor 3 [Source:HGNC Symbol;Acc:HGNC:1132] | *BTG3* | -4.01 | 0.0347 | Decreased in 3rd trimester pregnant individuals |
| ENSG00000170561 | iroquois homeobox 2 [Source:HGNC Symbol;Acc:HGNC:14359] | *IRX2* | -4.00 | 0.0206 | Decreased in 3rd trimester pregnant individuals |
| ENSG00000108666 | chromosome 17 open reading frame 75 [Source:HGNC Symbol;Acc:HGNC:30173] | *C17orf75* | -3.99 | 0.0329 | Decreased in 3rd trimester pregnant individuals |
| ENSG00000197620 | endothelium and lymphocyte associated ASCH domain 1 [Source:HGNC Symbol;Acc:HGNC:28089] | *EOLA1* | -3.99 | 0.0403 | Decreased in 3rd trimester pregnant individuals |
| ENSG00000183354 | KIAA2026 [Source:HGNC Symbol;Acc:HGNC:23378] | *KIAA2026* | -3.99 | 0.0432 | Decreased in 3rd trimester pregnant individuals |
| ENSG00000187954 | cysteine and histidine rich 1 [Source:HGNC Symbol;Acc:HGNC:17806] | *CYHR1* | -3.98 | 0.0495 | Decreased in 3rd trimester pregnant individuals |
| ENSG00000101247 | NADH:ubiquinone oxidoreductase complex assembly factor 5 [Source:HGNC Symbol;Acc:HGNC:15899] | *NDUFAF5* | -3.97 | 0.0382 | Decreased in 3rd trimester pregnant individuals |
| ENSG00000165704 | hypoxanthine phosphoribosyltransferase 1 [Source:HGNC Symbol;Acc:HGNC:5157] | *HPRT1* | -3.97 | 0.0462 | Decreased in 3rd trimester pregnant individuals |
| ENSG00000140262 | transcription factor 12 [Source:HGNC Symbol;Acc:HGNC:11623] | *TCF12* | -3.96 | 0.0472 | Decreased in 3rd trimester pregnant individuals |
| ENSG00000113649 | transcription elongation regulator 1 [Source:HGNC Symbol;Acc:HGNC:15630] | *TCERG1* | -3.96 | 0.0495 | Decreased in 3rd trimester pregnant individuals |
| ENSG00000184908 | chloride voltage-gated channel Kb [Source:HGNC Symbol;Acc:HGNC:2027] | *CLCNKB* | -3.96 | 0.0323 | Decreased in 3rd trimester pregnant individuals |

| | | | | | |
|---|---|---|---|---|---|
| ENSG00000186493 | chromosome 5 open reading frame 38 [Source:HGNC Symbol;Acc:HGNC:24226] | *C5orf38* | -3.95 | 0.0366 | Decreased in 3rd trimester pregnant individuals |
| ENSG00000160087 | ubiquitin conjugating enzyme E2 J2 [Source:HGNC Symbol;Acc:HGNC:19268] | *UBE2J2* | -3.95 | 0.0386 | Decreased in 3rd trimester pregnant individuals |
| ENSG00000011275 | ring finger protein 216 [Source:HGNC Symbol;Acc:HGNC:21698] | *RNF216* | -3.94 | 0.0392 | Decreased in 3rd trimester pregnant individuals |
| ENSG00000108671 | proteasome 26S subunit, non-ATPase 11 [Source:HGNC Symbol;Acc:HGNC:9556] | *PSMD11* | -3.93 | 0.0254 | Decreased in 3rd trimester pregnant individuals |
| ENSG00000144357 | ubiquitin protein ligase E3 component n-recognin 3 [Source:HGNC Symbol;Acc:HGNC:30467] | *UBR3* | -3.92 | 0.0323 | Decreased in 3rd trimester pregnant individuals |
| ENSG00000040633 | PHD finger protein 23 [Source:HGNC Symbol;Acc:HGNC:28428] | *PHF23* | -3.91 | 0.0413 | Decreased in 3rd trimester pregnant individuals |
| ENSG00000101335 | myosin light chain 9 [Source:HGNC Symbol;Acc:HGNC:15754] | *MYL9* | -3.91 | 0.0490 | Decreased in 3rd trimester pregnant individuals |
| ENSG00000130958 | solute carrier family 35 member D2 [Source:HGNC Symbol;Acc:HGNC:20799] | *SLC35D2* | -3.91 | 0.0417 | Decreased in 3rd trimester pregnant individuals |
| ENSG00000111725 | protein kinase AMP-activated non-catalytic subunit beta 1 [Source:HGNC Symbol;Acc:HGNC:9378] | *PRKAB1* | -3.91 | 0.0406 | Decreased in 3rd trimester pregnant individuals |
| ENSG00000167799 | nudix hydrolase 8 [Source:HGNC Symbol;Acc:HGNC:8055] | *NUDT8* | -3.89 | 0.0452 | Decreased in 3rd trimester pregnant individuals |
| ENSG00000149809 | transmembrane 7 superfamily member 2 [Source:HGNC Symbol;Acc:HGNC:11863] | *TM7SF2* | -3.88 | 0.0428 | Decreased in 3rd trimester pregnant individuals |
| ENSG00000110651 | CD81 molecule [Source:HGNC Symbol;Acc:HGNC:1701] | *CD81* | -3.88 | 0.0467 | Decreased in 3rd trimester pregnant individuals |
| ENSG00000119414 | protein phosphatase 6 catalytic subunit [Source:HGNC Symbol;Acc:HGNC:9323] | *PPP6C* | -3.86 | 0.0416 | Decreased in 3rd trimester pregnant individuals |

| | | | | | |
|---|---|---|---|---|---|
| ENSG00000120137 | pantothenate kinase 3 [Source:HGNC Symbol;Acc:HGNC:19365] | *PANK3* | -3.86 | 0.0388 | Decreased in 3rd trimester pregnant individuals |
| ENSG00000145879 | serine peptidase inhibitor Kazal type 7 [Source:HGNC Symbol;Acc:HGNC:24643] | *SPINK7* | -3.86 | 0.0387 | Decreased in 3rd trimester pregnant individuals |
| ENSG00000130227 | exportin 7 [Source:HGNC Symbol;Acc:HGNC:14108] | *XPO7* | -3.85 | 0.0472 | Decreased in 3rd trimester pregnant individuals |
| ENSG00000233639 | POU3F3 adjacent non-coding transcript 1 [Source:HGNC Symbol;Acc:HGNC:49513] | *PANTR1* | -3.85 | 0.0484 | Decreased in 3rd trimester pregnant individuals |
| ENSG00000213722 | dimethylarginine dimethylaminohydrolase 2 [Source:HGNC Symbol;Acc:HGNC:2716] | *DDAH2* | -3.84 | 0.0413 | Decreased in 3rd trimester pregnant individuals |
| ENSG00000099797 | trans-2,3-enoyl-CoA reductase [Source:HGNC Symbol;Acc:HGNC:4551] | *TECR* | -3.84 | 0.0413 | Decreased in 3rd trimester pregnant individuals |
| ENSG00000064652 | sorting nexin 24 [Source:HGNC Symbol;Acc:HGNC:21533] | *SNX24* | -3.83 | 0.0377 | Decreased in 3rd trimester pregnant individuals |
| ENSG00000143621 | interleukin enhancer binding factor 2 [Source:HGNC Symbol;Acc:HGNC:6037] | *ILF2* | -3.83 | 0.0462 | Decreased in 3rd trimester pregnant individuals |
| ENSG00000188001 | tumor protein p63 regulated 1 [Source:HGNC Symbol;Acc:HGNC:24759] | *TPRG1* | -3.81 | 0.0383 | Decreased in 3rd trimester pregnant individuals |
| ENSG00000138760 | scavenger receptor class B member 2 [Source:HGNC Symbol;Acc:HGNC:1665] | *SCARB2* | -3.80 | 0.0467 | Decreased in 3rd trimester pregnant individuals |
| ENSG00000125734 | G protein-coupled receptor 108 [Source:HGNC Symbol;Acc:HGNC:17829] | *GPR108* | -3.78 | 0.0462 | Decreased in 3rd trimester pregnant individuals |
| ENSG00000247077 | PGAM family member 5, mitochondrial serine/threonine protein phosphatase [Source:HGNC Symbol;Acc:HGNC:28763] | *PGAM5* | -3.78 | 0.0388 | Decreased in 3rd trimester pregnant individuals |
| ENSG00000175879 | homeobox D8 [Source:HGNC Symbol;Acc:HGNC:5139] | *HOXD8* | -3.76 | 0.0388 | Decreased in 3rd trimester pregnant individuals |

| | | | | | |
|---|---|---|---|---|---|
| ENSG00000092203 | TOX high mobility group box family member 4 [Source:HGNC Symbol;Acc:HGNC:20161] | *TOX4* | -3.74 | 0.0404 | Decreased in 3rd trimester pregnant individuals |
| ENSG00000147649 | metadherin [Source:HGNC Symbol;Acc:HGNC:29608] | *MTDH* | -3.70 | 0.0383 | Decreased in 3rd trimester pregnant individuals |
| ENSG00000174231 | pre-mRNA processing factor 8 [Source:HGNC Symbol;Acc:HGNC:17340] | *PRPF8* | -3.69 | 0.0329 | Decreased in 3rd trimester pregnant individuals |
| ENSG00000110619 | cysteinyl-tRNA synthetase 1 [Source:HGNC Symbol;Acc:HGNC:1493] | *CARS1* | -3.67 | 0.0490 | Decreased in 3rd trimester pregnant individuals |
| ENSG00000123728 | RAP2C, member of RAS oncogene family [Source:HGNC Symbol;Acc:HGNC:21165] | *RAP2C* | -3.65 | 0.0386 | Decreased in 3rd trimester pregnant individuals |
| ENSG00000197712 | family with sequence similarity 114 member A1 [Source:HGNC Symbol;Acc:HGNC:25087] | *FAM114A1* | -3.64 | 0.0389 | Decreased in 3rd trimester pregnant individuals |
| ENSG00000198142 | sosondowah ankyrin repeat domain family member C [Source:HGNC Symbol;Acc:HGNC:26149] | *SOWAHC* | -3.64 | 0.0490 | Decreased in 3rd trimester pregnant individuals |
| ENSG00000149743 | tRNA phosphotransferase 1 [Source:HGNC Symbol;Acc:HGNC:20316] | *TRPT1* | -3.57 | 0.0415 | Decreased in 3rd trimester pregnant individuals |
| ENSG00000114573 | ATPase H+ transporting V1 subunit A [Source:HGNC Symbol;Acc:HGNC:851] | *ATP6V1A* | -3.52 | 0.0347 | Decreased in 3rd trimester pregnant individuals |
| ENSG00000148143 | zinc finger protein 462 [Source:HGNC Symbol;Acc:HGNC:21684] | *ZNF462* | -3.52 | 0.0408 | Decreased in 3rd trimester pregnant individuals |
| ENSG00000100239 | protein phosphatase 6 regulatory subunit 2 [Source:HGNC Symbol;Acc:HGNC:19253] | *PPP6R2* | -3.51 | 0.0405 | Decreased in 3rd trimester pregnant individuals |
| ENSG00000091073 | deltex E3 ubiquitin ligase 2 [Source:HGNC Symbol;Acc:HGNC:15973] | *DTX2* | -3.48 | 0.0379 | Decreased in 3rd trimester pregnant individuals |
| ENSG00000139842 | cullin 4A [Source:HGNC Symbol;Acc:HGNC:2554] | *CUL4A* | -3.37 | 0.0467 | Decreased in 3rd trimester pregnant individuals |

| | | | | | |
|---|---|---|---|---|---|
| ENSG00000019144 | pleckstrin homology like domain family B member 1 [Source:HGNC Symbol;Acc:HGNC:23697] | *PHLDB1* | -3.29 | 0.0301 | Decreased in 3rd trimester pregnant individuals |
| ENSG00000163904 | SUMO specific peptidase 2 [Source:HGNC Symbol;Acc:HGNC:23116] | *SENP2* | -3.28 | 0.0473 | Decreased in 3rd trimester pregnant individuals |
| ENSG00000179041 | ribosome biogenesis regulator 1 homolog [Source:HGNC Symbol;Acc:HGNC:17083] | *RRS1* | -3.05 | 0.0384 | Decreased in 3rd trimester pregnant individuals |
| ENSG00000147604 | ribosomal protein L7 [Source:HGNC Symbol;Acc:HGNC:10363] | *RPL7* | -1.74 | 0.0100 | Decreased in 3rd trimester pregnant individuals |
| ENSG00000122026 | ribosomal protein L21 [Source:HGNC Symbol;Acc:HGNC:10313] | *RPL21* | -1.30 | 0.0350 | Decreased in 3rd trimester pregnant individuals |
| ENSG00000147403 | ribosomal protein L10 [Source:HGNC Symbol;Acc:HGNC:10298] | *RPL10* | -1.16 | 0.0278 | Decreased in 3rd trimester pregnant individuals |
| ENSG00000174748 | ribosomal protein L15 [Source:HGNC Symbol;Acc:HGNC:10306] | *RPL15* | -1.07 | 0.0387 | Decreased in 3rd trimester pregnant individuals |
| ENSG00000154518 | ATP synthase membrane subunit c locus 3 [Source:HGNC Symbol;Acc:HGNC:843] | *ATP5MC3* | -1.05 | 0.0393 | Decreased in 3rd trimester pregnant individuals |
| ENSG00000174444 | ribosomal protein L4 [Source:HGNC Symbol;Acc:HGNC:10353] | *RPL4* | -1.04 | 0.0169 | Decreased in 3rd trimester pregnant individuals |
| ENSG00000111716 | lactate dehydrogenase B [Source:HGNC Symbol;Acc:HGNC:6541] | *LDHB* | -0.96 | 0.0388 | Decreased in 3rd trimester pregnant individuals |
| ENSG00000265972 | thioredoxin interacting protein [Source:HGNC Symbol;Acc:HGNC:16952] | *TXNIP* | -0.92 | 0.0479 | Decreased in 3rd trimester pregnant individuals |
| ENSG00000142168 | superoxide dismutase 1 [Source:HGNC Symbol;Acc:HGNC:11179] | *SOD1* | -0.72 | 0.0388 | Decreased in 3rd trimester pregnant individuals |
| ENSG00000044459 | centlein [Source:HGNC Symbol;Acc:HGNC:23432] | *CNTLN* | 4.49 | 0.0383 | Increased in 3rd trimester pregnant individuals |

| | | | | | |
|---|---|---|---|---|---|
| ENSG00000258039 | novel transcript, antisense & intronic to ANKS1B | *ENSG00000258039* | 5.95 | 0.0100 | Increased in 3rd trimester pregnant individuals |
| ENSG00000135898 | G protein-coupled receptor 55 [Source:HGNC Symbol;Acc:HGNC:4511] | *GPR55* | 6.03 | 0.0100 | Increased in 3rd trimester pregnant individuals |
| ENSG00000256663 | ubiquitin-like with PHD and ring finger domains 1 (UHRF1) pseudogene | *ENSG00000256663* | 6.35 | 0.0072 | Increased in 3rd trimester pregnant individuals |
| ENSG00000271173 | SPRING1 pseudogene 1 [Source:HGNC Symbol;Acc:HGNC:44986] | *SPRING1P1* | 7.01 | 0.0100 | Increased in 3rd trimester pregnant individuals |
