## Supplementary vFC methods for "Miniaturized Workflow for Transcriptomic Profiling of Urinary Extracellular RNA during Pregnancy"

### Supporting Methods Figures

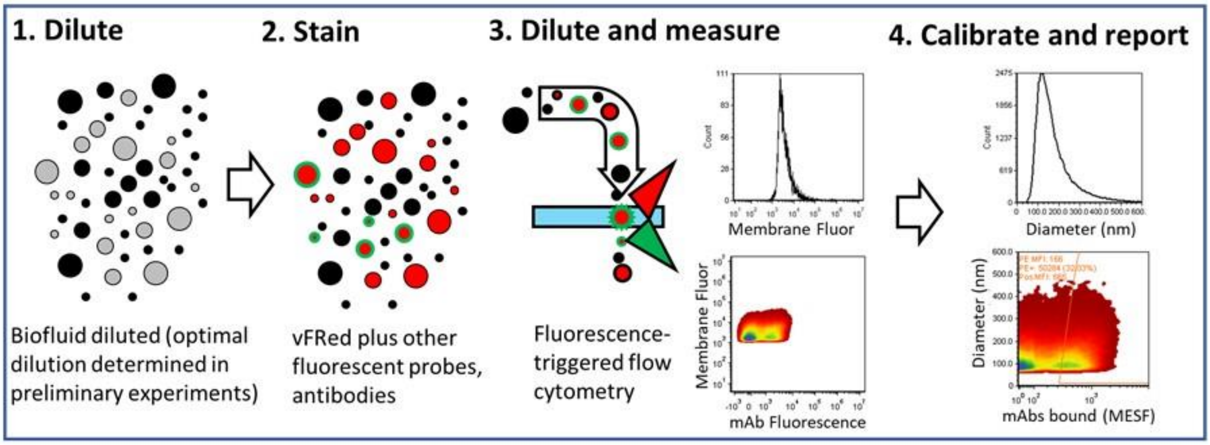

Figure SM1. Schematic of vFC workflow.

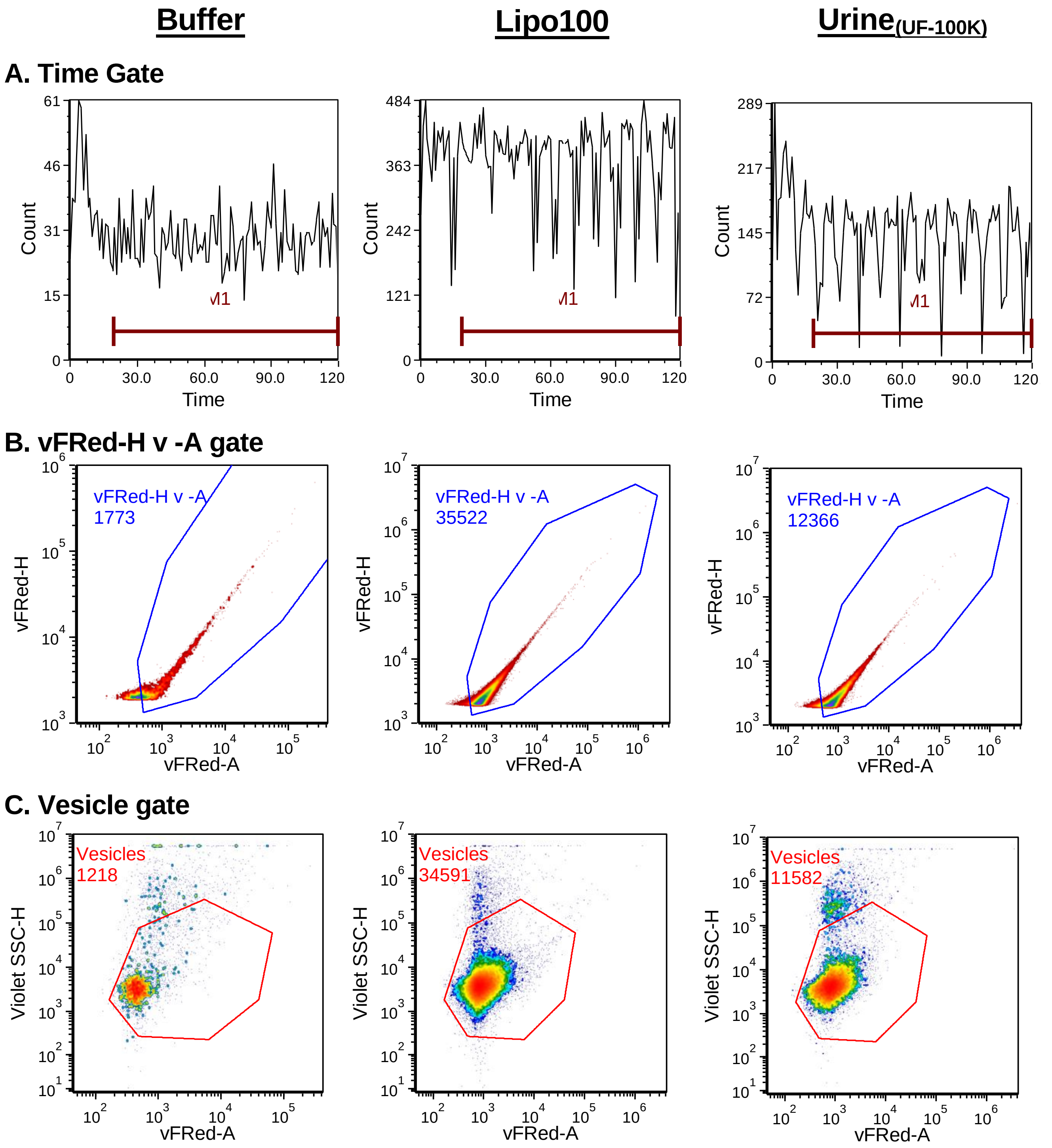

**Figure SM2. vFC gating.**

#### A. Gating

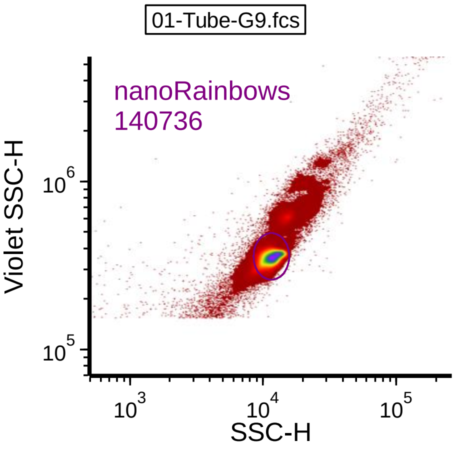

#### B. Uncalibrated vFRed Fluorescence

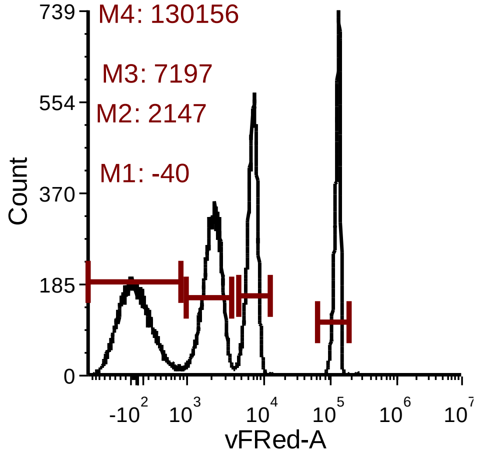

#### C. Regression

| | A | B | C | D | E | F |
|---|---|---|---|---|---|---|
| 1 | SA (nm2) | vFRed MFI | log SA | log MFI | | |
| 2 | 0 | -40 | | | | |
| 3 | 44922 | 2147 | 4.65 | 3.33 | | |
| 4 | 163832 | 7197 | 5.21 | 3.86 | | |
| 5 | 2833170 | 130156 | 6.45 | 5.11 | | |
| 6 | | | | | | |

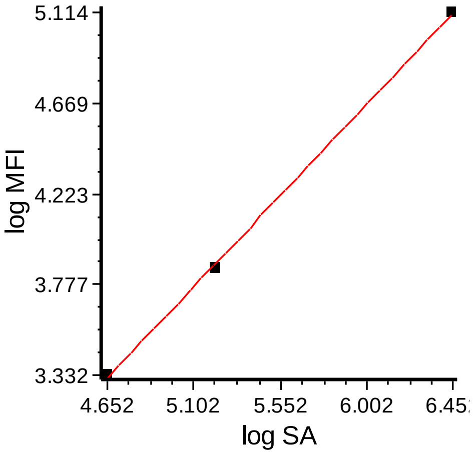

| Model name | m | b | r2 |
|---|---|---|---|
| Linear | 0.99 | -1.31 | 1.00 |

#### D. Calibrated vFRed Fluorescence

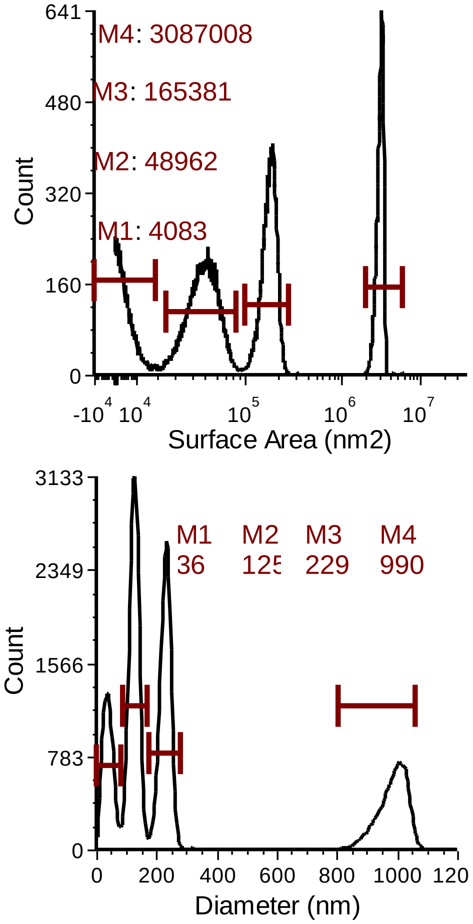

Figure SM3. Vesicle size calibration.

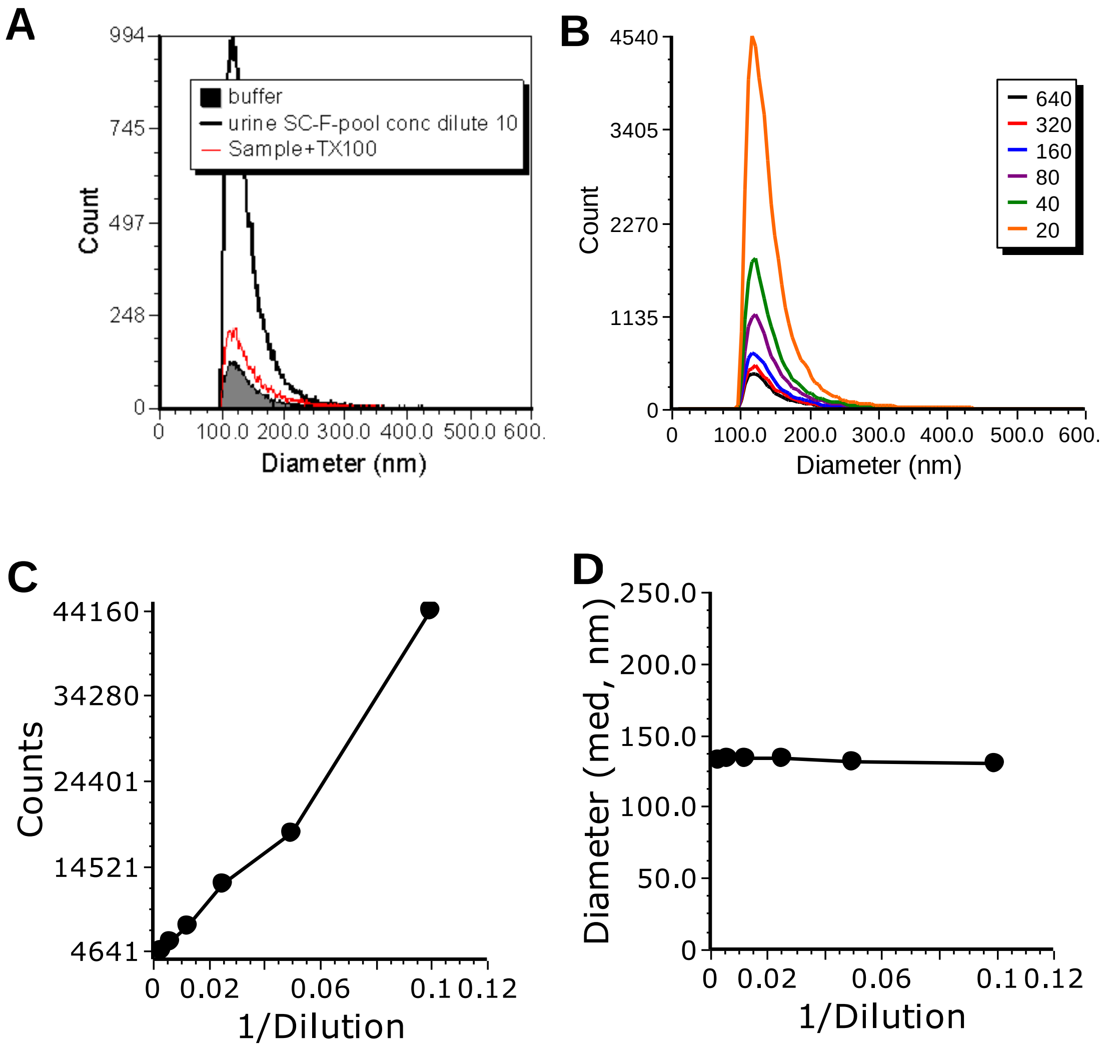

Figure SM4. Controls for single vesicle specificity.

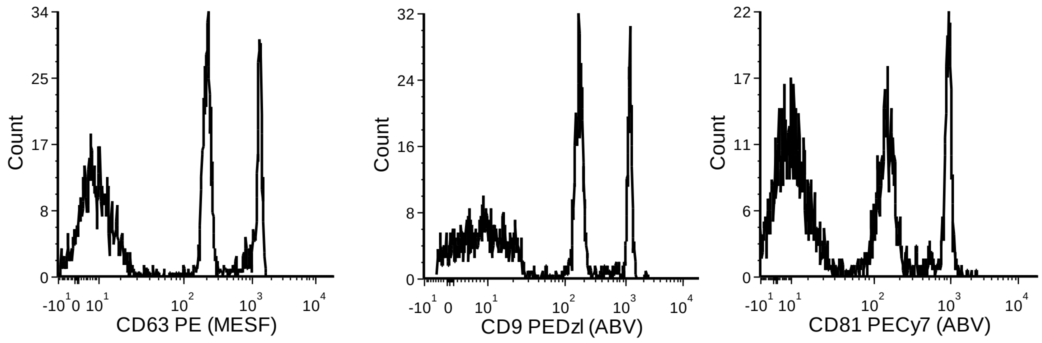

Figure SM5. Immunofluorescence controls.

**A. Non-pregnant pool (UF-100K)**

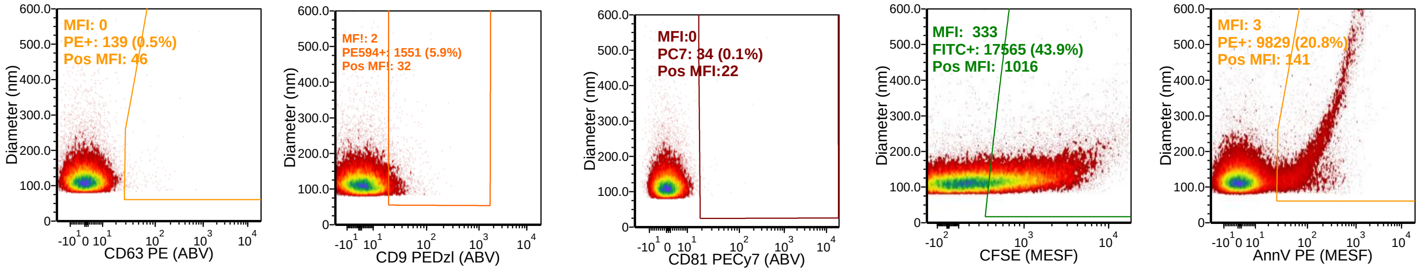

**B. Pregnant pool (UF-100K)**

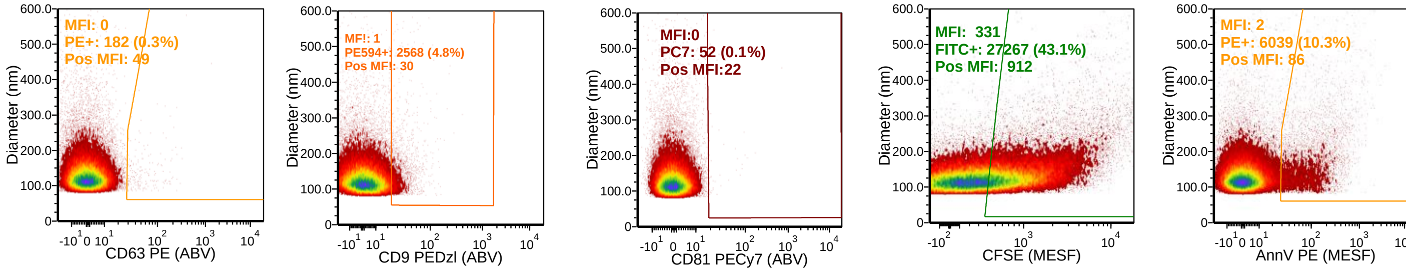

**C. Non-pregnant pool (UF-100K-ft)**

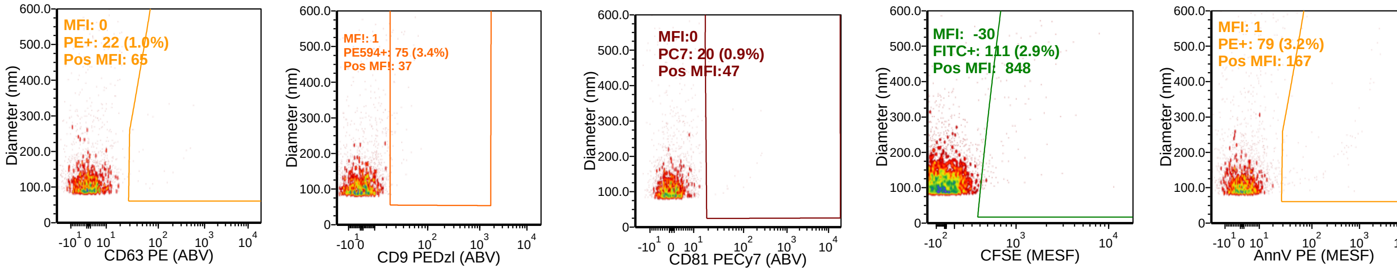

**D. Pregnant pool (UF-100K-ft)**

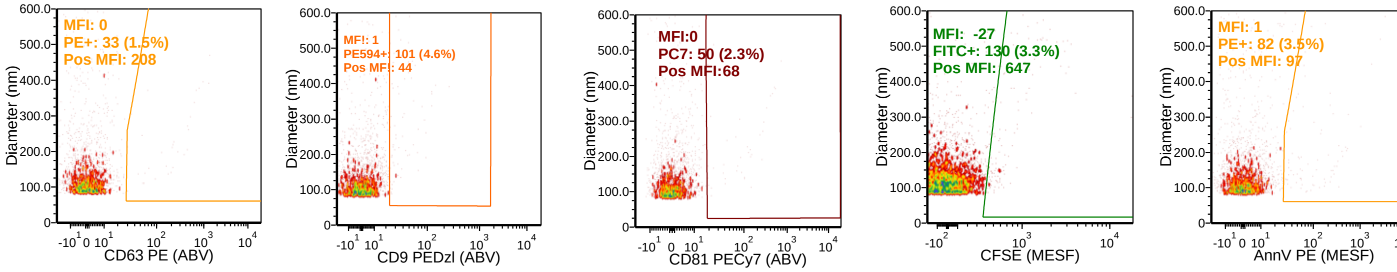

Figure SM6. vFC of neat and UF urine pools

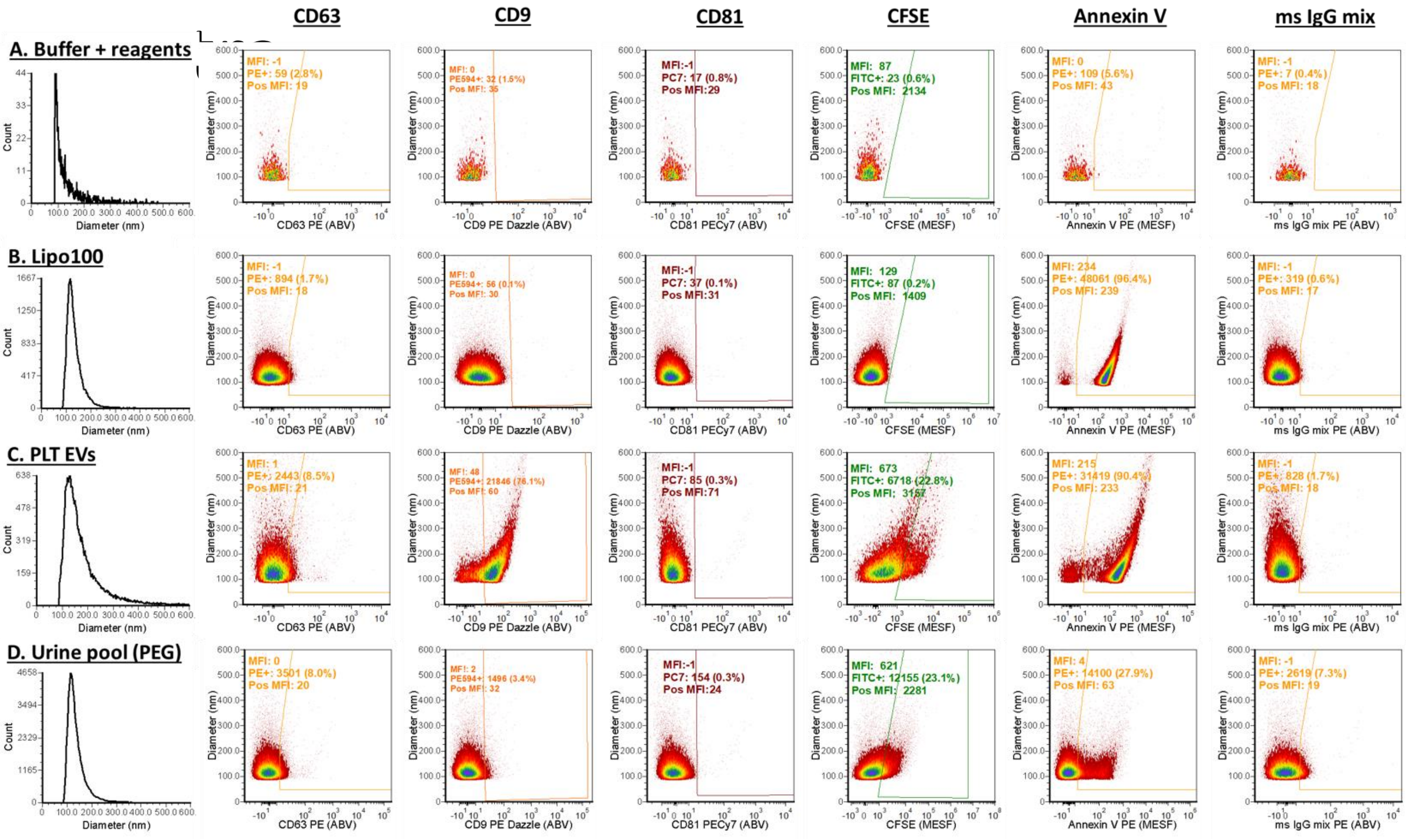

CD63
CD9
CD81
CFSE
Annexin V
ms IgG mix
A. Buffer + reagents
B. Lipo100
C. PLT EVs
D. Urine pool (PEG)
Count
Diameter (nm)
CD63 PE (ABV)
CD9 PE Dazzle (ABV)
CD81 PECy7 (ABV)
CFSE (MESF)
Annexin V PE (MESF)
ms IgG mix PE (ABV)

| Requirement | Please Include Requested Information | |
|---|---|---|
| 1.1. Purpose | To characterize EVs present in urine. | |
| 1.2. Keywords | | |
| 1.3. Experiment variables | | |
| 1.4. Organization name and address | | |
| 1.5. Primary contact name and email address | | |
| 1.6. Date or time period of experiment | | |
| 1.7. Conclusions | | |
| 1.8. Quality control measures | Instrument performance was characterized using a combination of multi-intensity single fluorophore beads (Quantum FITC, Bangs Labs Quantibrite PE, BD Biosciences) whose intensity had been calibrated in units of MESF, multi-intensity multifluorophore beads (vCal nanoRainbow, Cellarcus), and antibody capture beads (vCal nanoCal antibody capture beads, Cellarcus) calibrated to report results in units of antibodies bound per vesicle (ABV). EV analysis by vesicle flow cytometry (VFC) was conducted and reported as suggested by the MIFlowCyt-EV guidelines (See attached checklist). | |
| 2.1.1.1. (2.1.2.1., 2.1.3.1.) Sample description | | |
| 2.1.1.2. Biological sample source description | | |
| 2.1.1.3. Biological sample source organism description | | |
| 2.1.2.2. Environmental sample location | | |
| 2.3. Sample treatment description | | |
| 2.4. Fluorescence reagent(s) description | Table S1, Supporting Information | |
| 3.1. Instrument manufacturer | Beckman Coulter | Beckman C |
| 3.2. Instrument model | CytoFlex | CytoFlex |
| [illegible] | | The CytoFl<br>Detector<br>FSC<br>V-1<br>V-2<br>V-3<br>V-4<br>V-5<br>V-6<br>R-1<br>R-2<br>R-3 |

| | | |
|---|---|---|
| 3.3. Instrument configuration and settings | | B-1<br>B-2<br>B-3<br>Y-1<br>Y-2<br>Y-3<br>Y-4<br>Y-5 |
| 4.1. List-mode data files | *We recommend all authors to submit their data files to http://flowrepository.org and to make them available for the peer-review process. If you have done so, please let us know by inserting the following codes (replace the red text):<br><br>1) The link for peer-review process: http://flowrepository.org/id/<br><br>This link will only be shared with reviewers of your manuscript. | |
| 4.2. Compensation description | Single color spectral reference samples were prepared using antibody capture beads (nanoComp or nanoCal, Cellarcus), and run to determine spectra/spectral spillover of each for spectral unmixing/compensation. | |
| 4.3. Data transformation details | | |

| | |
|---|---|
| 4.4.1. Gate description | Data were analyzed using a standardized data analysis layout created in FCS Express version 7 (De Novo Software). Data were subjected to a Time gate (Figure S2A), to remove data associated with any fluidic anomalies (eg clogs or surges), vFRed Object Area (Figure S2B) to remove events recorded objects imaged over more than four pixels, which would be inconsistent with single EVs). These events were further gated (Vesicle gate, Figure S2C) to include events with membrane fluorescence in the blue-excited, red-emission channels of vesicles and to exclude low intensity low intensity background events. Presented are data from buffer (+vFRed, left), Lipo100 Vesicle Size Standard (center), and a VLP preparation (right). The vFRed fluorescence of Vesicle-gated events (Figure S2D) was used to estimate vesicle size and events counts in the Vesicle gate in 3.66 ul were used to estimate the EV concentrations in the original sample, after accounting for the pre-stain and post-stain dilutions. |
| 4.4.2. Gate statistics | EV immunofluorescence data was analyzed to calculate the median fluorescence intensity (MFI) of the entire EV population, the number of EVs with immunofluorescence positive above gate position at the upper threshold of an unstained sample (<0.5% “positive”), and the (MFI) of these positive EVs. |
| 4.4.3. Gate boundaries | EV immunofluorescence data was analyzed to calculate the median fluorescence intensity of the entire EV population, the number of EVs with immunofluorescence positive above gate position at the upper threshold of an unstained sample (<0.5% “positive”), and the (MFI) of these positive EVs. |

:oulter

ex flow cytometer with stock filters (see table below) was co

| Parameter Name ($PnN) | EX-EM/BP | Stain Name ($PnS) |
| --- | --- | --- |
| FSC | 488- | FSC |
| | 405-780/60 | |
| Violet SSC | 405-405/10 | VSSC |
| FL6 | 405-450/45 | V450 |
| FL7 | 405-525/40 | V525 |
| FL8 | 405-610/20 | V610 |
| | 405-660/20 | V660 |
| FL3 | 640-660/20 | APC |
| FL4 | 640-712/25 | APC700 |
| FL5 | 640-780/60 | APC780 |

| | | |
|---|---|---|
| SSC | 488-488/8 | SSC |
| FL1 | 488-525/40 | FITC |
| FL2 | 488-690/50 | vFRed |
| | 561-561/10 | |
| FL9 | 561-610/20 | PE610 |
| FL10 | 561-585/42 | PE |
| FL11 | 561-690/50 | PE690 |
| FL12 | 561-780/60 | PE780 |

| Framework Criteria | What to report | Please complete each criterion |
|---|---|---|
| 1.1 Preanalytical variables conforming to MISEV guidelines. | Preanalytical variables relating to EV sample including source, collection, isolation, storage, and any others relevant and available in the performed study. | See Methods. |
| 1.2 Experimental design according to MIFlowCyt guidelines. | EV-FC manuscripts should provide a brief description of the experimental aim, keywords, and variables for the performed FC experiment(s) using MIFlowCyt checklist criteria: 1.1, 1.2, and 1.3, respectively. Template found at www.evflowcytometry.org. | See Introduction, Supporting Info. |
| 2.1 Sample staining details | State any steps relating to the staining of samples. Along with the method used for staining, provide relevant reagent descriptions as listed in MIFlowCyt guidelines (Section 2.4 Fluorescence Reagent(s) Descriptions). | Sample staining was performed as directed in the vFC Protocol. A fresh 10x vFRed PLUS working solution was prepared from the 100x stock in Vesicle Staining Buffer. Staining reactions consisted of 5 ul diluted sample, 5 ul vFRed 10x Working Solution, and 5 uL 10x antibody added to a total volume of 50 uL in Vesicle Staining Buffer. Samples were incubated for 1 hour at ambient temperature. Following staining, sample was diluted 200-fold for analysis. |
| 2.2 Sample washing details | State any steps relating to the washing of samples. | No washing was performed. |
| 2.3 Sample dilution details | All methods and steps relating to sample dilution. | Samples were subjected to a pre-stain dilution (10-80-fold, as determined in preliminary experiments), and a 1000-fold post stain dilution prior to measurement. |
| 3.1 Buffer alone controls. | State whether a buffer-only control was analyzed at the same settings and during the same experiment as the samples of interest. If utilized it is recommended that all samples be recorded for a consistent set period of time e.g. 5 minutes, rather than stopping analysis at a set recorded event count e.g. 100,000 events. This allows comparisons of total particle counts between controls and samples. | Buffer-only controls showed fewer than 200 Vesicle-gated events per 3.66 uL analyzed. |
| 3.2 Buffer with reagent controls. | State whether a buffer with reagent control was analyzed at the same settings, same concentrations, and during the same experiment as the samples of interest. If used state what the results were. | Buffer plus reagent controls showed fewer than 1000 Vesicle-gated events per 3.66 uL analyzed. (Figure S2) |
| 3.3 Unstained controls. | State whether unstained control samples were analyzed at the same settings and during the same experiment as stained samples. If used, state what the results were, preferably in standard units. | Unstained controls were similar to Buffer-only controls and showed fewer than 2000 events per 3.66 uL analyzed. (Not shown) |
| 3.4 Isotype controls. | The use of isotype controls is applicable to immunofluorescence labelling only. State whether isotype controls were analyzed at the same settings and during the same experiment as stained samples. If utilized, state which antibody they are matched to, the concentration used, and what the results were (Section 4.2, 4.3, 4.4). Due to conjugation differences between manufacturers if should be stated if the isotype controls are from the same manufacturer as the matched antibodies. | A lack of detectable Fc Receptor mediated binding was assessed on a subset of samples using an irrelvent IgG1 at a concentration of 5 nM. The median fluorescence of the isotype-stained sample was not significantly above the background of unstained sample, indicating undetectable Fc Receptor binding. Figure S9. |
| 3.5 Single-stained controls. | State whether single-stained controls were included. If used state whether the single-stained controls were recorded using the same settings, dilutions, and during the same experiment as stained samples and state what the results were, preferably in standard units (Section 4.2, 4.3, 4.4). | Single stained controls were analyzed as part of optimization and validation of multicolor assays. |
| 3.6 Procedural controls. | State whether procedural controls were included. If used, state the procedure and if the procedural controls were acquired at the same settings and during the same experiment as stained samples. | No “procedural” controls were identified. |
| 3.7 Serial dilutions. | State whether serial dilutions were performed on samples and note the dilution range and manner of testing. The fluorescence and/or scatter signal intensity would ideally be reported in standard units (see Section 4.3, 4.4) but arbitrary units can also be used. This data is best reported by plotting the recorded number events/concentration over a set period of time at different sample dilution. The median fluorescence intensity at each of the dilutions should also ideally be plotted on the same or a separate plot. | In preliminary experiment, serial dilutions of samples were performed on selected samples to determine an optimal sample pre-stain dilution, which showed proportional decrease in events counts with minimal change in median fluorescence over a greater than >50-fold range (from ~ 1:10 – 1:640. (Figure S4) |
| 3.8. Detergent treated EV-samples | State whether samples were detergent treated to assess lability. If utilized, state what detergent was used, the end concentration of the detergent, and what the results were of the lysis. | Detergent lability was assessed on a sub-set of samples by treatment of stained sample with 0.1% Triton X100 prior to post-stain dilution and analysis. Greater than 90% of the marker-positive (antibody) events were eliminated by detergent treatment. (Figure S4) |
| 4.1 Trigger Channel(s) and Threshold(s). | The trigger channel(s) and threshold(s) used for event detection. Preferably, the fluorescence calibration (Section 4.3) and/or scatter calibration (Section 4.4) should be used in order to report the trigger channel(s) and threshold(s) in standardized units. | Data acquisition was triggered by fluorescence in the vFred channel (488-690/50) corresponding ot a diameter of ~95 nm). |
| 4.2 Flow Rate / Volumetric quantification. | State if the flow rate was quantified/validated and if so, report the result and how they were obtained. | The sample volumetric flow rate was calibrated using counting beads (Cellarcus, nanoRainbow), and was found to be within 10% of the instrument spec. |

| | | |
|---|---|---|
| 4.3 Fluorescence Calibration | State whether fluorescence calibration was implemented, and if so, report the materials and methods used, catalogue numbers, lot numbers, and supplied reference units for the standards. Fluorescence parameters may be reported in standardized units of MESF, ERF, or ABC beads. The type of regression used, and the resulting scatter plot of arbitrary data vs standard data for the reference particles should be supplied. | Fluorescence response was calibrated in units of ABV (antibodies bound per vesciel) using hard-dyed nanoRainbow calibration particles (Cellarcus Biosciences, #CBS6) that had been cross-calibrated against antibody capture beads (nanoCal, Cellarcus #CBS7-MS) and PE (QuantiBrite PE beads, BD Biosciences #340495) and MESF standards (Quantum FITC, Bangs labs on the same instrument. |
| 4.4 Light Scatter Calibration | State whether and how light scatter calibration was implemented. Light scatter parameters may be reported in standardized units of nm2, along with information required to reproduce the model. | No light scatter calibration was performed. |
| 5.1 EV diameter/surface area/volume approximation | State whether and how EV diameter, surface area, and/or volume has been calculated using FC measurements. | EV size was estimated from the vFRed$^{TM}$ intensity using the linear relationship between the population surface area and vFRed fluorescence distributions. Synthetic lipid vesicles with a uniform size distribution (estimated by NTA) and a lipid composition similar to a mammalian cell plasma membrane (Lipo100$^{TM}$, Cellarcus Biosciences, # CBS-1), were stained with vFRed$^{TM}$ and measured by flow cytometry. The Lipo100$^{TM}$ population diameter distribution was calculated from the NTA diameter distribution assuming a spherical geometry, and linear regression performed against the vFRed$^{TM}$ fluorescence distribution to determine the F/nm2, which was then used to estimate the size distribution of unknowns. Fig S3 |
| 5.2 EV refractive index approximation | State whether the EV refractive index has been approximated and how this was done. | No EV refractive index approximation was performed. |
| 5.3 EV epitope number approximation | State whether EV epitope number has been approximated, and if so, how it was approximated. | Immunofluorescence intensities are presented in units of ABV (antibodies bound per vesicle), which might be considered to represent the epitope abundance to within a factor of 2, given the bivalent nature of the IgGs used. |
| 6.1 Completion of MIFlowCyt checklist | Complete MIFlowCyt checklist criteria 1 to 4 using the MIFlowCyt guidelines. Template found at www.evflowcytometry.org. | Included in Supporting Information. |
| 6.2 Calibrated channel detection range | If fluorescence or scatter calibration has been carried out, authors should state whether the upper and lower limits of a calibrated detection channel were calculated in standardized units. This can be done by converting the arbitrary unit scale to a calibrated scaled, as discussed in Section 4.3 and 4.4, and providing the highest unit on this scale and the lowest detectable unit above the unstained population. The lowest unit at which a population is deemed 'positive' can be determined a variety of ways, including reporting the 99th percentile measurement unit of the unstained population for fluorescence. The chosen method for determining at what unit an event was deemed positive should be clearly outlined. | In general, fluorescence channels had a calibrated range of ~0-10,000 MESF/ABVs, with limits of detection (LOD, background +3SD) ranging from 10-200 MESF/ABVs, depending on the channel/fluorochrome. |
| 6.3 EV number/concentration. | State whether EV number/concentration has been reported. If calculated, it is preferable to report EV number/concentration in a standardized manner, stating the number/concentration between a set detection range. | EV concentrations are reported in EVs/mL, which is calculated from the number of events detected in the Vesicles gate and accounting for the volume analyzed and all pre- and post-stain dilutions. Marker-positive events we calculated from the number of events exceeding an arbitrary gate set at the ~99.5 percentile of the negative population. |
| 6.4 EV brightness. | When applicable, state the method by which the brightness of EVs is reported in standardized units of scatter and/or fluorescence. | EV brightness is reported in MESF/ABV units where possible and appropriate. (Figure S5) |
| 7.1. Sharing of data to a public repository. | Provide a link to the experimental data in a public data repository. | Data have been uploaded to the ISAC Flow Repository. |

#### SUPPORTING METHODS INFORMATION

##### Vesicle Flow Cytometry

Single vesicle flow cytometry (vFC$^{TM}$, **Figure SM1**) was performed using a commercial assay kit (vFC$^{TM}$ EV Analysis Kit, Cellarcus Biosciences) and flow cytometers (CytoFlexS, Beckman Coulter). Samples were diluted (optimal dilution determined in a preliminary experiments), stained with a fluorogenic membrane stain (vFRed$^{TM}$, Cellarcus Biosciences), and one of several selected surface markers (e.g. fluorescence-labeled antibodies, **Table SM1**) in a total volume of 50 uL in a 96-well v-bottom plate (Sarstedt 82.1583.001) for 1 hour at ambient temperature, according to the protocol instructions. The optimal concentrations of antibody and other reagents was determined by the manufacturer via titration and provided at 10x the final staining concentration, which was generally between 2 and 10 nM, depending on the antibody. Stained samples were diluted 1000-fold in Vesicle Staining Buffer and analyzed on the flow cytometer.

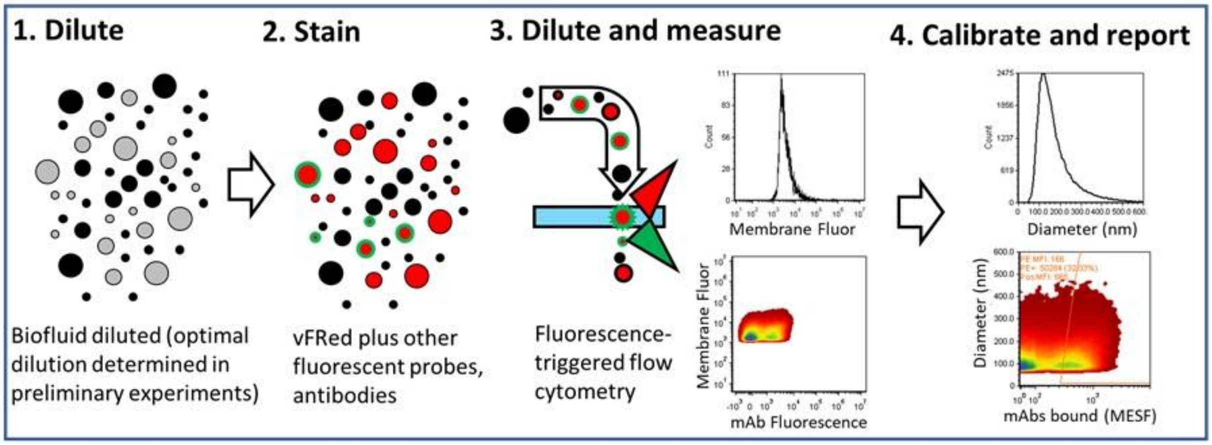

**Figure SM1. Schematic of VFC workflow.** Vesicle flow cytometry (vFC$^{TM}$) is a homogeneous assay in which a cell free sample, prepared by centrifugation, is stained with a fluorogenic membrane stain and one or more additional fluorescence probes then analyzed by flow cytometry with detection triggered by membrane fluorescence. The size distribution of a synthetic vesicle standard (Lipo100$^{TM}$), determined by NTA, is used to calibrate membrane fluorescence in terms of vesicle surface area, while fluorescence intensity and antibody capture standards are used to calibrate fluorescence intensity and antibody binding in units of MESF (mean equivalent soluble fluorochromes) or ABV (antibodies bound per vesicle).

**Table SM1. Reagents used**

| Reagent | Source | Cat number |
|---|---|---|
| vFC$^{TM}$ EV Analysis Assay kit | Cellarcus Biosciences | CBS4 |
| Anti-CD9 (HI9a) PE | Cellarcus Biosciences | CBS10-PE-100T |
| Anti-CD63 (H5C6) PE | Cellarcus Biosciences | CBS11-PE-100T |
| Anti-CD81 (5A6) PE | Cellarcus Biosciences | CBS12-PE-100T |
| Mouse IgG PE mix | Cellarcus Biosciences | CBS29-PE-100T |
| Annexin V PE | Cellarcus Biosciences | CBS34-PE-100T |
| vCal$^{TM}$ nanoRainbow beads | Cellarcus Biosciences | CBS6 |
| vCal$^{TM}$ nanoCal™ Ab Capture beads | Cellarcus Biosciences | CBS7 |

**CytoFlex configuration and sample analysis.**

The CytoFlex flow cytometer used the standard filters (MIFlowCyt Checklist) configured to measure violet side scatter (VSSC) as described in the CytoFLEX Instructions for Use (https://www.beckman.com/techdocs/B49006AP/wsr-168786). Briefly, the Violet 405nm filter is placed in position 2, the Violet 450nm filter in position 3, and an unused filter in position 1. The complete detector configuration with filters and gains is presented in Table SM2a. The instrument fluorescence response was calibrated in units of MESF in the PE (561-585/42) and FITC channels (488-525/40) using hard-dyed calibration beads (nanoRainbow calibration particles, Cellarcus) that had been cross-calibrated against PE (PE QuantiBrite beads, BD Biosciences), FITC (Quantum FITC MESF beads, Bangs Labs) MESF standards and antibody capture beads (units of Antibodies Bound per Vesicle, ABV; vCal™ AbCap, Cellarcus #CBS7-MS) on the same instrument.

Sample was introduced at the HIGH sample flow rate (60 μL/sec, confirmed using vCal™ nanoRainbow beads) and data acquisition was triggered by the 488 nm-excited red fluorescence of the vFRed membrane stain (488-690/50 channel), with a threshold set to accept ~2 events/second with a buffer-only sample, and ~20 events/sec for a buffer +vFRed sample. Samples were analyzed for 120 seconds.

**Table SM2a. CytoFlexS configuration**

| Detector | Parameter | Stain Name | Gain |
|---|---|---|---|
| V-1 | 405-780/60 | | 1000 |
| V-2 | 405-405/10 | VSSC | 500 |
| V-3 | 405-450/45 | | 1000 |
| V-4 | 405-525/40 | | 1000 |
| V-5 | 405-610/20 | | 1000 |
| V-6 | 405-660/20 | | 1000 |
| R-1 | 640-660/20 | AF647 | 1000 |
| R-2 | 640-712/25 | | 1000 |
| R-3 | 640-780/60 | | 1000 |
| B-1 | 488-488/8 | SSC | 100 |
| B-2 | 488-525/40 | | 1000 |
| B-3 | 488-690/50 | vFRed | 1000 |
| Y-1 | 561-561/10 | | 1000 |
| Y-2 | 561-610/20 | PE610 | 1000 |
| Y-3 | 561-585/42 | PE | 1000 |
| Y-4 | 561-690/50 | | 1000 |
| Y-5 | 561-780/60 | PECy7 | 1000 |

**Vesicle Gating and Data Analysis - CytoFlex.** Data were analyzed using a standardized data analysis layout in FCS Express version 7 (De Novo Software). For vFC data collected on the CytoFlexThe first 20 seconds of data were discarded via a Time gate (**Figure SM2A**) due to a consistent but unexplained background event anomaly observed on several different CytoFlex instruments. The remaining 100 seconds of data, corresponding to 100 μL of measured sample, and a plot of vFRed-A vs vFRed-H used to set a gate (**Figure SM2B**) excluding certain background events that could be identified by their lower signal pulse area and widths. These events were further gated (Vesicle gate) to include events with membrane fluorescence and light scatter intensity characteristic of EVs, and to exclude high light scatter intensity background events that have been noted in certain samples (**Figure SM2C**). Events counts in the Vesicle

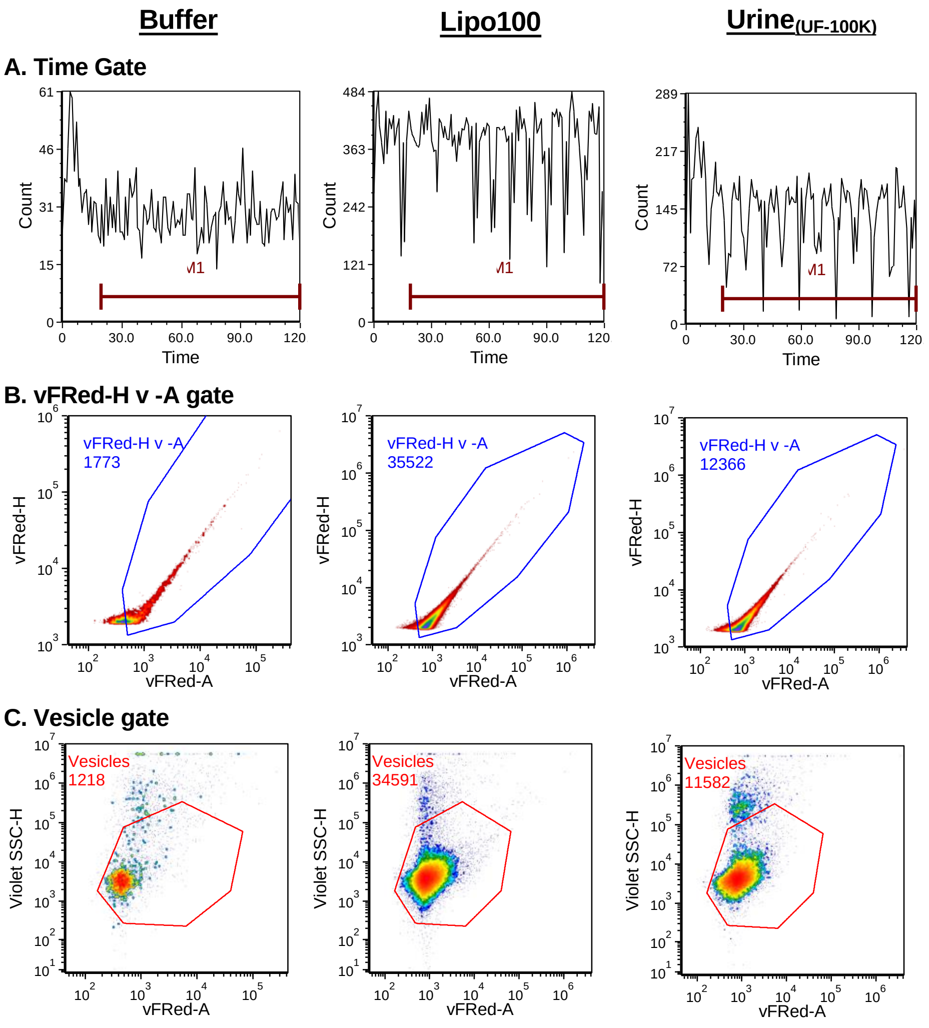

**Figure SM2. vFC gating.** vFC data from the Cytoflex is gated on time (A, to eliminate a fluidics-related background that occurs at the beginning of each sample measured in the CytoFlex), Membrane Stain fluorescence pulse height vs area (B, vFRed-H v- A, which can help resolve EV-associated events from some sources of background), and by a Vesicles Gate (C,) on vFRed-A vs VSSC-H, which selects events with characteristic EV fluorescence and light scatter.

gate in 100 μl analyzed were used to estimate EV concentrations after accounting for the pre-stain and post-stain dilutions.

**Vesicle size calibration.** Vesicle size was estimated using nanoRainbow beads that had been cross calibrated using a vesicle size standard (Lipo100, Cellarcus) whose size distribution was measured by nanoparticle tracking analysis (NTA, NanoSight, Malvern). The nanoRainbow beads were measured under the same conditions as samples (**Figure SM3B**), and the relationship between vFRed membrane fluorescence MFI mean equivalent surface area

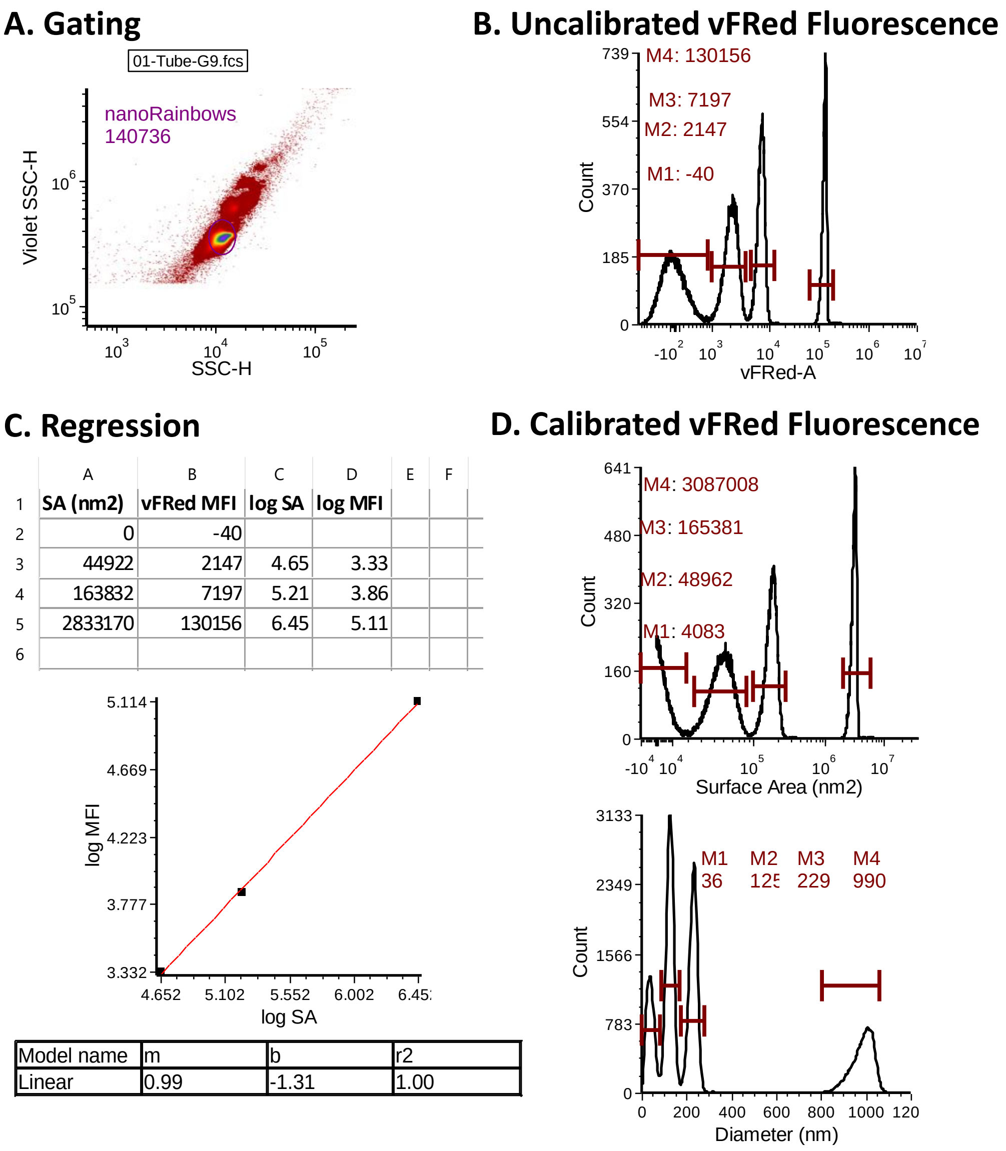

| | A | B | C | D | E | F |
|---|---|---|---|---|---|---|
| 1 | SA (nm2) | vFRed MFI | log SA | log MFI | | |
| 2 | 0 | -40 | | | | |
| 3 | 44922 | 2147 | 4.65 | 3.33 | | |
| 4 | 163832 | 7197 | 5.21 | 3.86 | | |
| 5 | 2833170 | 130156 | 6.45 | 5.11 | | |
| 6 | | | | | | |

| Model name | m | b | r2 |
|---|---|---|---|
| Linear | 0.99 | -1.31 | 1.00 |

**Figure SM3. Vesicle size estimation.** Vesicle size was estimated after calibration of vFRed Fluorescence using cross-calibrated nanoRainbow beads with assigned units of MESA (mean equivalent surface area, Cellarcus). A. Single nanoRainbow beads were gated and (B) the median fluorescence intensity fluorescence intensity of each peak was determined. (C) Linear regression of the log-transformed MESA and MFI values produce the slope (units of log SA (nm2)/log Fluorescence) and intercept, which wash used to estimate (D) the equivalent Surface Area and Diameter axes.

(MESA) was determined was characterized by linear regression after log transformation (**Figure SM3C**) to calculate the fluorescence intensity per unit surface area (F/$nm^2$). This factor was used to calibrate the membrane fluorescence axis in units of surface area and diameter (assuming spherical particles, (**Figure S3D**). The vesicle size detection limit for this instrument, estimated from the calibrated trigger threshold surface area and diameter detection, is ~95 nm.

**Controls for single vesicle analysis.** The specificity of single vesicle analysis was evaluated via several control measurements. Buffer-plus vFRed$^{TM}$ showed low levels of background events (~2000 in ~100 seconds of gated data; **Figure SM4A**). Serial dilution of sample showed the expected proportional decrease in detected events with minimal change in the brightness of those events (**Figure SM4B-C**), consistent with the measurement of single EVs.

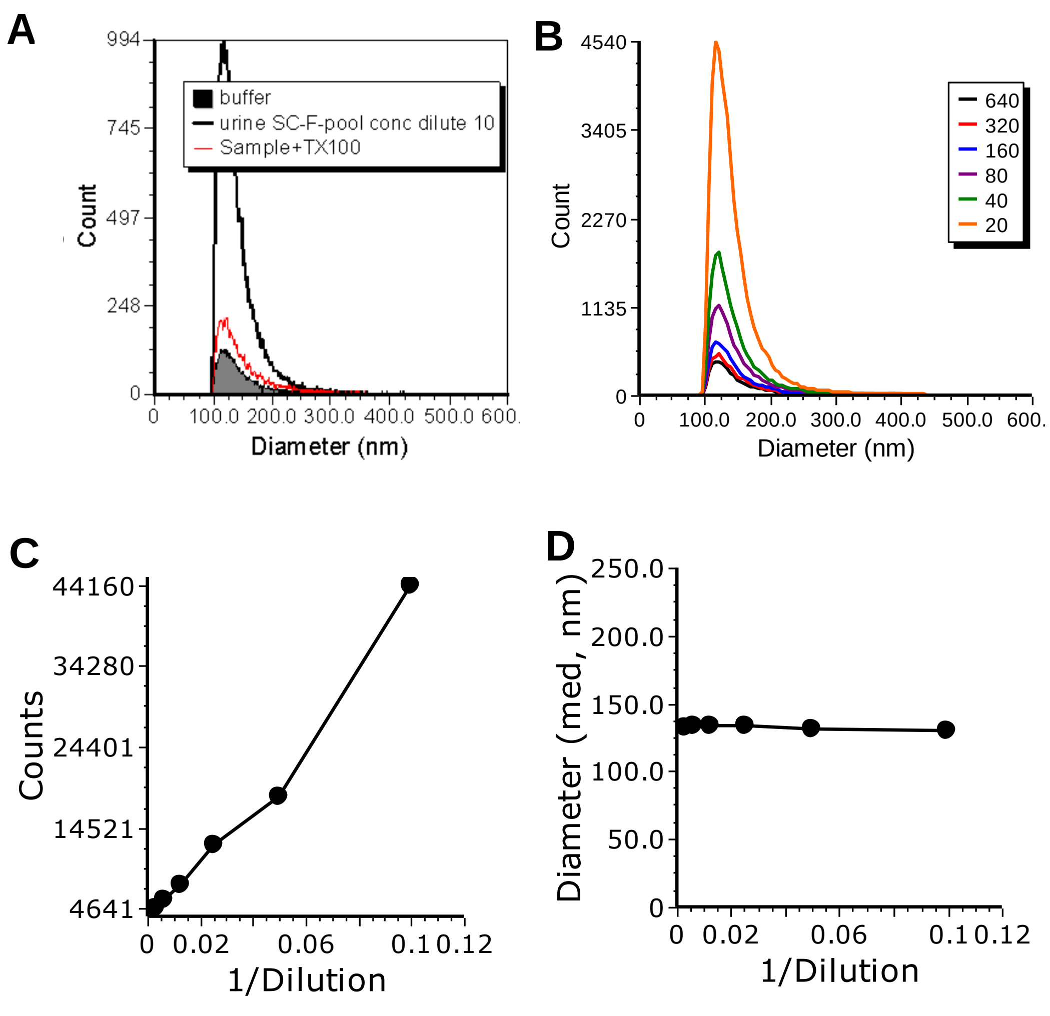

**Figure SM4. vFC Buffer +reagent and dilution controls.** The single EV specificity of the measured marker positive events (A) was demonstrated by comparison to a Buffer +vFRed (no vesicle) sample and by serial dilution (B) which demonstrated a ~2 log dynamic range (C) with no detectable coincidence/swarm.

**Fluorescence calibration.** EV immunofluorescence was calibrated in MESF (mean equivalent soluble fluorochromes) units for FITC and PE conjugates and units of ABV (antibodies bound per vesicle) using nanoRainbow beads and calibrated antibody capture beads (nanoCal, Cellarcus) that had been stained with the fluorescent antibody conjugates of interest.

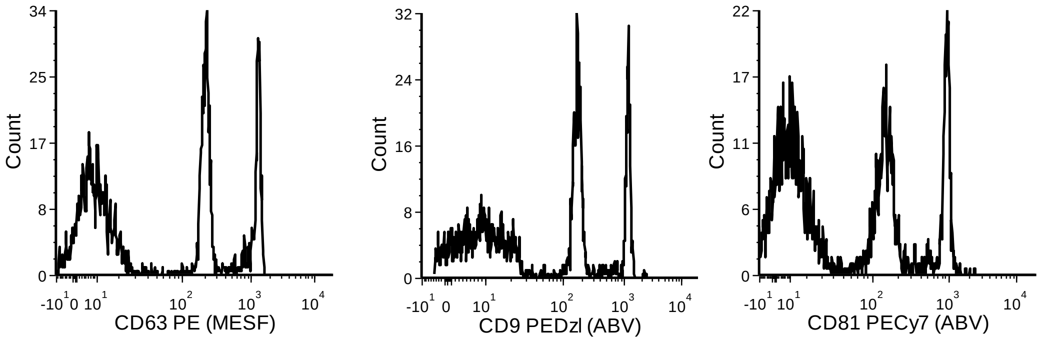

**Figure SM5. Calibrated antibody capture beads** were stained with saturating concentrations of the indicated antibodies, washed, and measured.

**Vesicle immunofluorescence controls and reporting.** Immunofluorescence negative controls included the Lipo100 vesicle standard, which bears no antigen, and a panel of IgG1, IgG2a and IgG2b PE conjugate isotype controls to detect antigen-independent binding to Fc receptors expressed on EVs (**Figure SM6**). Where available, EVs known to express the antigen of interest were used as positive controls.

EV immunofluorescence data was analyzed to calculate the median fluorescence intensity of the entire EV population, the number of EVs with immunofluorescence positive above gate position at the upper threshold of an unstained sample (<0.5% “positive”), and the (MFI) of these positive EVs.

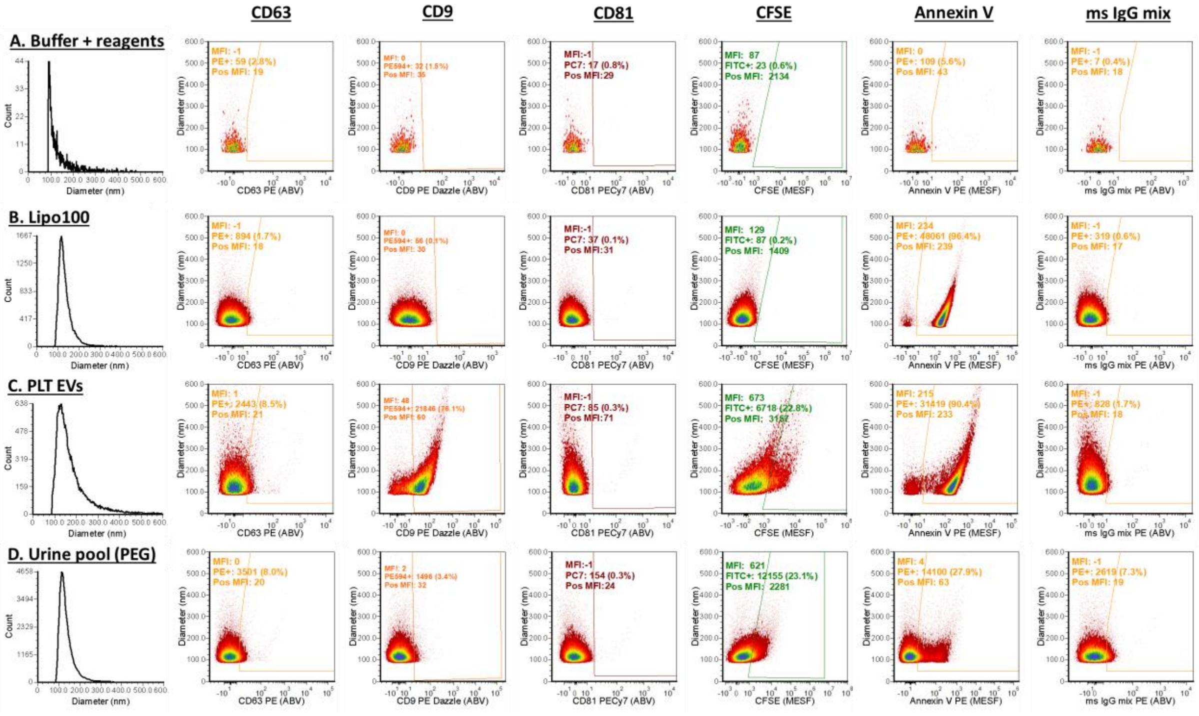

**Figure SM6. Negligible Fc Receptor-mediated antibody binding as measured using an isotype-matched irrelevant control antibody.** Representative EV samples and controls stained with the indicated probes and antibodies and a mouse IgG PE mix.

A. Non-pregnant pool (UF-100K)

B. Pregnant pool (UF-100K)

C. Non-pregnant pool (UF-100K-ft)

D. Pregnant pool (UF-100K-ft)

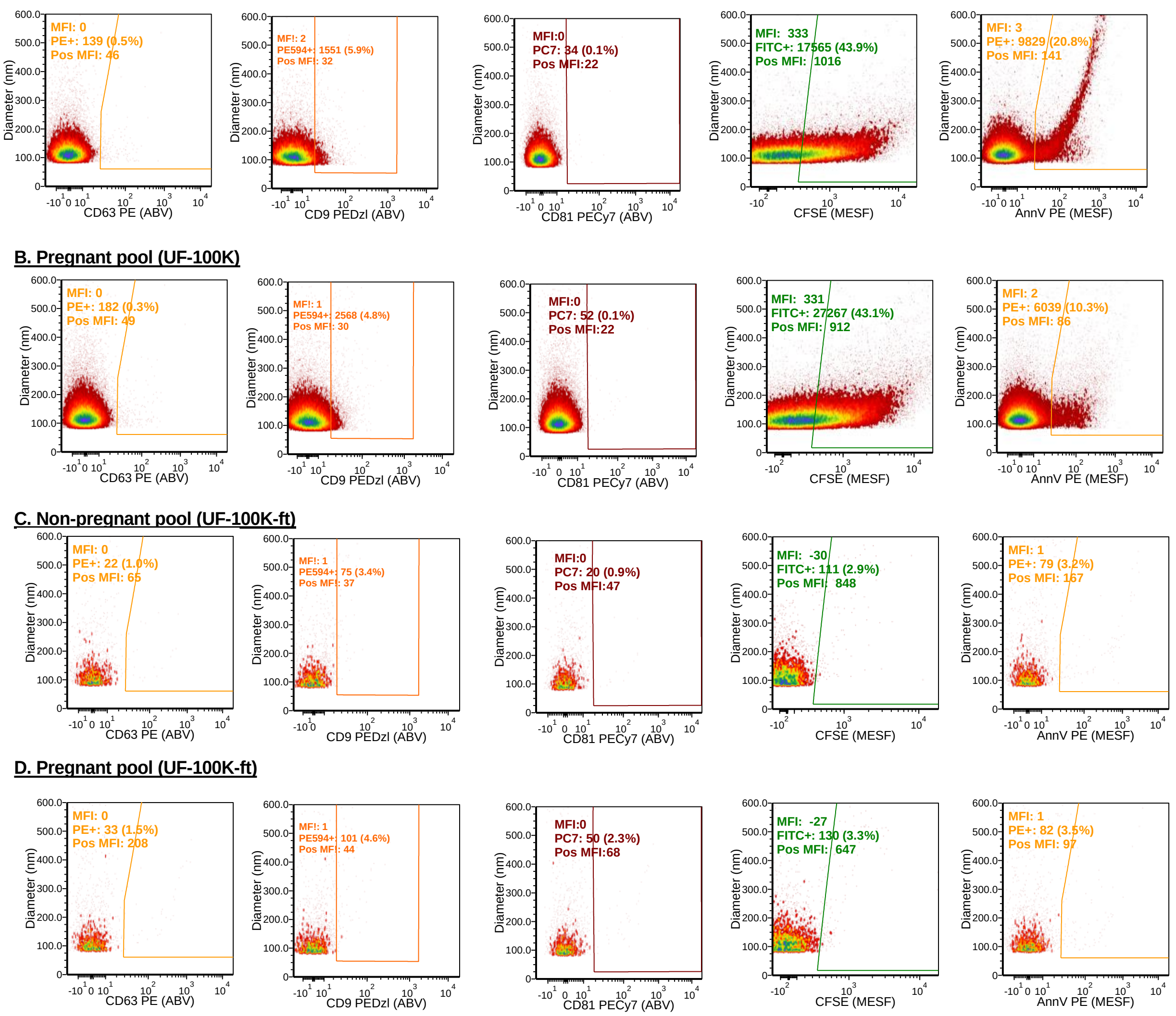

**Figure SM7. vFC measurement of Tetraspaninin expression and CFSE staining of neat and UF-100K fractionated urine pools.**
